# Dopamine modulates striatal activity when controlling the output of working memory

**DOI:** 10.64898/2026.02.01.703151

**Authors:** Natalie M. Nielsen, Rebecca D. Calcott, Ruben van den Bosch, Helena S. Olraun, Floortje S. Spronkers, Robbert-Jan Verkes, Hanneke E. M. den Ouden, Roshan Cools

**Affiliations:** Donders Institute, Nijmegen, the Netherlands; Radboud University Medical Centre, Nijmegen, the Netherlands; Radboud University, Nijmegen, the Netherlands; Donders Center for Cognition, Nijmegen, the Netherlands

**Keywords:** working memory gating, dopamine, fronto-striatal circuit, striatum

## Abstract

Cognitive control depends on capacity-limited mechanisms that require selection. Working memory (WM) therefore relies on an input-gate to filter information into WM and an output-gate to transform current WM representations into a prioritised state to guide behaviour. According to current theory, such WM gating processes are modulated by dopamine in the striatum, yet empirical evidence for this hypothesis remains sparse. We fill this gap by directly testing the role of dopamine for both WM input-and output-gating in a double-blind, placebo-controlled within-subject fMRI study (*N* = 34; 23 female, 11 male human subjects), using a 400 mg oral dose of the dopamine D2 receptor antagonist sulpiride. In our cued delayed-response task, which separated input-from output-gating, sulpiride altered neural activity in the striatum during output-gating. This effect was accompanied by a sulpiride-induced decrease in the neural stimulus selectivity of gated sensory representations in visual association cortex, with greater decreases in stimulus selectivity correlating with reduced behavioural accuracy in the task. Together, the results establish a role for striatal dopamine during the control of cortical outflow of WM.

**Significance Statement:** To act on a goal, the brain holds information in mind and brings the important parts into focus to guide what we do next. Scientists believe that this “selection” process involves dopamine, a chemical messenger, in a brain area called the striatum. We tested this by giving people a drug that alters dopamine activity in the brain. The drug changed how the striatum reacted when people selected information from memory. It also made the brain’s visual areas represent this information less clearly, and this change affected people’s behaviour. Therefore, we conclude that dopamine in the striatum plays a role when our brain controls which information in working memory is sent to other parts of the brain to guide actions.

## Introduction

Working memory (WM) is capacity-limited (Cowan, 2005). Effective goal-directed behaviour therefore requires control mechanisms to prevent interference and cognitive overload. Contemporary theories propose that dopamine is critical for these processes, regulating gating in cortico-striatal circuits by controlling both the in-and outflow of WM (Badre and Frank, 2012; Frank and Badre, 2012; Hazy et al., 2006; O’Reilly and Frank, 2006). These models distinguish an input-gate governing access of sensory information to WM and an output-gate determining which representations influence downstream processing. Despite theoretical prominence, causal evidence that dopamine controls WM input-and output-gating remains limited. Here, we fill this gap by testing whether pharmacological dopamine D2 receptor modulation with sulpiride affects visual WM gating and associated neural activity, measured with fMRI, using a task separating WM input-and output-gating.

A substantial body of work has established a critical role for prefrontal dopamine in WM. Seminal studies showed that dopamine depletion in the dorsolateral prefrontal cortex (dlPFC) of nonhuman primates impairs performance on delayed response tasks, comparable to dlPFC lesions (Brozoski et al., 1979). Subsequent neurophysiological studies further substantiated that dopamine D1 receptor stimulation modulates delay-period activity in PFC (Sawaguchi and Goldman-Rakic, 1991; Williams and Goldman-Rakic, 1995), while human pharmacological imaging studies showed dopaminergic modulation of distractor resistance during WM maintenance in dlPFC (Bloemendaal et al., 2015; Gibbs and D’Esposito, 2005). These findings support theories in which prefrontal dopamine sharpens current goal representations by enhancing the signal-to-noise ratio (Cohen and Servan-Schreiber, 1992; Durstewitz et al., 1999; Seamans and Yang, 2004; Vijayraghavan et al., 2017) and biases competitive processing in posterior sensory regions in a top-down manner (Desimone and Duncan, 1995; Gazzaley et al., 2005; Miller and Cohen, 2001; Noudoost and Moore, 2011).

However, prefrontal dopamine mechanisms alone cannot fully explain rapid gating of WM based on changing behavioural relevance (Badre, 2012). Phasic dopamine projections from the ventral tegmental area to PFC have been proposed to gate contents of WM in PFC (Cohen et al., 2002; Braver et al., 1999; D’Ardenne et al., 2012), but the slow dynamics of PFC dopamine may be incompatible with fast gating operations (Holloway et al., 2019). In contrast, striatal dopamine is ideally positioned to control rapid WM gating via cortico-striatal loops. Indeed, modern gating models extend classical basal ganglia theories of motor action selection (Gurney et al., 2001; Mink, 1996) to rapid cognitive control mechanisms (e.g., prefrontal basal ganglia working memory (PBWM) model; Hazy et al., 2007; Frank and O’Reilly, 2006; Frank et al., 2001). Cognitive gating operations build on motor mechanisms, with dopaminergic modulation of Go and NoGo pathways enhancing relevant cortical representations and suppressing irrelevant ones, thereby facilitating input-and output-gating (Kessler, 2017; Rac-Lubashevsky and Frank, 2021; Kessler and Rozanis, 2023; Chatham et al., 2014).

In keeping with this observation, accumulating evidence from (pharmacological) fMRI and PET imaging studies supports a key role for striatal activity (Trutti et al., 2024; Chatham et al., 2014; Dahlin et al., 2008; Nir-Cohen et al., 2020), and dopamine in flexible WM updating and/or input-gating (Mehta et al., 2004; Bloemendaal et al., 2015; Cools et al., 2007; Braun et al., 2021; Papenberg et al., 2020; McNab et al., 2009). For example, pharmacological studies with the selective dopamine D2 receptor antagonist sulpiride showed impaired WM updating (Mehta et al., 1999), however, there is no empirical evidence for a key role of dopamine in output-gating of WM specifically.

The present study goes beyond this previous work by explicitly dissociating effects of dopamine on distinct input-and output-gating processes. To this end, healthy participants received 400 mg of sulpiride and performed a cued delayed response task with pre-and retro-cues, enabling separate assessment of input-and output-gating. We predicted that sulpiride, given its modulatory effect on the D2 receptor-mediated NoGo pathway, would modulate gating-related enhancement of currently relevant information in stimulus-specific visual areas, accompanied by corresponding modulation of striatal activity during both input-and output-gating (preregistration: https://aspredicted.org/NQW_BVP).

## Methods

### Participants

Thirty-eight healthy volunteers were recruited via Radboud University electronic database online platform (SONA) and flyers at Radboud University, the Netherlands. Participants were English-or Dutch-speaking, right-handed and MRI-compatible. Prior to inclusion, each participant was screened extensively for exclusion criteria, including adverse psychological and physical conditions like chronical illnesses, psychiatric or neurological disorders or cardiovascular diseases (full list of exclusion criteria: Supplementary Methods). The study was approved by the regional research ethics committee (“Medisch-Ethische Toetsingscommissie Oost-Nederland”; 2020-7199; case number NL76159.091.21). Participants gave written consent and were compensated monetarily (146-166 euros) for their participation. One participant dropped out due to personal reasons and one participant had to be excluded from the analysis due to a technical failure of the MRI scanner. Moreover, two participants were excluded from the analysis due to overall task performance below 50% (including missed trials, nonresponses, and incorrect button presses), indicating insufficient task compliance. Hence, data from *N* = 34 participants were used in the present analysis (age at inclusion: 19-30; mean(SD): 22.4(2.4) years; 23 women).

### General procedure

The present placebo-controlled, cross-over pharmacological functional magnetic resonance imaging (fMRI) study employed a within-subject, double-blind design. Prior to the first visit, participants underwent a phone screening in which mental and physical health were evaluated based on a list of yes/no questions (Supplementary Methods). When admitted to the study, participants completed three sessions at the Donders Centre for Cognitive Neuroimaging in Nijmegen, the Netherlands (Figure 1).

**Figure 1.**
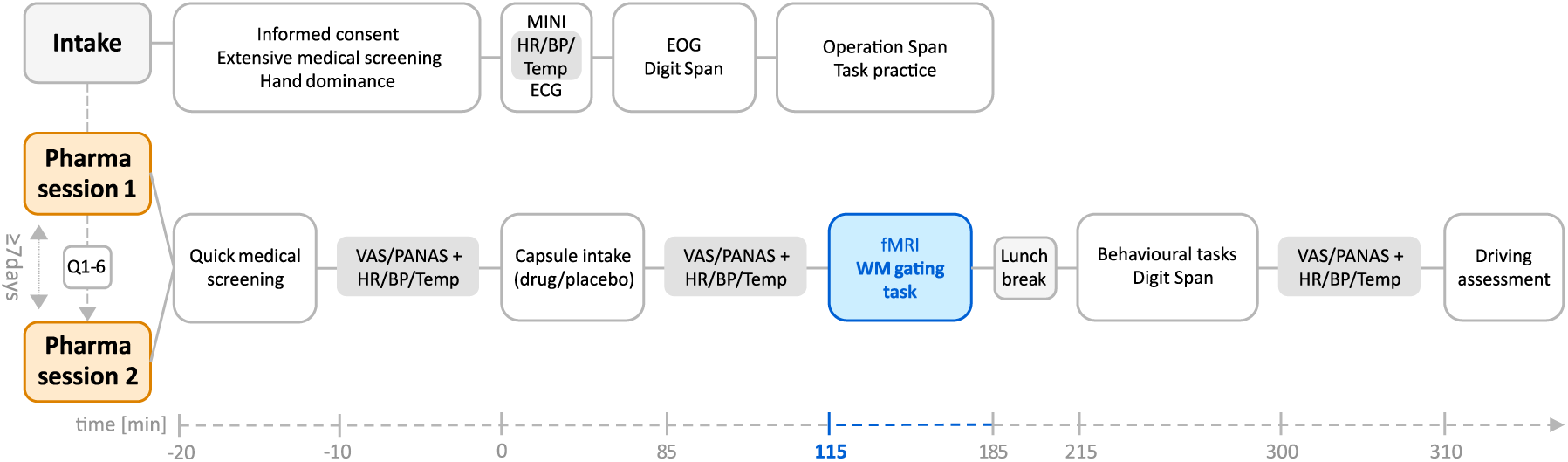
Study overview. The study included three sessions: intake, pharmacological session 1, and pharmacological session 2. The pharmacological sessions 1 and 2 were separated by at least seven days. In between these two test sessions, participants completed six questionnaires. The present study focusses on the WM gating task administered in the MRI scanner, highlighted in blue. Effects on the behavioural tasks are reported elsewhere (Spronkers et al., in preparation). MINI, Mini-International Neuropsychiatric Interview; HR, heart rate; BP, blood pressure; Temp, temperature; ECG, electrocardiogram; EOG, electrooculogram; Q1-6, questionnaires; VAS, visual analogue scale in rating subjective feelings; PANAS, positive and negative affect scale.

#### Intake session

At their first visit, participants were screened carefully within a three-hour period. This screening procedure included a medical screening (Supplementary Methods), a hand dominance question-naire (Supplementary Methods), the Mini-International Neuropsychiatric Interview (MINI for DSM-IV; Overbeek et al., 1999; Sheehan et al., 1998), an electrocardiogram (ECG) and electrooculogram (EOG) recording, as well as measurements of body temperature, heart rate and blood pressure. Additionally, participants completed two WM capacity tasks (i.e., forward and backward Digit Span task; Wechsler, 1997; Operation Span task; Turner and Engle, 1989) and practised shortened versions of all tasks of interest. Subsequently, a medical doctor decided on each individual’s eligibility to participate in the study based on the medical assessment (Supplementary Questionnaires).

#### Pharmacological sessions

The following two sessions were conducted exactly in the same way and scheduled at least one week apart to mitigate practice effects and to ensure complete clearance of the medication from the body. In between the two sessions, participants completed six short online questionnaires about their general well-being at home, enabling us to characterise the sample (Supplementary Table S4). These included questions about depressive symptoms (BDI; Beck et al., 2011), impulsivity (BIS-11; Barratt, 1994), behavioural activation and inhibition (BIS/BAS; Carver and White, 1994), state and trait anxiety (STAI; Ploeg, 1980; Ploeg et al., 1981), burnout symptoms (MBI; Maslach et al., 1997) and COVID-related stress and anxiety (CSS; Taylor et al., 2020).

The sessions started with a short medical screening (Supplementary Methods) and all women underwent a pregnancy test (hCG Zwangerschapstest; AccuBioTech). Furthermore, participants were required to abstain from cannabis for two weeks prior to each test session, from psychiatric and recreational drugs for 72 hours, and from alcohol for 24 hours before the session. After 20 minutes, participants received either a placebo or an active drug (400 mg sulpiride) through oral intake. Sulpiride is a selective dopamine D2-receptor antagonist (Caley and Weber, 1995), predominantly acting on the striatum where D2/3 receptors are abundant (Hall et al., 1994). The drug is commonly used clinically to treat depression (Rüther et al., 1999) and schizophrenia (Wagstaff et al., 1994), while it has also been repeatedly shown to alter cognitive function (Mehta, 2003; Naef et al., 2017; Van Den Bosch et al., 2022; Dodds et al., 2009; Mehta et al., 2004; Mehta et al., 2008; Westbrook et al., 2020). An independent researcher randomised the drug order and ensured counterbalancing between subjects and sessions.

Following participant exclusion, equal group sizes were maintained (*N* = 17 drug first; *N* = 17 placebo first). Physical and psychological measurements were taken three times throughout the day to keep track of subjects’ mood and medical state, including the Positive and Negative Affect Scale (PANAS; Watson et al., 1988) and the Visual Analogue Scale in rating subjective feelings and medical symptoms (VAS; Bond and Lader, 1974; Supplementary Table S3). The WM gating task started 130 minutes after the intake of the drug to maximise the blood plasma concentration of sulpiride, which has a mean time to maximal plasma concentration of about 3 hours with a half-life of about 12 hours (Mehta, 2003). The time window of data acquisition was chosen to be approximately in the middle of the entire test battery, while first data acquisition did not take place earlier than 120 min after the administration of the agent (Mehta, 2003; Mehta et al., 2004; Mehta et al., 2008). In the waiting period, subjects rested and had a snack just before they went into the MRI scanner (10 minutes) in which they completed an anatomical scan (5 minutes) and the WM gating task (4 x 14 minutes). The task was administered in English and was presented on a 32-inch full colour IPS LCD screen (1920 x 1080 resolution, 120 Hz refresh rate). After the lunch break (25 minutes), participants performed a perceptual decision-making task (Spronkers et al., in preparation) and the Simon task (Spronkers et al., in preparation), which are reported elsewhere (75 minutes in total). Finally, subjects’ travel fitness was evaluated by means of a driving impairment test including the Romberg test (Romberg, 1846; Ropper, 1985) a walk and turn test, a one leg stand test, and a finger to nose test. The precise schedules of all three days are described in Supplementary Tables S1-S2.

### Working memory gating paradigm

The cued delayed-response task was inspired by multiple prior studies that have succeeded in capturing striatal activity during input-and output-gating (e.g., Chatham et al., 2014; Schouwenburg et al., 2010; Figure 2). On each trial, subjects were presented two encoding samples: a face and a scene. This was followed by a variable delay interval during which subjects had to maintain the face and/or scene in memory. On different trials they were cued, with a FACE or SCENE cue, to remember either the face or the scene (gating trials, order randomised), or with a BOTH cue to remember both the face and the scene (global trials). During the probe phase, subjects had to indicate whether the probe did or did not match the cued encoding sample (face and/or scene) using a left or right button press with the index versus middle finger of their right hand.

**Figure 2.**
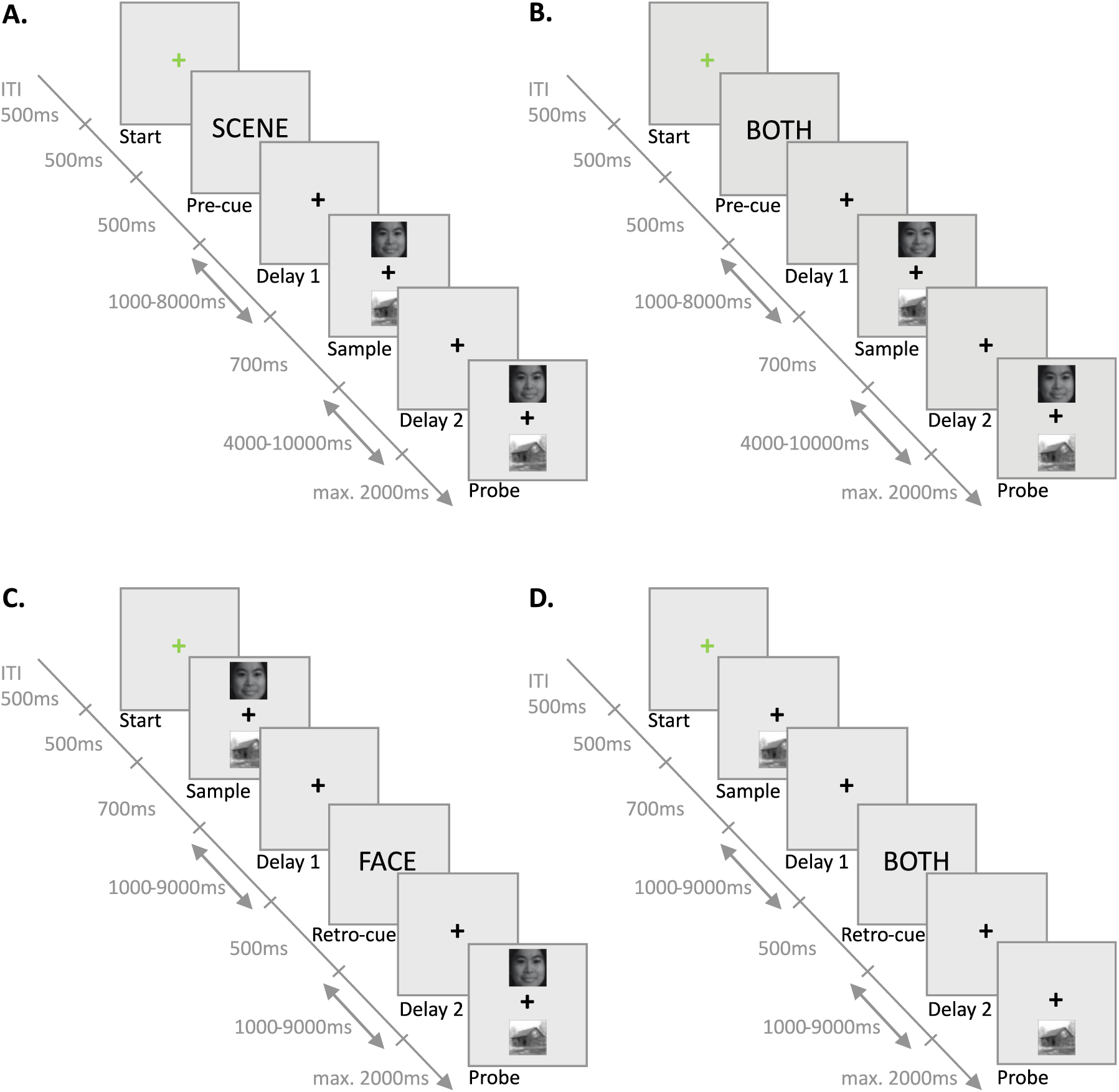
Working memory gating task. This pre-and retro-cue working memory paradigm instructs participants to remember stimuli shown at the sample and match them with stimuli shown at the probe. A. Input-gating of one out of two stimuli following a gating pre-cue (FACE or SCENE) B. Encoding of one or two stimuli following a global pre-cue (BOTH) C. Output-gating of one stimulus at a gating retro-cue (FACE or SCENE) D. Retrieval at a global retro-cue (BOTH). ITI, inter-trial Interval; ms, milliseconds.

Input-and output-gating were implemented in the task using pre-and retro-cues respectively. On pre-cue trials, a FACE, SCENE or BOTH cue was presented prior to the encoding of the sample stimuli. No cue was presented after encoding. Conversely, on retro-cue trials, a FACE, SCENE or BOTH cue was presented after encoding, but no cue was presented prior to encoding. To dissociate effects of WM gating from those of relative cognitive load, the design also included trials on which only one face or one scene was presented during both encoding and probe. These trials were also always accompanied by a BOTH cue (either prior to or post encoding). The effect of cognitive load was captured by comparing these global-load 1 trials with global-load 2 trials, on which both a face and a scene sample were presented together with a BOTH cue (Supplementary Figure S9; Supplementary Table S20).

The task comprised 4 runs (12 minutes per run), with each run consisting of 4 blocks of 12 trials, with the block structure following a 2 x 2 factorial design crossing trial type (pre-cue vs. retro-cue) x selectivity (gating vs. global) (Figures 2A-D). Within the global blocks, we differentiated between global-load 1 and global-load 2 trials (order randomised). The numbers denote the number of sample stimuli presented on the screen at the sample and the probe. Conversely, in the gating condition, two stimuli were always presented, but we distinguished face-gating trials from scene-gating trials. Thus, in total, there were 2 (cue: pre vs. retro) x 2 (global: 1 vs. 2) x 2 (selective: face vs. scene) = 8 different conditions. Participants could take a short break (1-2 minutes) between each of the 4 runs, but the 4 blocks within a run were administered continuously, separated only by a short instruction about the relevant task conditions (pre-cue, retro-cue, global, or gating).

### Input-gating

is captured by the contrast between so-called gating trials, in which pre-cues presented before encoding cued the relevance of one of two stimuli, and so-called global trials, in which pre-cues cued the relevance of both stimuli. First, a green fixation cross (500 ms) was presented, followed by the pre-cue (500 ms). After a jittered delay period (mean_delay1_ = 2926 ms, range_delay1_ = 1002–7990 ms), the stimuli were on the screen for 700 ms. Subsequently, another jittered delay period (mean_delay2_ = 7000 ms, range_delay2_ = 4000–10000 ms) was followed by the probe (max. 2000 ms), to which the participants had to respond with a button press. Reaction times (RT) were recorded from the onset of the probe until the response or until the probe disappeared. The inter-trial interval was 500 ms, following the button press (in case of response) or probe (in case of nonresponse).

### Output-gating

mechanisms were captured with the same gating versus global contrast, with retro-cues were presented after encoding to indicate stimulus relevance at retrieval. On all retro-cue trials, the green fixation cross (500 ms) preceded the stimulus presentation (700 ms), which was followed by a jittered delay period (mean_delay1_ = 3469 ms, range_delay1_ = 1004–8911 ms). After the delay, the retro-cue was presented for 500 ms before another jittered delay period (mean_delay2_ = 3531 ms, range_delay2_ = 1001–8956 ms) preceded the probe (max. 2000 ms).

Delays were randomly sampled and adjusted to the total trial length. Above, we reported the actual delay periods that occurred in the task. For global trials containing two stimuli, a probe was chosen exclusively from one of the two encoded stimulus categories. In the gating condition, by contrast, random stimulus selection allowed probe matches to occur in both task-relevant and task-irrelevant categories within a trial. Overall, matches of the relevant category occurred on 50% of the trials.

### MRI acquisition and preprocessing

Functional brain data were acquired at both pharmacological sessions using a 3T Siemens Magnetom Skyra MRI scanner at the Donders Centre of Cognitive Neuroimaging, Nijmegen, equipped with a 32-channel head coil. Before the start of the task, a whole-brain structural T1-weighted image was acquired using a T1-weighted magnetisation prepared, rapid-acquisition gradient echo sequence (sagittal slices; repetition time, 2300ms; echo time, 3.03ms; voxel size, 1.0×1.0×1.0mm; slice thickness, 1mm; flip angle, 8°; field of view, 256×256mm). During all four runs, functional brain images were obtained using a whole-brain T2-weighted multi-echo gradient-echo echo planar imaging (EPI) sequence (52 slices per volume; repetition time, 1500ms; echo times, 12.4ms, 34.3ms, and 56.2ms; voxel size, 2.5×2.5×2.5mm; interslice gap, 2.5mm; slice thickness, 2.5mm; flip angle, 75°; field of view, 210×210mm). After acquisition, all MRI data were converted from DICOM into NIfTI format using BIDScoin software (version 3.7.2; Zwiers et al., 2022). The fMRIPrep toolbox was used for subsequent preprocessing (version 20.2.6; Esteban et al., 2018c; Esteban et al., 2020). Amongst other steps, this toolbox includes the combination of echoes, realignment, coregistration of functional images to each participant’s T1-weighted anatomical image and spatial normalisation. Further details regarding the fMRIPrep preprocessing can be found in Supplementary Methods. Afterwards, Statistical Parametric Mapping 12 (SPM12; https://www.fil.ion.ucl.ac.uk/spm/software/spm12/) was used to spatially smooth all functional scans with a Gaussian kernel of 8mm full width at half maximum (FWHM).

## Statistical analysis

### Behaviour

To ensure a balanced design between different gating conditions within the task, the data analysis was based on data from all completed runs (*N*_trials_ = 48 x 3 or 4 runs per subject) that were accompanied by neuroimaging data, leading to a removal of in total seven runs from different participants. Subsequently, trials with premature responses (RT < 100 ms) and nonresponse were excluded. More details regarding behavioural data preparation and analysis are presented in Supplementary Methods.

All behavioural analyses were performed with R (version 4.1.0; Team, 2022a) in R Studio (version 1.4.1717; Team, 2022b) with Bayesian mixed-effects regression models using the brms package (version 2.17.0; Bürkner, 2017; Bürkner, 2018; Bürkner, 2021). Accuracy rates were modelled with a Bernoulli distribution and a logit link function, and RTs were modelled with a skewed normal distribution. For interpretability, we report odds ratios (OR; exponential of estimates on the logit scale) for the results for accuracy. For RTs, we report the regular Bayesian model estimates (B). We considered model effects to be statistically significant when the 95% credible interval (CI) did not contain zero for model estimates; and ORs were reported as significant when their 95% CI did not contain one.

The outcome variables were modelled as a function of three independent, categorical predictor variables: selectivity (gating vs. global 1 vs. global 2), drug (sulpiride vs. placebo), and trial type (pre-cue vs. retro-cue). The three-level factor selectivity was coded using simple contrast coding with gating as reference category, while the two-level factors were sum-to-zero coded and scaled. Notably, reported effects of gating are reported in the following direction: global 1 > gating and global 2 > gating. Apart from the descriptives, the categorical predictor selectivity was reduced to two levels of interest: gating and global 1. All predictors and the intercept were added to the model as fixed and random effects with subject as a grouping factor (precise model specifications: Supplementary Tables S5-S6).

### fMRI

All fMRI data were analysed with the SPM12 toolbox for MATLAB R2021b (Mathworks Inc.; https://nl.mathworks.com/products/matlab.html). In a general linear model, we included 11 task regressors of interest, modelling the presentation of the cues and the stimuli for both stimulus categories (face and scene) and for all conditions of the task (gating, global 1, and global 2). An additional regressor of interest modelled probe onset. Nuisance regressors included three rotation and three translation realignment parameters, ICA-AROMA components that shared less than 20% of variance with the task components, and confound regressors comprising global signal, global cerebral spinal fluid, global white matter signal, framewise displacement and the first five anatomical principal component noise regressors (aCompCor).

To assess the quality of the MRI data, we checked the raw data and summary reports from MRIQC and fMRIPrep for artefacts. Additionally, in the general linear model of the task data, a button-press control contrast at probe was used to check whether the MRI signal is of sufficient quality. Significant BOLD signal (family-wise error (FWE) corrected p <.05) was required to surface in one of the following three regions defined by the automated anatomical labelling (AAL) atlas using the WFU_PickAtlas toolbox for SPM12 (https://www.nitrc.org/projects/wfu_pickatlas/): (1) left precentral gyrus, left postcentral gyrus, and left supplementary motor area, (2) bilateral fusiform gyrus and (3) bilateral lingual gyrus and bilateral parahippocampal gyrus. For each region, the mean activity was standardised per measurement over participants and sessions. When the mean activity or the number of voxels above the threshold of FWE p <.05 exceeded the 99% confidence interval within at least one region, and when this lack of signal could be related to bad image quality, as assessed by visual inspection of fMRIPrep and MRIQC output, the subject was excluded from the analysis. The subject (*N* = 1) who met this criterion also did not meet the required task performance of at least 50%.

For each pharmacological session and subject, we created a first-level model to model each event of interest separately. For input-and output-gating, we specified baseline contrasts at sample and retro-cue, respectively, for FACE and SCENE cues, as well as for global 1 and 2 BOTH cues. The resulting contrast images were further analysed with regards to the effect of drug, gating selectivity and gating load using a 2 (placebo and sulpiride) x 3 (gating, global 1, and global 2) within-subject ANOVA. We defined contrasts to examine input gating, output gating, and cognitive load at encoding and retrieval, as well as their modulation by drug. Input-gating effects at encoding were assessed by comparing trials with a gating cue indicating which of two simultaneously presented stimuli was relevant to trials in which only a single stimulus was displayed (sample: gating cue > global 1 cue). The output-gating contrast was defined analogously at retrieval (retro-cue: gating cue > global 1 cue). To evaluate whether these gating effects were influenced by cognitive load, we contrasted trials containing two stimuli with those containing one stimulus at both encoding (sample) and retrieval (retro-cue) (global 2 cue > global 1 cue). For each of these contrasts, we additionally tested an interaction with drug. We report striatal activity of voxels FWE-corrected and thresholded at p <.05 after small-volume correction for the whole striatum, which was defined independently using functional connectivity-based parcellation of the striatum (Piray et al., 2017). Activity in the rest of the brain is reported for exploratory purposes only, from clusters thresholded at p <.001 at whole-brain level uncorrected for multiple comparisons.

Also, for validation purposes, we report the main task effects of encoding at pre-cue and sample, of retrieval at retro-cue, and of response selection at probe across the placebo and drug session by modelling each of the task events as baseline contrast in one first-level model per subject. At the group level, a one-sample t-test was performed on these contrast images separately for every task event. Here, we report clusters that reach the threshold of p <.05, FWE corrected for multiple comparisons at the whole-brain level. For full disclosure of the results, we report all voxels exceeding a threshold of p <.001 whole brain (uncorrected) in supplementary tables S11-S22).

### Stimulus specificity in visual association cortex

To extract stimulus-specific activity from task-relevant posterior areas, regions of interest (ROI) were defined for each individual for both stimulus categories, i.e., fusiform face area (FFA) for face stimuli and the parahippocampal place area (PPA) for scene stimuli, following procedures used in prior studies (Van Den Bosch et al., 2022; Schouwenburg et al., 2010). The ROIs were created based on a first-level contrast from a model including both pharmacological sessions to avoid drug bias (FFA: global 1 face > global 1 scene at the probe; PPA: global 1 scene > global 1 face at the probe). A sphere of 3mm around the peak voxel within an anatomically defined mask from the AAL atlas using the WFU_PickAtlas toolbox for SPM12 (https://www.nitrc.org/projects/wfu_pickatlas/) served as ROI for subsequent analyses. For the FFA, the bilateral fusiform gyrus was used as a mask, while the combination of bilateral parahippocampal gyrus and the lingual gyrus was used as a mask for the definition of the PPA.

As preparation for brain-behaviour correlation analyses and for visualisation purposes, a neural stimulus-selectivity index (SSI) was calculated per participant and trial type to express the balance between FFA and PPA signal in response to gating of face and scene stimuli (category x selectivity x brain area). For each subject, we created a contrast for gating of faces (e.g., output-gating: [gating face retro-cue > global 1 face retro-cue]) and gating of scenes (e.g., output-gating: [gating scene retro-cue > global 1 scene retro-cue]), using all input-and output-gating trials. We extracted the contrast estimates from the individually defined FFA and PPA and averaged them across all voxels within the ROI for each subject, trial type and stimulus-category. For each stimulus category, we then subtracted the mean BOLD signal in the irrelevant stimulus-specific posterior area from signal in the relevant corresponding posterior cortex (e.g., scene stimuli, output-gating: PPA [gating scene retro-cue > global 1 scene retro-cue] - FFA [gating scene retro-cue > global 1 scene retro-cue]). Finally, we created a sum score by adding the values for scenes and faces. Positive values of the resulting SSI reflect that the BOLD signal in the relevant posterior area is stronger than BOLD signal in the stimulus-irrelevant posterior cortex during gating of the preferred category for that area.

To assess the effect of sulpiride versus placebo on BOLD signal during gating in the visual association cortex, we modelled BOLD signal (beta values) with a Bayesian mixed-effects regression model for input-and output-gating separately. Drug (sulpiride vs. placebo), category (face vs. scene), selectivity (gating vs. global 1) and brain area (FFA vs. PPA) were modelled as within-subject factors and the participant as grouping factor. The SSI was implemented in the analysis as an interaction between category x selectivity x brain area. Potential outliers were identified at the level of the constituent two-way interactions comprising the three-way interaction of interest, using a common 95% confidence interval as a criterion (Sullivan et al., 2021). These data points were not excluded, but analyses were repeated to verify that inclusion of the potential outliers did not qualitatively change the results. The analysis was executed with R (version 4.1.0; R Core Team, 2024) in R Studio (version 1.4.1717; Team, 2022b) using the brms package (version 2.17.0; Bürkner, 2017; Bürkner, 2018; Bürkner, 2021).

### Signal extraction from striatal peak voxel

To break down sulpiride’s effect on the BOLD signal in the striatum during output-gating of scenes versus faces and to correlate the effect with participants’ behaviour, we extracted the first eigenvariate of the BOLD timeseries from the peak voxel of the corresponding output-gating contrast: [sulpiride > placebo] x [gating retro-cue > global 1 retro-cue] x [scene > face]. We then analysed the extracted signal with a Bayesian linear mixed-effects model separately for each stimulus category, modelling drug and specificity as within-subject factors and participant as grouping factor.

### Brain-behaviour correlations

An exploratory analysis of between-subject brain-behaviour correlations was conducted to examine the correspondence between stimulus-specific activity in task-relevant visual association cortex and behaviour across participants. Separately for sulpiride and placebo, we correlated the SSI for each gating-type and participant with their reaction time and accuracy score, averaged across stimulus category. Similarly, correlation analyses were executed between behaviour and the extracted signal from the striatal peak voxel identified in the drug × gating interaction contrast during output-gating. We used a Pearson correlation analysis using the stats package (version 3.6.2; R Core Team, 2024) in R (version 4.1.0; Team, 2022a).

## Results

The task activated the expected networks of interest during pre-cue, encoding, retrieval and probe (response) (Figure 3; Supplementary Tables S11-S14). Throughout the task, areas of the fronto-parietal control network were increased, including the dorsolateral prefrontal cortex (DLPFC), the intraparietal sulcus (IPS) and the posterior parietal cortex (PPC). As expected, we also observed strong BOLD signal in the FFA and PPA responding to pictures of faces and scenes, respectively, during encoding and at probe when these images were presented in isolation on the screen (i.e., global 1 trials, Supplementary Tables S21-S22).

**Figure 3.**
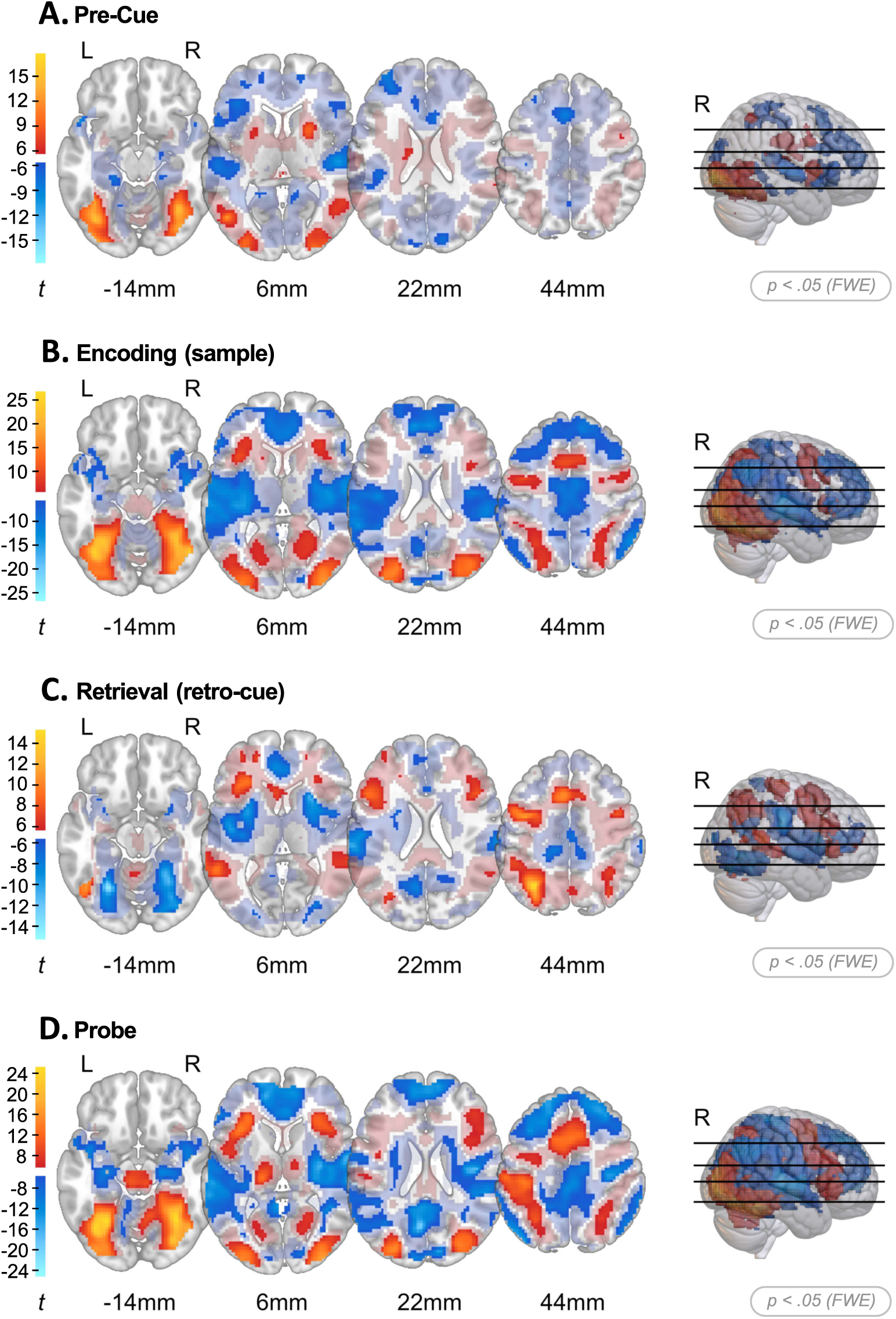
BOLD signal during different phases of the task. Whole brain t-map showing voxels after family-wise error correction at p <.05 whole brain in full opacity. BOLD signal increases compared with baseline are shown in warm colours while BOLD signal decreases are shown in cold colours. In addition, low-intensity, transparent colours present voxels below threshold (p <.1 unc. whole brain) for visualisation purposes. A. Pre-cue, striatal signal increase B. Encoding, striatal signal increase C. Retrieval, striatal signal decrease D. Probe. L, left hemisphere; R, right hemisphere; FWE, family-wise error; unc, uncorrected.

### Behavioural evidence for gating task compliance

Behavioural results confirmed that the gating manipulation was successful (Figure 4; Supplementary Tables S5-S6). Participants were faster and more accurate after gating one of two stimuli into WM compared with keeping two stimuli in WM (global 2 vs. gating; accuracy: odds ratio [OR] = 0.42, 95% credible interval [CI] = [0.34, 0.52]; RT: estimate [B] = 130.80, CI = [113.55, 148.28]). However, compared with encoding or retrieving only one stimulus, gating one of two stimuli induced a gating cost: performance on gating trials was significantly lower than on trials on which only one stimulus was presented (global 1 vs. gating; accuracy: OR = 1.41, CI = [1.18, 1.70]; RT: B = *−*25.93, CI = [*−*37.55*, −*14.10]). The selective D_2_ receptor antagonist, sulpiride, did not significantly affect participants’ overall accuracy and RTs compared with placebo, when averaged across gating and stimulus conditions (accuracy: OR = 0.88, CI = [0.71, 1.08]; RT: B = *−*5.21, CI = [*−*25.00, 14.27]; Figure 4).

**Figure 4.**
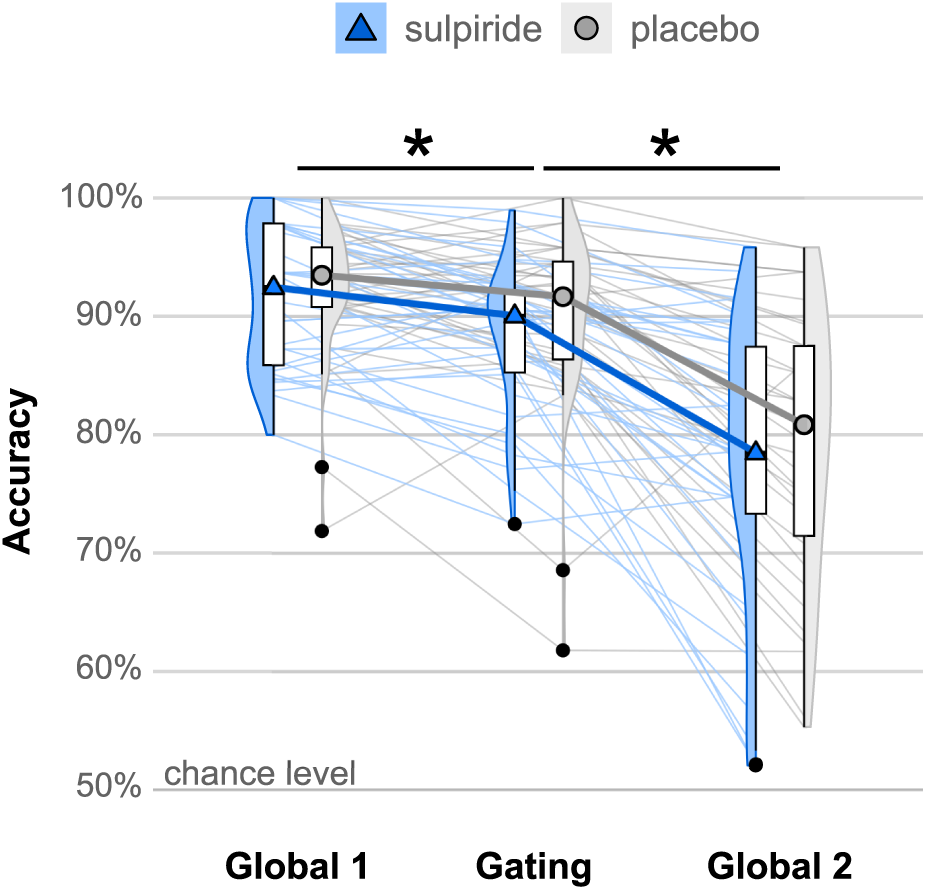
Behavioural performance on the task. Accuracy scores for each participant for every condition, averaged across input-and output-gating trials. Participants perform better when prioritising only one image (global 1) than when gating one of two images (gating), indicating a gating cost. There was also a benefit of gating one of two stimuli. On gating trials participants performed better than on global 2 trials where two stimuli needed to be kept in WM.

Sulpiride affected response speed for input-and output-gating (global 1 vs. gating) differently (selectivity_global1-gating_ x trial type_pre-retro_ x drug_sulpiride-placebo_; RT: B = 36.99, CI = [3.45, 70.38]), driven by reduced RT gating benefits on input-gating (selectivity_global1-gating_ x drug_sulpiride-placebo_; RT: B = 24.93, CI = [0.76, 48.93]; Supplementary Tables S7-S8) but not output-gating trials (selectivity_global1-gating_ x drug_sulpiride-placebo_; RT: B = *−*12.13, CI = [*−*37.40, 13.33]; Supplementary Tables S9-S10).

### Enhanced stimulus-selectivity in visual association cortex during input-and output-gating

To es-tablish that task-relevant representations were successfully gated (irrespective of drug session), we investigated effects of gating on a neural index of stimulus-selectivity in visual association cortex. Specifically, we asked whether stimulus-selective activity in visual association cortex was enhanced when the preferred stimulus was gated. The SSI represents a relative enhancement of BOLD signal during gating of scenes in the PPA (vs. FFA) and faces in the FFA (vs. PPA). As predicted, both input-and output-gating enhanced stimulus-selectivity in the FFA and PPA (area_FFA-PPA_ *×* category_face-scene_ *×* selectivity_global1-gating_: B_input-gating_ = −8.28, CI_input-gating_ = [−12.16, −4.43]; area_FFA-PPA_ × category_face-scene_ × selectivity_global1-gating_: B_output-gating_ = *−*11.19, CI_output-gating_ = [*−*14.02*, −*8.34]; Figure 7D).

Breaking down this three-way interaction during input-gating shows that this effect was driven by a boost in the PPA response during gating of scenes (compared with faces) (category_face-scene_ *×* selectivity_global1-gating_: B_PPA_ = 7.21, CI_PPA_ = [4.23, 10.15]; B_FFA_ = *−*1.01, CI_FFA_ = [*−*2.86, 0.85]).

Conversely, during output-gating, the interaction effect was present for both FFA and PPA. At the retro-cue, signal in both visual association cortices was boosted when the preferred stimuli were output-gated (category_face-scene_ *×* selectivity_global1-gating_: B_PPA_ = 7.21, CI_PPA_ = [4.83, 9.60]; B_FFA_ = *−*3.99, CI_FFA_ = [*−*5.43*, −*2.57]).

When modelling the absolute difference between FFA and PPA responses, we observed a general face bias during retrieval: The absolute difference between FFA–PPA was greater for faces than for scenes (category_face-scene_: B_retrieval_ = 2.60, CI_retrieval_ = [1.32, 3.87]). This face bias in visual association cortex varied as a function of gating (category_face-scene_ *×* selectivity_global1-gating_: B_output-gating_ = *−*7.27, CI_output-gating_ = [*−*9.13*, −*5.39]).

### Effects of output-gating on stimulus-selectivity in visual association cortex correlate with be-havioural gating benefits

Between-subject brain–behaviour correlations corroborated the behavioural relevance of the effect of gating on the neural SSI. The neural SSI in visual association cortex during output-gating was associated significantly with gating-related response speed, with negative correlations, of moderate effect size, observed in both the sulpiride (Pearson correlation *r*_RT_*_×_*_SSI_(32) = *−.*43, *p* = 0.012) and placebo conditions (*r*_RT_*_×_*_SSI_(32) = *−.*40, *p* = 0.020), but not with accuracy (sulpiride: *r*_accuracy_*_×_*_SSI_(32) =.26, *p* = 0.143; placebo: *r*_accuracy_*_×_*_SSI_(32) =.28, *p* = 0.110; Figure 5). Participants with a greater neural selectivity index exhibited faster (but not significantly) more accurate responses on output-gating trials compared with global 1 trials. Stimulus-selectivity during input-gating, instead, mostly did not correlate with input-gating effects on performance across drug conditions (sulpiride: *r*_RT_*_×_*_SSI_(32) =.01, *p* = 0.970; placebo: *r*_RT_*_×_*_SSI_(32) =.03, *p* = 0.860; sulpiride: *r*_accuracy_*_×_*_SSI_(32) =.09, *p* = 0.594). However, in the placebo condition, effects of input-gating on accuracy correlated moderately with the effect of input-gating on neural selectivity (placebo: *r*_accuracy_*_×_*_SSI_(32) =.41, *p* =.016), confirming the correspondence between behavioural measurements and neural BOLD signal (Figure 5).

**Figure 5.**
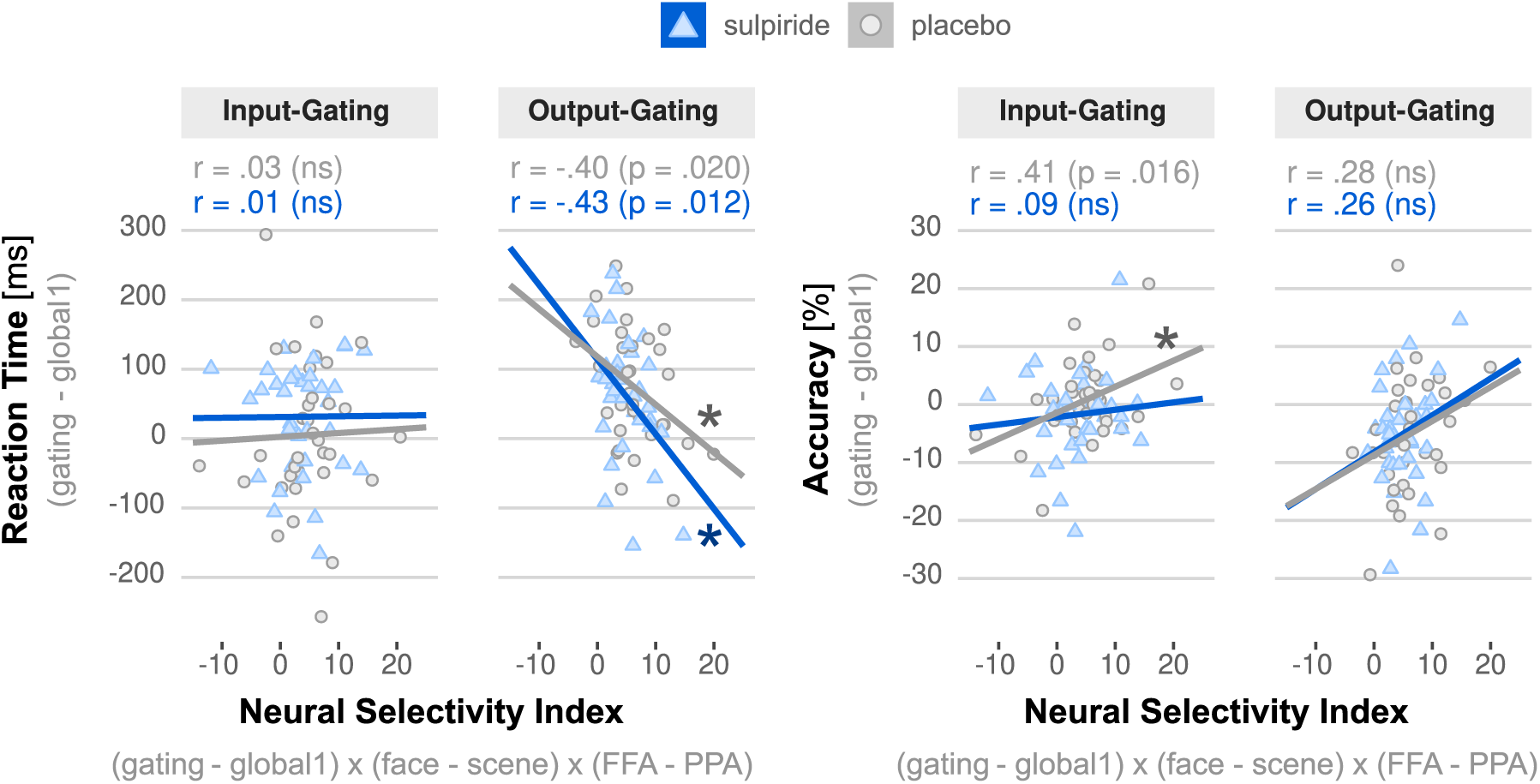
Brain-behaviour correlations between stimulus-selectivity in posterior areas and task performance under sulpiride and placebo. Reaction time and accuracy scores for each participant and drug condition are shown as a function of neural selectivity in stimulus-association cortex. Significant correlations with moderate effect sizes were observed between reaction time and neural selectivity during output gating under both sulpiride and placebo, and between accuracy and neural selectivity during input gating under placebo. No other correlations reached significance. ms, milliseconds; FFA, fusiform face area; PPA, parahippocampal place area; r, Pearson correlation index; *, p <.05; ns, nonsignificant (p >.05).

### Increases and decreases in striatal BOLD signal during WM input-and output-gating

Inde-pendent of the drug manipulation, input-gating (gating vs. global during encoding) was associated with BOLD signal changes in a broad fronto–parieto–visual network of brain regions, including the dlPFC, IPS, insula, and PPC (Figure 6A; Supplementary Table S15). Critically, in line with our predictions, striatal BOLD signal was increased bilaterally in the caudate nucleus during WM input-gating (gating vs. global 1; small-volume correction (SVC) over the whole striatum; left caudate nucleus: *N*_voxels_ = 70, *x, y, z* = (*−*16, 12, 9), *Z* = 6.31, *p*_peak FWE SVC_ *<.*001; right caudate nucleus: *N*_voxels_ = 14, *x, y, z* = (16, 12, 9), *Z* = 4.43, *p*_peak FWE SVC_ =.003). Conversely, in contrast to our hypothesis, during output-gating, striatal BOLD signal was significantly decreased bilaterally in the puta-men and unilaterally in the caudate nucleus (gating vs. global 1; right putamen: *N*_voxels_ = 320, *x, y, z* = (21, 8, 6), *Z* = *−∞*, *p*_peak FWE SVC_ *<.*001; left putamen: *N*_voxels_ = 301, *x, y, z* = (*−*26, 2*, −*8), *Z* = *−∞*, *p*_peak FWE SVC_ *<.*001; right caudate: *N*_voxels_ = 105, *x, y, z* = (18, 15, 9), *Z* = *−∞*, *p*_peak FWE SVC_ *<.*001). Notably, this decrease of striatal BOLD signal was observed in the presence of increases in the same broad fronto–parieto–visual network that was also activated during input-gating (Figure 6B; Supplementary Table S16). Comparison of the input-and output-gating contrasts between faces and scenes suggests that gating-related BOLD signal, e.g., in the fronto-parietal control network, was relatively similar for faces and scenes (Supplementary Figure S10). These results were replicated when the analysis was restricted to the placebo condition (Supplementary Results).

**Figure 6.**
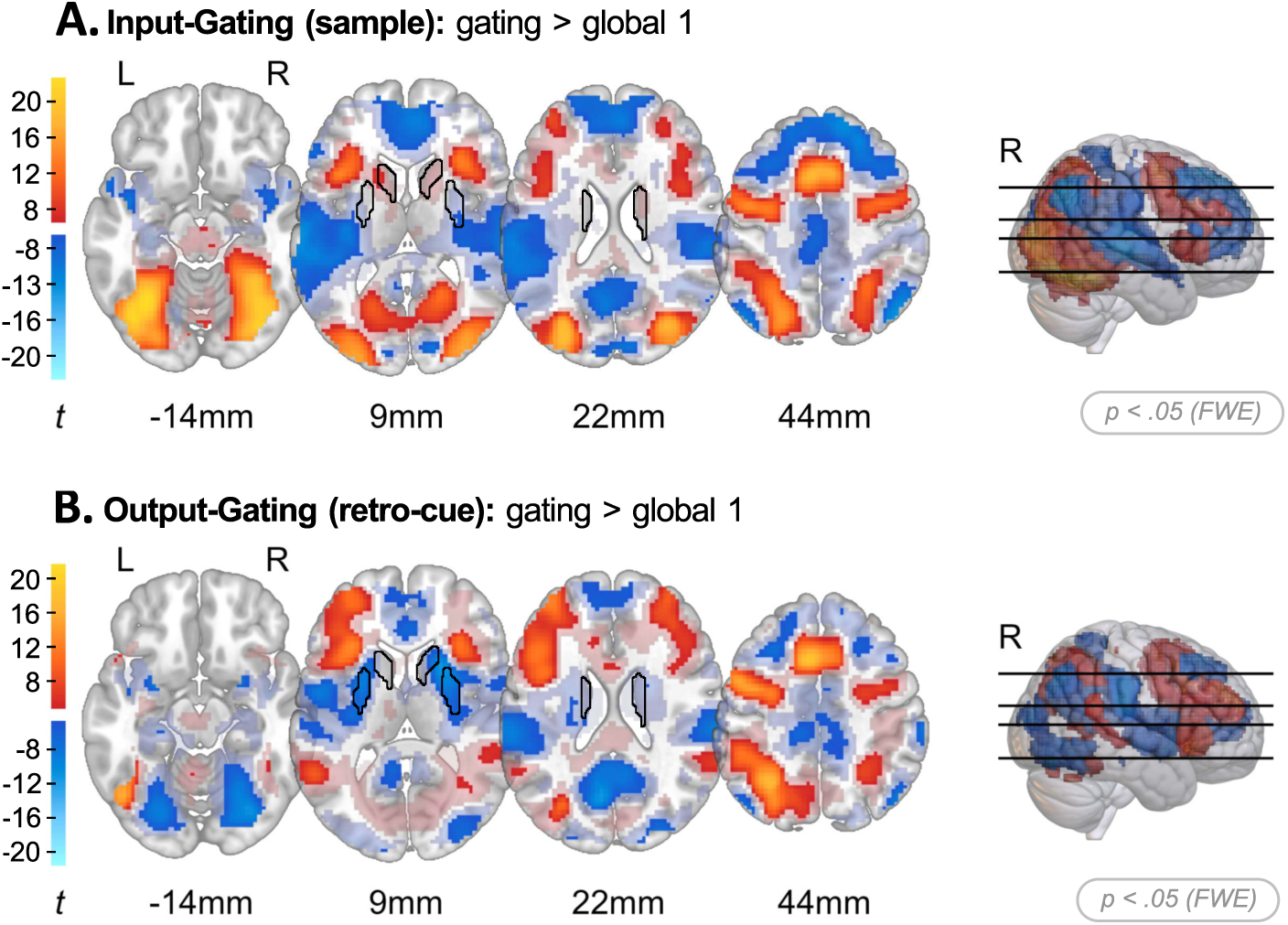
Input-and output-gating show different patterns of responses in the basal ganglia. Whole brain t-map showing voxels after family-wise error correction at p <.05 whole brain in full opacity. BOLD signal increases compared with baseline are shown in warm colours while BOLD signal decreases are shown in cold colours. The striatal mask used for small volume correction is outlined in black. In addition, low-intensity, transparent colours present voxels below threshold (p <.1 unc. whole brain) for visualisation purposes.**A.** Input-gating, displaying the BOLD signal at sample for gating > global 1 trials, with an increase in striatal BOLD signal. **B.** Output-gating, showing the BOLD signal at retro-cue for gating > global 1 trials, striatal BOLD signal is decreased. L, left hemisphere; R, right hemisphere; FWE, family-wise error; unc, uncorrected.

**Figure 7.**
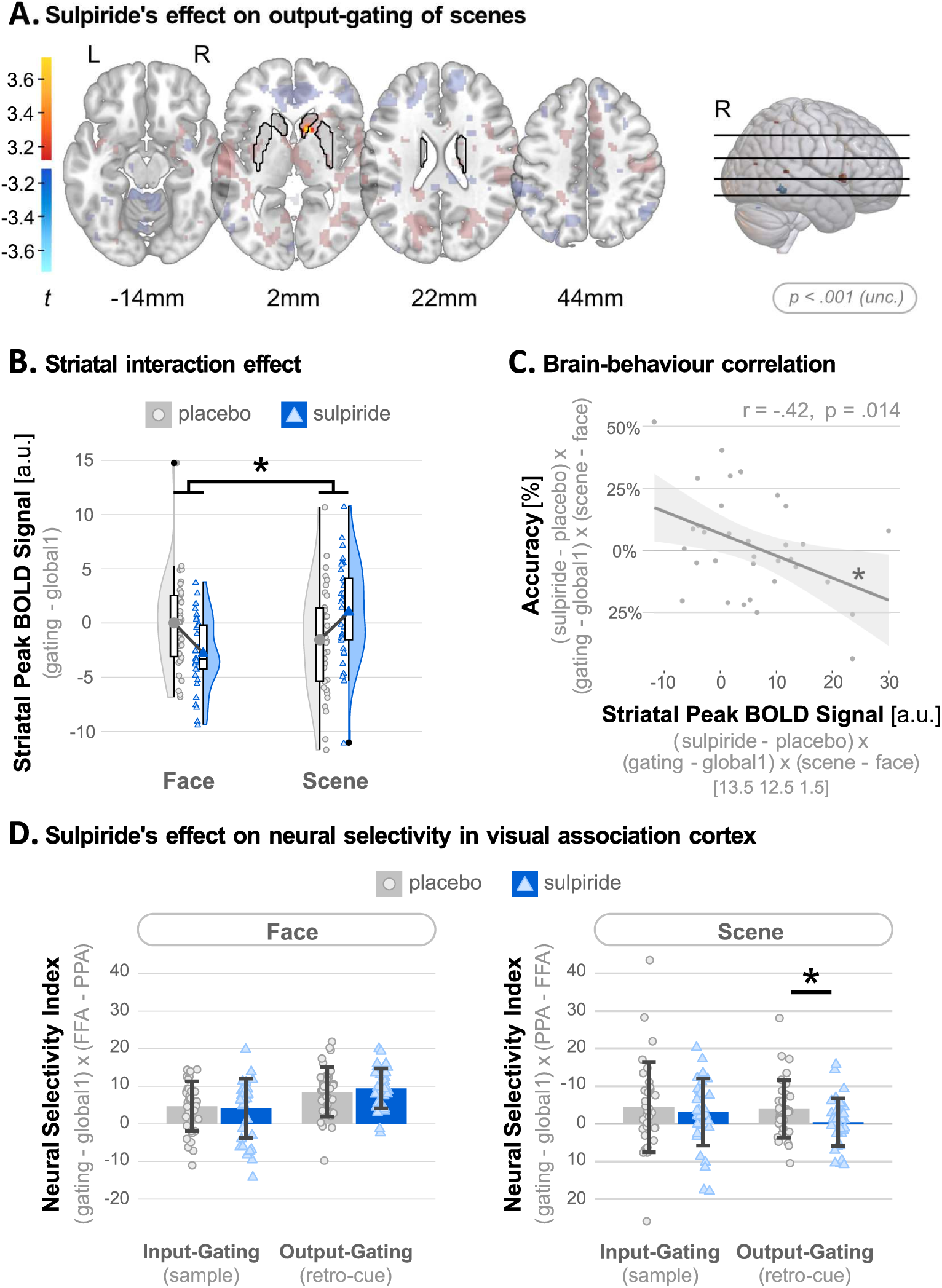
Sulpiride increased striatal BOLD signal during output-gating of scenes versus faces. A. Whole brain contrasts showing BOLD signal at p <.001 uncorrected in full opacity ([sulpiride > placebo] x [scenes > faces] x [gating > global 1] at retro-cue). BOLD signal increases compared with baseline are shown in warm colours while BOLD signal decreases are shown in cold colours. The striatal mask used for small volume correction is outlined in black. In addition, low-intensity, transparent colours present voxels below threshold (p <.1 unc. whole brain) for visualisation purposes. B. BOLD signal extracted from the striatal peak voxel, broken down into drug and stimulus category. C. Accuracy on the task correlated with BOLD signal extracted from the striatal peak voxel, such that a greater increase in striatal BOLD signal correlated with lower accuracy. D. Stimulus-selectivity in the PPA (versus FFA) was disrupted by sulpiride during the gating of scene stimuli during retrieval. L, left hemisphere; R, right hemisphere; unc, uncorrected; a.u., arbitrary unit.

### Sulpiride disrupts neural stimulus-specificity during output-gating of scene stimuli

Partly in line with our gating hypothesis, sulpiride impaired gating-related stimulus-specificity in visual association areas during output-gating, but this was seen only for scene stimuli (scene only: drug_sulpiride–placebo_ × area_PPA–FFA_ × selectivity_global1–selective_: B_output-gating_ = −3.51, CI_output-gating_ = [−6.95, −0.07]), and not for face stimuli (face only: drug_sulpiride–placebo_ × area_FFA–PPA_ × selectivity_global1–selective_: B_output-gating_ = *−*0.89, CI_output-gating_ = [*−*3.95, 2.10]; Figure 7D; Supplementary Table S19). As a function of input-gating, there were no effects of sulpiride on the responses to any gated category in the FFA or PPA (scene only: _drugsulpiride–placebo_ × area_PPA–FFA_ × selectivity_global1–gating_: B_input-gating_ = −1.27, CI_input-gating_ = [*−*5.48, 2.98]; face only: drug_sulpiride–placebo_ *×* area_FFA–PPA_ *×* selectivity_global1–gating_: B_input-gating_ = 0.53, CI_input-gating_ = [*−*2.85, 3.88]).

### Sulpiride enhances retrieval-related BOLD signal in the striatum for scenes

As predicted, there was a significant effect of sulpiride on striatal BOLD signal, specifically, in the right caudate nu-cleus, as a function of output-gating (gating retro-cues vs. global 1 retro-cues), but again this effect was qualified by stimulus category: It differed between face and scene stimuli (drug_sulpiride–placebo_ *×* category_face–scene_ *×* selectivity_global1–gating_; right caudate nucleus: *N*_voxels_ = 13, *x, y, z* = (11, 12, 2), *Z* = 3.69, *p*_peak FWE SVC_ =.046; Figure 7A). Exploration of the first eigenvariate from the peak voxel of this interaction effect showed that it was driven by a sulpiride-induced increase of output-gating related signal for scenes (drug_sulpiride–placebo_ *×* selectivity_global1–gating_: B_SCENE_ = *−*3.06, CI_SCENE_ = [*−*5.93*, −*0.24]; Figure 7B), but a decrease for faces (drug_sulpiride–placebo_ *×* selectivity_global1–gating_: B_FACE_ = 3.02, CI_FACE_ = [0.93, 5.09]). The interaction effect remained significant after the removal of outliers that have been defined as data points falling outside the 95% confidence interval of any relevant two-way interaction. No main effects of sulpiride on striatal BOLD signal were observed across gating conditions (Supplementary Table S18), nor were effects detected in other brain regions during input-or output-gating. During encoding, however, sulpiride modulated striatal BOLD signal in the caudate nucleus (Supplementary Figure S8; Supplementary Table S17), but not during retrieval.

Individual differences in the effect of sulpiride on stimulus-specific gating signal in the striatum (extracted from the peak voxel revealed by the drug x stimulus category x gating interaction contrast) correlated with individual differences in the effect of sulpiride on behavioural stimulus-specific output-gating benefits. Specifically, participants who exhibited greater gating-related increases in striatal signal for scenes (vs. faces) after sulpiride also showed greater disruption of gating-related accuracy benefits for scenes (vs. faces) after sulpiride (drug_sulpiride–placebo_ *×* category_face–scene_ *×* selectivity_global1–gating_: *r*_striatal peak_*_×_*_accuracy_(32) = *−*0.42, *p* =.014; Figure 7C). The more sulpiride increased striatal BOLD signal, the more it impaired gating. Reaction times (RTs), did not correlate with the striatal peak BOLD signal (drug_sulpiride–placebo_ × category_face–scene_ × selectivity_global1–gating_: r_striatal peak×RT_(32) = −0.03, *p* =.87).

## Discussion

Dopamine is well established to contribute to WM (Sawaguchi and Goldman-Rakic, 1991; Williams and Goldman-Rakic, 1995; Cools and Arnsten, 2022; Cools and D’Esposito, 2011), but the specific mechanisms by which dopamine alters WM control have remained unclear. Empirical evidence for a role of striatal dopamine in the output-gating of WM has been lacking. Here, we demonstrate that changing dopamine with sulpiride alters striatal activity during output gating, accompanied by a sulpiride-induced reduced neural precision of gated sensory representations in visual association cortex, and lower behavioural accuracy. Together, the results establish a role for striatal dopamine in controlling cortical WM outflow.

Our hypotheses were inspired by current WM gating models (Badre and Frank, 2012; Hazy et al., 2007; Kessler, 2017; O’Reilly and Frank, 2006; Rac-Lubashevsky and Frank, 2021; Trutti et al., 2024; O’Reilly et al., 2024), according to which striatal dopamine controls the gating of both input and output of WM. We assessed effects of the selective dopamine D2 receptor antagonist sulpiride on striatal BOLD signal during the performance of a task separating input-and output-gating of WM. Partly supporting these models, our results reveal that sulpiride increased neural activity in the caudate nucleus during output-gating of scenes, albeit not faces. Importantly, this effect was accompanied by sulpiride disrupting gating-related activity in stimulus-specific posterior cortex, again for scenes only. This sulpiride-induced increase in striatal activity correlated with sulpiride-induced decreases in behavioural gating benefits, indicating that sulpiride-related disruption of output-gating was accompanied by a sulpiride-related increase of striatal signal.

This pattern of effects generally concurs with our hypothesis that dopamine acts on the striatum while regulating cortical output of WM. The striatal gating theory underlying this hypothesis posits that striatal dopamine alters gating by regulating the balance between Go and NoGo pathways of the fronto-thalamic basal ganglia circuit (Badre and Frank, 2012; Frank and Badre, 2012; Hazy et al., 2007). This account predicts, however, increased striatal activity under high gating demands, which was not the case. Instead, striatal activity was decreased during gating versus global retrieval (Figure 6B). Importantly, the output-gating effect was negative, contrasting with prior findings of striatal signal increases rather than decreases during WM updating and gating-related operations (Chatham et al., 2014; Nir-Cohen et al., 2020; Nir-Cohen et al., 2023; Trutti et al., 2024). It is also inconsistent with the hypothesis that the striatum facilitates gating by jointly enhancing Go and suppressing NoGo pathway activity (Schouwenburg et al., 2015).

However, instead of recruiting selective WM gating mechanism that resolve competition between relevant and irrelevant WM representation, our paradigm captured nonselective input-and output-gating operations, unlike those captured by continuous updating paradigms that require updating or retrieval of some and maintenance of other items in WM (Badre and Frank, 2012; Nir-Cohen et al., 2020; Nir-Cohen et al., 2023). Accordingly, the absence of striatal increases in our output-gating manipulation does not undermine predictions of selective gating models, including the PBWM model. Instead, it raises the question whether the observed sulpiride effects reflect changes in nonspecific retrieval processes. Thus, while the present study demonstrates a role for striatal dopamine during the nonselective gating of cortical WM output, the question whether striatal dopamine is similarly required for the selective form of WM gating that requires more sophisticated push-pull mechanisms remains unanswered.

One possibility is that sulpiride effects on striatal activity during the nonselective gating reflect reduced cognitive control effort. The main striatal effect during output-gating might reflect greater retrieval demands on global than gating trials. Gating retro-cues instructed the retrieval of either the face or scene category, while global retro-cues instructed the retrieval of both categories (global 1 and global 2). The sulpiride-related enhancement of striatal activity for output-gating of scenes (versus faces), coupled with disrupted stimulus selectivity in visual cortex for scenes, may reflect greater difficulty and lower salience of scenes relative to faces (Bollinger et al., 2010; Lepsien and Nobre, 2007). Accordingly, dopamine has been linked to effort-based control allocation, with D2 receptors implicated in signalling cognitive control costs (Collins and Frank, 2014; Westbrook et al., 2020). Relatedly, the striatum has been proposed to implement evidence accumulation over time rather than a pure gating mechanism (O’Reilly et al., 2024; Bogacz and Gurney, 2007; Dunovan et al., 2015; Doi et al., 2020; Yartsev et al., 2018). In hindsight, we hypothesise that sulpiride modulates striatal activity by disrupting controlled retrieval, leading to a bias away from the nonsalient scenes towards the salient faces.

In contrast to its effect on output-gating, sulpiride had no effect on input-gating (no difference between gating and global 1 trials). This was despite both input-and output-gating manipulations being successful: Consistent with previous research (Lewis-Peacock and Norman, 2014; Nir-Cohen et al., 2020; Rose et al., 2016), gating enhanced stimulus-specific neural activity in posterior visual association cortex during both input-and output-gating. As in previous studies (Bollinger et al., 2010; Chatham et al., 2014; Wallis et al., 2001), gating improved performance, increasing accuracy and reducing RTs on pre-and retro-cue trials requiring gating of one of two stimuli, compared with global 2 trials, which required maintenance of both. These behavioural benefits suggest reduced cognitive load under gating. Moreover, greater gating-related increases in stimulus selectivity in visual association cortex correlated with faster responses on gating versus global trials. Together, these findings demonstrate the effectiveness of both gating processes and replicate attentional gain effects in visual association cortex (Desimone and Duncan, 1995; D’Esposito and Postle, 2015; Oberauer and Hein, 2012; Schouwenburg et al., 2015; Schouwenburg et al., 2010). Notably, the absence of a sulpiride effect on striatal activity during input-gating is striking. We speculate this reflects the nonselective and relatively low-cost nature of input-gating, which, in the present study, may rely on early attentional filtering prior to perceptual encoding, whereas output-gating likely engages more demanding executive processes involving the striatum for active retrieval from WM. One limitation of the current task design is that it primarily captured nonselective input-and output-gating rather than the selective gating processes proposed by WM gating models (Badre and Frank, 2012; Nir-Cohen et al., 2020; Nir-Cohen et al., 2023). Future studies should use adapted reference back tasks better suited to capture selective control operations as new information must be incorporated into WM while other items are maintained and protected (Kessler, 2017; Rac-Lubashevsky and Frank, 2021; Trutti et al., 2024). By independently manipulating selective gating and retrieval demands in these paradigms, it could be determined whether striatal dopamine supports selective push-pull gating mechanisms or instead acts on retrieval effort-based cognitive control mechanisms.

The effect of sulpiride on output-gating-related striatal signal varied by stimulus category. Sulpiride attenuated the striatal decrease during output-gating of scenes but not faces. This may reflect differential dynamic range in stimulus processing. Previous studies have reported better WM performance for faces than scenes (Bollinger et al., 2010; Lepsien and Nobre, 2007), consistent with our finding of higher accuracy and faster responses for faces (Supplementary Tables S7-S10), suggesting a face-processing bias reflected in higher signal-to-noise ratios. Supporting this behavioural effect, our neuroimaging results and prior studies show stronger face selectivity in the FFA compared with scene/object-selective posterior regions (Druzgal and D’Esposito, 2003; Ishai et al., 2000; Lin et al., 2019). We speculate that this intrinsic face bias limited further sulpiride-induced increases in signal-to-noise ratio for faces, making the subtle enhancing dopaminergic effects on retrieval only measurable for scenes. Although dopaminergic effects may vary by sex (Zachry et al., 2021), the present sample was underpowered to reliably examine sex differences.

Finally, a limitation D2-receptor agents such as sulpiride is uncertainty about whether they primarily block postsynaptic D2 receptors, mimicking reduced endogenous dopamine, or presynaptic D2 (auto)receptors, located on striatal terminals and midbrain dopamine neurons (Ford, 2014). Postsynaptic blockade is known to disinhibit striatal NoGo pathway activity, strengthening resistance to flexible gating (Frank et al., 2001). Instead, presynaptic blockade would increase dopamine release and firing, which is proposed to instead inhibit the NoGo pathway and facilitate flexible gating (Ago et al., 2005; Chavanon et al., 2013; Serra et al., 1990; Richfield et al., 1989; Schoemaker et al., 1997; Frank and O’Reilly, 2006; Moustafa et al., 2008). The 400 mg dose used here is not large enough to exclude presynaptic effects (Eisenegger et al., 2014; Westbrook et al., 2021). However, the observed pattern during output-gating is more consistent with a postsynaptic D2 receptor blockade (Frank et al., 2001; Frank and O’Reilly, 2006), given reduced performance, increased striatal activity during gating versus global trials, and attenuated neural selectivity in visual association cortex.

In conclusion, in the present pharmacological fMRI study, sulpiride decreased neural precision in visual cortex, while increasing striatal BOLD signal during retrieval of nonsalient versus salient stimuli. This drug-induced increase in retrieval-related striatal activity correlated with poorer WM output-gating of nonsalient stimuli. This finding suggests that the striatum may not directly control WM gating but instead track its effort cost in a dopamine D2-dependent manner. This aligns with work on the expected value of cognitive control, implicating D2 receptors specifically in the costs rather than the benefits of cognitive effort (Westbrook et al., 2020; Collins and Frank, 2014).

