## Supplement for "Dopamine modulates striatal activity when controlling the output of working memory"

#### Supplementary Methods

**Control analyses.** To ensure that the effects of sulpiride were not confounded by differences in physiological measures or subjective ratings from mood and medical questionnaires, we tested differences between the drug and placebo session using MANOVA in R (version 4.1.0; Team, 2022a) in R Studio (version 1.4.1717; Team, 2022b). For each measure, we calculated the difference between baseline (time point 1, immediately before drug administration) and after the waiting period of 70 minutes (time point 2) and used these as outcome variables with drug as predictor in the model. Moreover, we estimated a linear mixed effects model to test for differences in performance on the Digit Span Task between drug and placebo, with direction (forward vs. backward) as second predictor for the total number of remembered numerical sequences, and a random intercept for each subject. In addition, we used the James Blinding Index (James et al., 1996) to evaluate whether participants correctly guessed in which session they received the drug and in which placebo by using the BI package (version 1.2.0; Schwartz and Mercaldo, 2021) in R. More importantly, to check whether sulpiride indeed acted on the striatum as predicted by previous literature, we contrasted all task events from the drug session against all task events from the placebo session.

**Exclusions.** Participants were excluded based on overall performance averaged across the four task runs. Thus, performance could fall below 50% accuracy in a single run if higher performance in the remaining runs counterbalanced it. This occurred once: one participant performed below chance in a run (47.62% correct). Overall, three runs from different participants were removed because they consisted of fewer trials than a complete run ($N_{\text{trials}} < 48$). In total, $5.64\%$ of the remaining trials ($N_{\text{trials}} = 13{,}872$) were excluded from analysis due to nonresponse, and two trials were removed due to premature responses ($RT_{\text{premature}} = \{13.78\,\text{ms}, 94.43\,\text{ms}\}$). Three additional trials were excluded because participants pressed neither of the two specified buttons (buttons 1 and 2).

**Response recoding.** According to our coding scheme for buttons 1 and 2 (button 3 was always coded as incorrect), a period in which participants shifted their fingers one button to the right was characterised by a high error rate because participants responded correctly but in a reversed manner. Seven participants accidentally shifted their fingers one button to the right, pressing buttons 2 and 3 for a period of time. To prevent further data exclusion and potential bias, we recoded their responses for this time period. To differentiate errors from shifted responses, we applied the following recoding strategy. We first defined the period of shifted responses by identifying trials with shifted button presses (button 3) as well as previous and following trials with a correct left button press (button 1). Within this period, we calculated the error rate and compared it to the participant's error rate on the remaining trials. When the error rate within the shifted-response period was not included within the 90% confidence interval of the error rate on the remaining trials, performance during the shifted-response period was significantly worse due to inverse button presses. Therefore, participants actually performed equally well during this period, and these responses were recoded. Four periods of trials were recoded ($N_{\text{trials}} = 85$), distributed across four participants. When this recoding criterion did not apply, the trials were removed from analysis. This occurred for three periods ($N_{\text{trials}} = 27$) from different participants.

**Extended behavioural models.** To further explore the behavioural interaction effect between drug, gating and trial type and relate it to the neuroimaging results, we additionally specified separate accuracy and RT models for input- and output-gating including stimulus category (face vs. scene) as a fixed effect (Supplementary Tables S7–S10).

#### fMRIPrep Preprocessing

Results included in this manuscript come from preprocessing performed using `fMRIPrep` 20.2.6 (Esteban et al., 2018b; Esteban et al., 2018a; RRID:SCR_016216), which is based on `Nipype` 1.7.0 (Gorgolewski et al., 2011; Gorgolewski et al., 2018; RRID:SCR_002502).

**Anatomical data preprocessing.** A total of 2 T1-weighted (T1w) images were found within the input BIDS dataset. All of them were corrected for intensity non-uniformity (INU) with N4BiasFieldCorrection Tustison et al., 2010, distributed with ANTs 2.3.3 (Avants et al., 2008; RRID: SCR_004757). The T1w reference was then skull-stripped with a Nipype implementation of the `antsBrainExtraction.sh` workflow (from ANTs), using OASIS30ANTs as the target template. Brain tissue segmentation of cerebrospinal fluid (CSF), white matter (WM), and gray matter (GM) was performed on the brain-extracted T1w image using FAST (FSL 5.0.9, RRID: SCR_002823; Zhang et al., 2001).

A T1w reference map was computed after registration of the two T1w images (after INU correction) using `mri_robust_template` (FreeSurfer 6.0.1; Reuter et al., 2010). Volume-based spatial normalization to one standard space (MNI152NLin2009cAsym) was performed through nonlinear registration with `antsRegistration` (ANTs 2.3.3), using brain-extracted versions of both the T1w reference and the T1w template. The following template was selected for spatial normalization: ICBM 152 Nonlinear Asymmetrical template version 2009c (Fonov et al., 2009; RRID: SCR_008796, TemplateFlow: MNI152NLin2009cAsym).

**Functional data preprocessing.** For each of the 8 BOLD runs found per subject (across all tasks and sessions), the following preprocessing was performed. First, a reference volume and its skull-stripped version were generated by aligning and averaging the first echo of 3 single-band references (SBRefs). Susceptibility distortion correction (SDC) was omitted. The BOLD reference was then co-registered to the T1w reference using `flirt` (FSL 5.0.9; Jenkinson and Smith, 2001) with the boundary-based registration cost function Greve and Fischl, 2009. Co-registration was configured with nine degrees of freedom to account for distortions remaining in the BOLD reference. Head-motion parameters with respect to the BOLD reference (transformation matrices, and six corresponding rotation and translation parameters) were estimated before any spatiotemporal filtering using `mcflirt` (FSL 5.0.9; Jenkinson et al., 2002). BOLD runs were slice-time corrected to 0.696 s (0.5 of slice acquisition range 0–1.39 s) using `3dTshift` from AFNI 20160207 (Cox and Hyde, 1997; RRID: SCR_005927). The BOLD time series (including slice-timing correction when applied) were resampled onto their original native space by applying the transforms to correct for head motion. These resampled BOLD time series are referred to as preprocessed BOLD in original space, or simply preprocessed BOLD.

A T2* map was estimated from the preprocessed BOLD by fitting a monoexponential signal decay model with nonlinear regression, using T2*/S0 estimates from a log-linear regression fit as initial values. For each voxel, the maximal number of echoes with reliable signal in that voxel were used to fit the model. The calculated T2* map was then used to optimally combine preprocessed BOLD across echoes following the method described by Posse et al., 1999. The optimally combined time series was carried forward as the preprocessed BOLD. The BOLD time series were subsequently resampled into standard space, generating a preprocessed BOLD run in MNI152NLin2009cAsym space.

Several confounding time series were calculated based on the preprocessed BOLD: framewise displacement (FD), DVARS, and three region-wise global signals. FD was computed using two formulations—following Power (absolute sum of relative motions; Power et al., 2014) and Jenkinson (relative root mean square displacement between affines; Jenkinson et al., 2002). FD and DVARS were calculated for each functional run using their Nipype implementations (following the definitions by Power et al., 2014). The three global signals were extracted within the CSF, WM, and whole-brain masks. Additionally, physiological regressors were extracted to allow for component-based noise correction (CompCor; Behzadi et al., 2007). Principal components were estimated after high-pass filtering the preprocessed BOLD time series (discrete cosine filter with 128 s cutoff) for both temporal (tCompCor) and anatomical (aCompCor) variants. tCompCor components were calculated from the top 2% most variable voxels within the brain mask. For aCompCor, three probabilistic masks (CSF, WM, and combined CSF+WM) were generated in anatomical space. The implementation differs from Behzadi et al., 2007 in that, instead of eroding masks by two pixels in BOLD space, the aCompCor masks exclude voxels likely containing gray matter (GM) by subtracting a mask thresholded at 0.05 from the corresponding partial volume map, ensuring components are not extracted from voxels with minimal GM content. These masks were resampled into BOLD space and binarised by thresholding at 0.99, consistent with the original implementation. Components were also calculated separately within the WM and CSF masks. For each CompCor decomposition, the $k$ components with the largest singular values were retained such that the retained components explained at least 50% of the variance across the nuisance mask (CSF, WM, combined, or temporal). The remaining components were dropped.

Head-motion estimates from the correction step were also included in the corresponding confounds file. Confound time series derived from head motion estimates and global signals were expanded with temporal derivatives and quadratic terms for each (Satterthwaite et al., 2013). Frames exceeding 0.5 mm FD or 1.5 standardised DVARS were annotated as motion outliers. All resamplings were performed in a single interpolation step by composing the pertinent transformations (i.e., head-motion transforms, susceptibility distortion correction when available, and co-registrations to anatomical and output spaces). Gridded (volumetric) resamplings were performed using `antsApplyTransforms` (ANTs), configured with Lanczos interpolation to minimize smoothing effects Lanczos, 1964. Non-gridded (surface) resamplings were performed using `mri_vol2surf` (FreeSurfer).

Many internal operations of fMRIPrep use Nilearn 0.6.2 (Abraham et al., 2014; RRID: SCR_001362), primarily within the functional processing workflow. For more details on pipeline components, see the corresponding workflow sections in the fMRIPrep documentation.

**Copyright waiver.** The above boilerplate text was automatically generated by fMRIPrep with the express intention that users should copy and paste this text into their manuscripts unchanged. It is released under the CC0 license.

#### Schedules

**Table S1** | Intake session. S

| Time | Activity |
|---|---|
| 0:00 | Informed consent |
| 0:05 | Demographic variables |
| 0:10 | Medical screening |
| 0:15 | Hand dominance |

*Continued on next page*

Table S1 – *Continued from previous page*

| Time | Activity |
|---|---|
| 0:20 | MINI |
| 0:45 | HR/BP/Temp |
| 0:50 | ECG |
| 1:10 | Inclusion/exclusion questionnaire |
| 1:20 | Take to behavioural lab |
| 1:25 | EOG |
| 1:50 | Digit Span task |
| 2:00 | Take to cubicles |
| 2:05 | Operation Span task |
| 2:20 | WM Gating task practice |
| 2:35 | Perceptual decision-making task practice |
| 2:50 | Simon task practice |
| 3:00 | End of session, save data on P-drive |

*Note.* The intake session could take place at any time during work-time hours (08:00am-06:00pm).

MINI, Mini-International Neuropsychiatric Interview; HR, heart rate; BP, blood pressure; Temp, body temperature; ECG, electrocardiogram; EOG, electrooculogram; WM, working memory.

**Table S2** | Pharmacological session 1 & 2.

| Time | Activity |
|---|---|
| 9:00 | Medical Screening + Pregnancy test for women |
| 9:10 | VAS/PANAS |
| 9:15 | HR/BP/Temp |
| 9:20 | Give Capsule + Water |
| 9:20 | Bring to waiting room |
| | Make sure cups, water, magazines are available in waiting room |
| | Waiting Period (70 min) |
| 10:30 | Bring participant to MRI |

*Continued on next page*

Table S2 – *Continued from previous page*

| Time | Activity |
|---|---|
| 10:35 | MRI Screening + Check for metal |
| 10:45 | HR/BP/Temp |
| 10:50 | VAS/PANAS |
| 10:55 | (snack) |
| 11:00 | Put subject in scanner |
| 11:10 | Structural scan |
| 11:15 | Gating task (fMRI) |
| 12:15 | Get subject out of scanner |
| 12:20 | Clean/Save Data |
| 12:25 | Bring subject for lunch (ca. 25 min) |
| 12:50 | Bring subject to cubicles |
| 12:55 | Perceptual decision-making task (ca. 30 min) |
| 13:30 | Break (5 min) |
| 13:35 | Simon Task (ca. 30 min) |
| 14:10 | Digit span |
| 14:20 | VAS/PANAS |
| | HR/BP/Temp |
| 14:30 | Driving Assessment |
| 14:35 | Save data on P-drive |

*Note.* The start of the pharmacological session could be at 09:00, 09:45, 10:30 or 10:45. Both sessions of one participant were scheduled at the same time.

VAS, visual analogue scales; PANAS, positive and negative affect scale; HR, heart rate; BP, blood pressure; Temp, body temperature; MRI, magnetic resonance imaging.

#### Supplementary Results

**Control analyses.** The control analyses showed that changes from baseline to after the waiting period in physiological measures, subjective mood and medical questionnaire ratings did not differ significantly between sulpiride and placebo sessions (Pillai's Trace = $.15$, F(7, 60) = $1.54$, p = $.17$; Supplementary

Table S3). Also, cognitive performance on the Digit Span Task did not differ between the drug and placebo session within subjects ($B_{\text{drug-placebo}} = 0.76$ (SE $= 0.54$), p $= .156$). There were also no differential effects on the forward or backward versions of the Digit Span Task ($B_{\text{drug-placebo} \times \text{forward-backward}}$ $= 0.94$ (SE $= 1.07$), p $= .382$). Moreover, the James Blinding Index confirmed that blinding was successful (JBI $= .52(.42, .62)$), even though participants were slightly more likely to guess that they received the opposite agent. Therefore, any observed effects of sulpiride on BOLD signal and task performance cannot be attributed to physiological and subjective differences between the drug and placebo sessions.

Control analyses to check whether sulpiride acted on the striatum, show that BOLD signal is increased throughout the entire task in the striatum under sulpiride compared with placebo (sulpiride vs. placebo; small-volume correction (SVC) over the whole striatum; left caudate nucleus: $N_{\text{voxels}} = 48$, $x, y, z = (-12, 12, 12)$, Z $= 3.98$, $p_{\text{peak FWE SVC}} = .018$). No other areas in the brain showed significantly different BOLD signal between the drug and placebo sessions. Even though we cannot directly translate the direction of the BOLD effect into the direction of the drug effect, this suggests that sulpiride specifically affected BOLD signal in the striatum throughout the whole task, confirming the predicted and previously reported neural effects of this selective dopamine D2 receptor antagonist.

**Behavioural.** Across all participants and runs, mean accuracy was $M = 87.24\%$, $SD = 33.36\%$. The lowest individual accuracy was $M = 69.69\%$, $SD = 46.03\%$, and the highest was $M = 98.16\%$, $SD = 13.45\%$. Overall mean response time (RT) was $M = 967.89$ ms, $SD = 313.77$ ms. The fastest participant responded on average $M = 586.29$ ms, $SD = 141.86$ ms, and the slowest $M = 1207.01$ ms, $SD = 302.50$ ms.

Gating costs differed between input- and output-gating. The Bayesian regression model (BRMS) showed that accuracy was generally worse for output-gating trials than input-gating trials ($\text{trialType}_{\text{pre-retro}}$; accuracy: OR $= 1.28$, CI $= [1.12, 1.46]$). Also, the effect of gating (global 1 vs. gating) was greater on retro-cue trials than pre-cue trials ($\text{selectivity}_{\text{global1-gating}} \times \text{trialType}_{\text{pre-retro}}$; accuracy: OR $= 0.61$, CI $= [0.41, 0.89]$; RT: B $= 22.51$, CI $= [4.56, 40.18]$). A breakdown of this interaction effect revealed that the effect is driven by behavioural gating costs being evident for output-gating trials ($\text{selectivity}_{\text{global1-gating}}$; accuracy: OR $= 1.81$, CI $= [1.35, 2.48]$; RT: B $= 40.79$, CI $= [26.50, 54.96]$), but not for input-gating trials ($\text{selectivity}_{\text{global1-gating}}$; accuracy: OR $= 1.08$, CI $= [0.81, 1.44]$; RT: B $= 14.21$, CI $= [-1.06, 29.60]$). This suggests that output-gating required more cognitive effort than input-gating, though this may be confounded by load: on retro-cue trials, two items must be remembered until cue onset, whereas on global 1 trials only one item is retained. On pre-cue trials, however, only one item is encoded, so differences may arise already at encoding.

Sulpiride reduced the RT gating benefits on input-gating trials. A breakdown of this interaction (reported in the main text) between drug (sulpiride vs. placebo) and selectivity (global 1 vs. gating), using estimated marginal means, revealed that the effect of drug on RTs was numerically numerically different between global 1 and gating trials ($RT_{\text{sulpiride-placebo}}$; global 1: B $= -13.50$, CI $= [-40.70, 13.10]$; gating: B $= 11.70$, CI $= [-10.10, 35.20]$), such that the behavioural gating benefit was reduced under sulpiride by speeding responses on global 1 trials and slowing responses on gating trials. This pattern suggests that stimulus salience generally was enhanced under sulpiride.

Additional behavioural analyses including the delay periods as regressors indicate that a longer delay between the pre-cue and the stimulus for input-gating trials as well as a longer delay between the stimulus and the retro-cue for output-gating trials resulted in slower responses (RT: $B_{\text{delay1}} = 3.26$, $CI_{\text{delay1}} = [1.12, 5.38]$). The second delay period, just before responding, influenced participants' accuracy negatively (accuracy: $B_{\text{delay2}} = -0.09$, $CI_{\text{delay2}} = [-0.12, -0.05]$). These delay effects did not differ between sulpiride and placebo.

**Table S3** | Participant characteristics.

| **Measurement** - (time point 1–2) | **Drug** - Mean (Std. Error) | **Placebo** - Mean (Std. Error) |
|---|---:|---:|
| systolic blood pressure | 3.35 (1.19) | 1.32 (1.82) |
| diastolic blood pressure | 48.29 (1.56) | 47.09 (1.63) |
| heart rate | 10.79 (1.54) | 14.41 (1.58) |
| temperature | 0.04 (0.04) | -0.06 (0.06) |
| VAS mood | 0.10 (0.07) | -0.12 (0.05) |
| VAS medical | 0.06 (0.04) | 0.05 (0.04) |
| PANAS | 0.23 (0.04) | 0.15 (0.03) |

VAS, visual analogue scales; PANAS, positive and negative affect scale.

**Table S4** | Questionnaire results.

| **Measurement** | **Scale** | **Mean (Std. Error)** |
|---|---|---:|
| BDI | total | 4.74 (0.70) |
| BIS-11 | Attention | 9.02 (0.46) |
| BIS-11 | Cognitive_Complexity | 9.58 (0.41) |

*Continued on next page*

*Continued from previous page*

| **Measurement** | **Scale** | **Mean (Std. Error)** |
|---|---|---|
| BIS-11 | Cognitive_Instability | 6.08 (0.32) |
| BIS-11 | Motor | 13.31 (0.57) |
| BIS-11 | Perseverance | 6.60 (0.29) |
| BIS-11 | Second_Attentional_Impulsiveness | 15.10 (0.71) |
| BIS-11 | Second_Motor_Impulsiveness | 19.92 (0.74) |
| BIS-11 | Second_Nonplanning_Impulsiveness | 20.67 (0.84) |
| BIS-11 | Self_Control | 11.08 (0.52) |
| BIS/BAS | BAS_drive | 9.09 (0.39) |
| BIS/BAS | BAS_fun | 10.87 (0.34) |
| BIS/BAS | BAS_reward | 14.39 (0.54) |
| BIS/BAS | BIS | 19.28 (0.61) |
| CSS | Checking | 2.15 (0.39) |
| CSS | Contamination | 2.44 (0.38) |
| CSS | Danger | 4.04 (0.63) |
| CSS | Socio_Economic_Consequences | 0.56 (0.20) |
| CSS | Traumatic_Stress | 1.06 (0.29) |
| CSS | Xenophobia | 0.00 (0.00) |
| MBI | Cynicism | 12.00 (1.27) |
| MBI | Emotional_Exhaustion | 14.12 (1.28) |
| MBI | Personal_Efficacy | 24.39 (1.02) |
| STAI | State | 36.83 (1.51) |

BDI, Beck's Depression Inventory; BIS-11, Barratt Impulsiveness Scale; BIS/BAS, Behavioral Activation and Behavioral Inhibition Scale; CSS, Covid-19 Stress Scale; MBI, Maslach Burnout Inventory; STAI, State-Trait Anxiety Inventory.

**Bayesian model results.**

**Table S5** | Full reaction time model coefficients and 95% credible intervals, including the global 2 level of the variable selectivity: `RT ~ selectivity * trialType * drug + (1 + selectivity * trialType * drug | subject)`

| Coefficient | Estimate | Std. Error | Q2.5 | Q97.5 |
|---|---|---|---|---|
| Intercept | 989.04 | 20.05 | 949.78 | 1028.77 |
| selectivityGlo1vsGat | -25.93 | 5.95 | -37.55 | -14.10 |
| selectivityGlo2vsGat | 130.80 | 8.79 | 113.55 | 148.28 |
| trialTypeRetvsPre | -7.47 | 5.88 | -18.79 | 4.06 |
| drugSulvsPla | -5.21 | 9.97 | -25.00 | 14.22 |
| selectivityGlo1vsGat:trialTypeRetvsPre | -22.51 | 9.12 | -40.18 | -4.56 |
| selectivityGlo1vsGat:drugSulvsPla | -4.11 | 9.96 | -23.74 | 15.75 |
| selectivityGlo2vsGat:trialTypeRetvsPre | -7.47 | 10.26 | -28.15 | 12.90 |
| selectivityGlo2vsGat:drugSulvsPla | -21.51 | 9.96 | -39.24 | -3.28 |
| trialTypeRetvsPre:drugSulvsPla | 0.76 | 8.25 | -15.81 | 16.67 |
| selectivityGlo1vsGat:trialTypeRetvsPre:drugSulvsPla | 36.99 | 17.21 | 3.45 | 70.38 |
| selectivityGlo2vsGat:trialTypeRetvsPre:drugSulvsPla | -1.23 | 18.62 | -37.99 | 35.27 |

**Table S6** | Full accuracy model coefficients and 95% credible intervals, including the global 2 level of the variable selectivity: `accuracy ~ selectivity * trialType * drug + (1 + selectivity * trialType * drug | subject)`

| Coefficient | Estimate | Std. Error | Q2.5 | Q97.5 |
|---|---|---|---|---|
| Intercept | 2.12 | 0.12 | 1.89 | 2.35 |
| selectivityGlo1vsGat | 0.34 | 0.09 | 0.16 | 0.53 |
| selectivityGlo2vsGat | -0.86 | 0.10 | -1.07 | -0.64 |
| trialTypeRetvsPre | -0.25 | 0.07 | -0.38 | -0.11 |
| drugSulvsPla | -0.13 | 0.11 | -0.34 | 0.08 |
| selectivityGlo1vsGat:trialTypeRetvsPre | 0.50 | 0.20 | 0.12 | 0.88 |
| selectivityGlo2vsGat:trialTypeRetvsPre | 0.30 | 0.17 | -0.03 | 0.63 |
| selectivityGlo1vsGat:drugSulvsPla | 0.17 | 0.17 | -0.18 | 0.50 |
| selectivityGlo2vsGat:drugSulvsPla | 0.11 | 0.14 | -0.16 | 0.38 |
| trialTypeRetvsPre:drugSulvsPla | 0.10 | 0.15 | -0.22 | 0.38 |
| selectivityGlo1vsGat:trialTypeRetvsPre:drugSulvsPla | -0.28 | 0.35 | -0.96 | 0.42 |

*Continued on next page*

*Continued from previous page*

| Coefficient | Estimate | Std. Error | Q2.5 | Q97.5 |
|---|---|---|---|---|
| selectivityGlo2vsGat:trialTypeRetvsPre:drugSulvsPla | -0.23 | 0.28 | -0.79 | 0.34 |

**Table S7** | Model coefficients and 95% credible intervals for **reaction time** model with stimulus category for **input-gating** trials, excluding the global 2 level of the variable selectivity: `RT ~ selectivity * stimulusCategory * drug + (1 + selectivity * stimulusCategory * drug | subject)`

| Coefficient | Estimate | Std. Error | Q2.5 | Q97.5 |
|---|---|---|---|---|
| Intercept | 933.36 | 19.82 | 894.08 | 971.44 |
| selectivityGlo1vsGat | 14.21 | 7.88 | -1.06 | 29.60 |
| drugSulvsPla | -0.95 | 11.08 | -23.06 | 21.37 |
| stimFacvsSce | -19.56 | 7.79 | -35.21 | -4.17 |
| selectivityGlo1vsGat:drugSulvsPla | 24.93 | 12.24 | 0.76 | 48.93 |
| selectivityGlo1vsGat:stimFacvsSce | -15.96 | 12.97 | -41.34 | 9.53 |
| drugSulvsPla:stimFacvsSce | -26.55 | 11.47 | -49.12 | -4.12 |
| selectivityGlo1vsGat:drugSulvsPla:stimFacvsSce | -21.19 | 23.81 | -68.42 | 25.41 |

**Table S8** | Model coefficients and 95% credible intervals for **accuracy** model with stimulus category for **input-gating** trials, excluding the global 2 level of the variable selectivity: `accuracy ~ selectivity * stimulusCategory * drug + (1 + selectivity * stimulusCategory * drug | subject)`

| Coefficient | Estimate | Std. Error | Q2.5 | Q97.5 |
|---|---|---|---|---|
| Intercept | 2.73 | 0.16 | 2.42 | 3.07 |
| selectivityGlo1vsGat | 0.08 | 0.15 | -0.21 | 0.36 |
| drugSulvsPla | -0.19 | 0.19 | -0.55 | 0.19 |
| stimFacvsSce | 0.39 | 0.17 | 0.03 | 0.73 |
| selectivityGlo1vsGat:drugSulvsPla | 0.36 | 0.26 | -0.14 | 0.89 |
| selectivityGlo1vsGat:stimFacvsSce | -0.21 | 0.31 | -0.83 | 0.40 |
| drugSulvsPla:stimFacvsSce | 0.46 | 0.27 | -0.09 | 1.00 |
| selectivityGlo1vsGat:drugSulvsPla:stimFacvsSce | 0.12 | 0.59 | -1.05 | 1.27 |

**Table S9** | Model coefficients and 95% credible intervals for **reaction time** model with stimulus category for **output-gating** trials, excluding the global 2 level of the variable selectivity: `RT ~ selectivity * stimulusCategory * drug + (1 + selectivity * stimulusCategory * drug | subject)`

| Coefficient | Estimate | Std. Error | Q2.5 | Q97.5 |
|---|---|---|---|---|
| Intercept | 936.01 | 18.16 | 900.68 | 971.65 |
| selectivityGlo1vsSel | 40.79 | 7.17 | 26.50 | 54.96 |
| drugSulvsPla | 5.98 | 9.43 | -12.77 | 24.89 |
| stimFacvsSce | -25.62 | 7.92 | -41.06 | -10.03 |
| selectivityGlo1vsSel:drugSulvsPla | -12.13 | 13.01 | -37.40 | 13.33 |
| selectivityGlo1vsSel:stimFacvsSce | -43.35 | 12.91 | -68.76 | -18.03 |
| drugSulvsPla:stimFacvsSce | -10.40 | 12.11 | -34.46 | 13.04 |
| selectivityGlo1vsSel:drugSulvsPla:stimFacvsSce | -16.21 | 23.07 | -61.23 | 29.70 |

**Table S10** | Model coefficients and 95% credible intervals for **accuracy model** with stimulus category for **output-gating** trials, excluding the global 2 level of the variable selectivity: `accuracy ~ selectivity * stimulusCategory * drug + (1 + selectivity * stimulusCategory * drug | subject)`

| Coefficient | Estimate | Std. Error | Q2.5 | Q97.5 |
|---|---|---|---|---|
| Intercept | 2.35 | 0.12 | 2.13 | 2.59 |
| selectivityGlo1vsGat | 0.59 | 0.15 | 0.30 | 0.91 |
| drugSulvsPla | -0.06 | 0.14 | -0.33 | 0.22 |
| stimFacvsSce | 0.46 | 0.12 | 0.21 | 0.70 |
| selectivityGlo1vsGat:drugSulvsPla | 0.03 | 0.24 | -0.44 | 0.50 |
| selectivityGlo1vsGat:stimFacvsSce | -0.10 | 0.24 | -0.57 | 0.37 |
| drugSulvsPla:stimFacvsSce | 0.26 | 0.26 | -0.24 | 0.78 |
| selectivityGlo1vsGat:drugSulvsPla:stimFacvsSce | 0.18 | 0.48 | -0.76 | 1.11 |

**fMRI.**

**Sulpiride increased striatal BOLD signal in the caudate nucleus during encoding.** Striatal BOLD signal was increased by sulpiride during encoding. Specifically, the drug enhanced BOLD signal in the caudate nucleus bilaterally and in the putamen unilaterally during stimulus encoding, regardless of whether participants had to input-gate stimuli into their WM or just encode all present stimuli into their WM (right caudate nucleus: $N_{\text{voxels}} = 111$, $x, y, z = (11, -2, 16)$, $Z = 5.11$, $p_{\text{peak FWE SVC}} < .001$; left caudate nucleus: $N_{\text{voxels}} = 24$, $x, y, z = (-12, 15, 9)$, $Z = 4.01$, $p_{\text{peak FWE SVC}} = .014$; right putamen:

$N_{\text{voxels}} = 23$, $x, y, z = (24, -10, -4)$, $Z = 3.89$, $p_{\text{peak FWE SVC}} = .023$ (Figure S8; Supplementary Table S17). Contrary to the hypothesis that striatal dopamine modulates gating specifically, the effect of sulpiride on striatal BOLD signal (or elsewhere in the brain; Supplementary Table S18) did not differ between gating and global 1 trials at the time of encoding (sample), suggesting that sulpiride had a nonspecific effect on stimulus detection and not on WM gating.

**Supplementary motor cortex responded to global encoding load.** Higher load during global encoding was associated with enhanced BOLD signal in the supplementary motor area (SMA) (global 2 vs. global 1; right SMA: $N_{\text{voxels}} = 356$, $x, y, z = (1, 12, 49)$, $Z = 7.174$, $p_{\text{peak FWE}} < .001$), the insula (right insula: $N_{\text{voxels}} = 68$, $x, y, z = (36, 18, 6)$, $Z = 5.72$, $p_{\text{peak FWE}} < .001$; left insula: $N_{\text{voxels}} = 11$, $x, y, z = (-42, 15, 4)$, $Z = 5.05$, $p_{\text{peak FWE}} = .007$), and the left lingual gyrus (left lingual gyrus: $N_{\text{voxels}} = 18$, $x, y, z = (-2, -78, -4)$, $Z = 5.64$, $p_{\text{peak FWE}} < .001$) (Figure S9; Supplementary Table S20). By contrast, there were no effects of load (global 2 vs. global 1) at the retro-cue. Additionally, there were no effects of sulpiride on load-related frontal BOLD signal during input- or output-gating.

**Increases and decreases in striatal BOLD signal during WM input- and output-gating for placebo session only.** A replication of the analyses to validate the experimental manipulations on the placebo data only showed no differences to the analyses including both the drug and placebo data. For example, stimulus-selectivity in visual association cortex remained significant for input-gating ($\text{area}_{\text{FFA-PPA}} \times \text{category}_{\text{face-scene}} \times \text{selectivity}_{\text{global1-gating}}$: $B = -9.19$, $CI = [-13.96, -4.44]$) and output-gating ($\text{area}_{\text{FFA-PPA}} \times \text{category}_{\text{face-scene}} \times \text{selectivity}_{\text{global1-gating}}$: $B = -12.50$, $CI = [-16.29, -8.74]$). Also, striatal BOLD signal increases during input-gating (gating vs. global 1; small-volume correction (SVC)

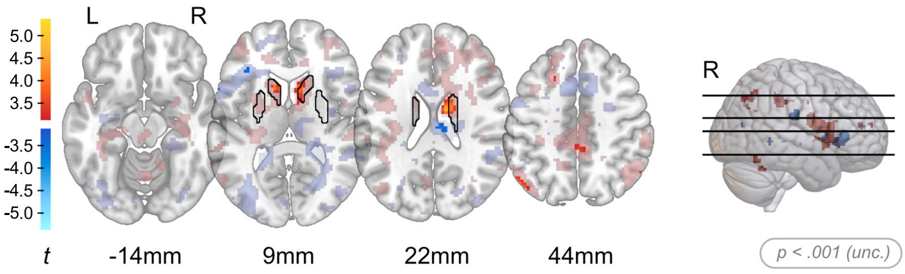


**Figure S8 | Sulpiride increased striatal BOLD signal during encoding (across gating and global trials).** Whole brain contrasts showing BOLD signal at p < .001 unc. in full opacity (sulpiride > placebo at encoding). BOLD signal increases compared with baseline are shown in warm colours while BOLD signal decreases are shown in cold colours. The striatal mask used for small volume correction is outlined in black. In addition, low-intensity, transparent colours present voxels below threshold (p < .1 unc. whole brain) for visualisation purposes.

L, left hemisphere; R, right hemisphere; unc, uncorrected.

over the whole striatum; left caudate: $N_{\text{voxels}} = 24$, $x, y, z = (-16, 15, 6)$, Z $= 4.22$, $p_{\text{peak FWE SVC}} = .007$) and decreases during output-gating (gating vs. global 1; SVC over the whole striatum; right putamen: $N_{\text{voxels}} = 308$, $x, y, z = (24, 10, 9)$, Z $= -\infty$, $p_{\text{peak FWE SVC}} < .001$; right caudate nucleus: $N_{\text{voxels}} = 95$, $x, y, z = (18, 12, 12)$, Z $= -7.57$, $p_{\text{peak FWE SVC}} < .001$; left putamen: $N_{\text{voxels}} = 275$, $x, y, z = (-24, 8, 6)$, Z $= -6.74$, $p_{\text{peak FWE SVC}} < .001$) were replicated in the model with only the placebo data).

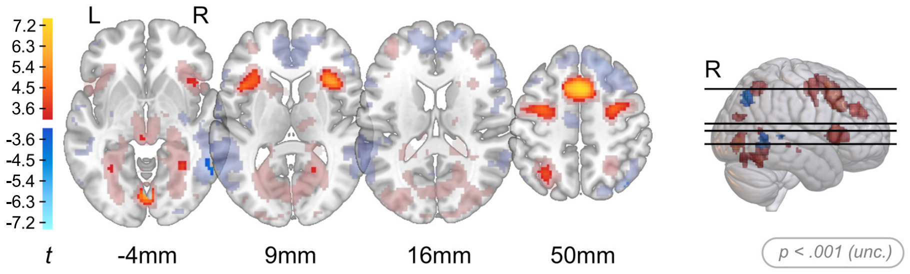


**Figure S9 | Supplementary motor cortex responded to global encoding load.** Whole brain contrasts showing BOLD signal at p $<$ .001 uncorrected in full opacity (global 2 $>$ global 1 at encoding). BOLD signal increases compared with baseline are shown in warm colours while BOLD signal decreases are shown in cold colours. In addition, low-intensity, transparent colours present voxels below threshold (p $<$ .1 unc. whole brain) for visualisation purposes.

L, left hemisphere; R, right hemisphere; unc, uncorrected.

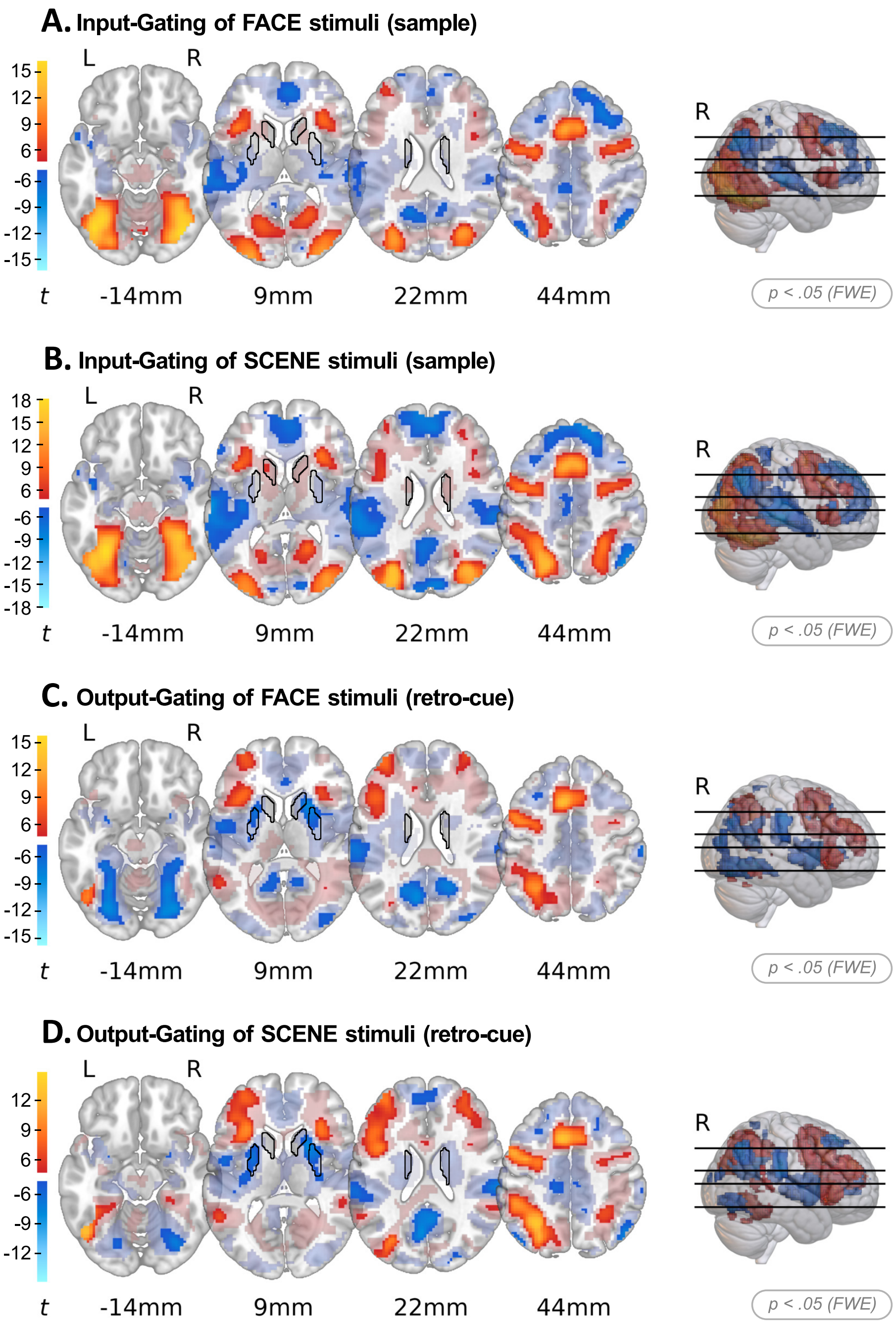


**Figure S10 | Input- and output-gating of scene and face stimuli activate the fronto-parietal control network.** Whole brain contrasts showing BOLD signal at $p < .05$ FWE corrected in full opacity. BOLD signal increases compared with baseline are shown in warm colours while BOLD signal decreases are shown in cold colours. The striatal mask used for small volume correction is outlined in black. In addition, low-intensity, transparent colours present voxels below threshold ($p < .1$ unc. whole brain) for visualisation purposes. **A.** Input-gating of face stimuli (face gating > face global 1 at sample) **B.** Input-gating of scene stimuli (scene gating > scene global 1 at sample) **C.** Output-gating of face stimuli (face gating > face global 1 at retro-cue) **D.** Output-gating of scene stimuli (scene gating > scene global 1 at retro-cue).

L, left hemisphere; R, right hemisphere; FWE, family-wise error; unc, uncorrected.

#### Discussion

Contrary to our prediction, sulpiride increased striatal BOLD signal during encoding in a manner that did not depend on the pre-cue that was presented. Thus, the sulpiride-induced increase in striatal signal during encoding did not differ between the 'face', 'scene', or 'both' pre-cues. This indicates that the effect is unlikely to reflect a modulation of WM gating specifically, a conclusion that is supported by the lack of an effect of sulpiride on gating gain effects in visual association cortex during input-gating. At the same time, sulpiride speeded behavioural responding on global 1 trials while slowing it on input-gating trials. It is possible that this pattern of behavioural effects as a function of input-gating reflects sulpiride-induced increases in nonspecific stimulus salience, in line with the nonspecific neural effect of sulpiride during encoding. Indeed, enhanced stimulus salience might have increased competition between the two stimuli presented on the screen during encoding on gating trials, while facilitating responding on global 1 trials on which only 1 stimulus was presented on the screen during encoding. Thus, instead of reflecting modulation of WM gating, the effect might reflect modulation of a nonspecific salience, stimulus detection or warning signal (Redgrave et al., 1999).

The dissociation between behavioural and neural effects, with sulpiride altering neural activity as a function of output-gating but not input-gating, suggests that the mechanisms of input-gating for encoding and maintenance may differ from those involved in the output-gating for retrieval. Consequently, dopaminergic effects may differentially manifest across behavioural and neural measures for these processes. One possible explanation is that input-gating relies more strongly on perceptual filtering processes that are primarily reflected in behaviour, whereas output-gating may depend more on executive control mechanisms that are more directly captured at a neural level.

The effects as a function of cognitive load suggest that WM input- and output-gating operations are distinct from those recruited as a function of encoding and retrieval load. Consistent with previous work showing that the SMA and insula are engaged under higher WM demands (Miri Ashtiani and Daliri, 2023), BOLD signal in these regions was increased at higher encoding load, for both input- and output-gating. Activity in other brain areas, including critically the striatum, was not modulated by load, either at encoding or retrieval. This finding suggests that the effect of sulpiride during WM gating, which we interpret to reflect changes in the effort costs of cognitive control, are unlikely to reflect changes in load per se.

### Tables

**Table S11** | Pre-cue > baseline.

| Direction | Hemisphere | MNI Coordinates | | | Peak | | Cluster | | | Peak (SVC) | Region |
|---|---|---|---|---|---|---|---|---|---|---|---|
| | | x | y | z | Z | p(Z) *unc.* | k | p(k) *unc.* | p(k) *FWE* | p(Z) *FWE* | |
| + | R | 13.5 | -70 | -3.5 | Inf | 0 | 1913 | 0 | 0 | | Lingual R, BA 18, Lingual Gyrus |
| + | R | 3.5 | -67.5 | -11 | Inf | 0 | | | | | Vermis 6, Declive |
| + | R | 3.5 | -50 | -13.5 | Inf | 0 | | | | | Vermis 4 5, Culmen |
| + | R | 11 | -7.5 | 34 | 6.513 | 0 | 1930 | 0 | 0 | | Cingulate Mid R, Cingulate Gyrus |
| + | R | 28.5 | 12.5 | 16.5 | 5.797 | 0 | | | | | Extra-Nuclear |
| + | R | 1 | 17.5 | 29 | 5.754 | 0 | | | | | Cingulate Ant R |
| + | R | 11 | -37.5 | 51.5 | 4.516 | 0 | 166 | 0.001 | 0.006 | | Precuneus R, BA 5, Paracentral Lobule |
| + | R | 11 | -47.5 | 44 | 3.753 | 0 | | | | | Precuneus R, BA 7, Precuneus |
| + | R | 61 | -10 | 4 | 3.532 | 0 | 9 | 0.365 | 0.924 | | Temporal Sup R, BA 22, Superior Temporal Gyrus |

*Continued on next page*

Table S11 – *Continued from previous page*

| Direction | Hemisphere | MNI Coordinates | | | Peak | | Cluster | | | Peak (SVC) | Region |
|---|---|---|---|---|---|---|---|---|---|---|---|
| | | x | y | z | Z | p(Z) *unc.* | k | p(k) *unc.* | p(k) *FWE* | p(Z) *FWE* | |
| + | L | -14 | -12.5 | 51.5 | 3.349 | 0 | 6 | 0.464 | 0.962 | | BA 6, Medial Frontal Gyrus |
| + | L | -9 | -15 | 36.5 | 3.27 | 0.001 | 1 | 0.792 | 0.996 | | Cingulate Mid L, Cingulate Gyrus |
| + | R | 46 | 0 | 26.5 | 3.257 | 0.001 | 7 | 0.427 | 0.951 | | BA 6, Precentral Gyrus |
| + | R | 38.5 | -10 | 29 | 3.128 | 0.001 | 7 | 0.427 | 0.951 | | Precentral Gyrus |
| + | R | 11 | -52.5 | 24 | 3.097 | 0.001 | 1 | 0.792 | 0.996 | | Precuneus R, Posterior Cingulate |
| - | R | 48.5 | -20 | 6.5 | 6.871 | 0 | 1707 | 0 | 0 | | Heschl R, Superior Temporal Gyrus |
| - | R | 61 | -22.5 | 6.5 | 6.43 | 0 | | | | | Temporal Sup R, Superior Temporal Gyrus |
| - | R | 41 | -25 | 11.5 | 6.379 | 0 | | | | | Heschl R, BA 41, Transverse Temporal Gyrus |
| - | L | -41.5 | -30 | 11.5 | 6.746 | 0 | 13911 | 0 | 0 | | Temporal Sup L, BA 41, Transverse Temporal Gyrus |

*Continued on next page*

Table S11 – *Continued from previous page*

| Direction | Hemisphere | MNI Coordinates | | | Peak | | Cluster | | | Peak (SVC) | Region |
|---|---|---|---|---|---|---|---|---|---|---|---|
| | | x | y | z | Z | p(Z) *unc.* | k | p(k) *unc.* | p(k) *FWE* | p(Z) *FWE* | |
| - | L | -46.5 | -37.5 | 14 | 6.725 | 0 | | | | | Temporal Sup L, Superior Temporal Gyrus |
| - | L | -49 | 15 | -13.5 | 6.689 | 0 | | | | | Temporal Pole Sup L, BA 38, Superior Temporal Gyrus |
| - | L | -34 | -60 | 24 | 4.889 | 0 | 40 | 0.066 | 0.372 | | Middle Temporal Gyrus |
| - | R | 43.5 | -77.5 | 29 | 4.455 | 0 | 38 | 0.072 | 0.399 | | Occipital Mid R, BA 19, Superior Occipital Gyrus |
| - | L | -41.5 | -80 | 31.5 | 4.253 | 0 | 22 | 0.161 | 0.679 | | Occipital Mid L |
| - | L | -4 | -22.5 | -23.5 | 3.844 | 0 | 13 | 0.276 | 0.858 | | |
| - | R | 28.5 | -17.5 | 61.5 | 3.772 | 0 | 15 | 0.243 | 0.82 | | Precentral R, Precentral Gyrus |
| - | L | -16.5 | -52.5 | 21.5 | 3.483 | 0 | 3 | 0.616 | 0.987 | | Sub-Gyral |
| - | R | 8.5 | -20 | -1 | 3.477 | 0 | 12 | 0.295 | 0.876 | | Thalamus R |
| - | R | 33.5 | -12.5 | 24 | 3.337 | 0 | 5 | 0.506 | 0.972 | | Extra-Nuclear |
| - | R | 11 | -12.5 | -6 | 3.299 | 0 | 1 | 0.792 | 0.996 | | |
| - | L | -11.5 | -25 | 1.5 | 3.185 | 0.001 | 1 | 0.792 | 0.996 | | Thalamus L, Thalamus |

*Continued on next page*

| **Direction** | **Hemisphere** | **MNI Coordinates** | | | **Peak** | | **Cluster** | | | **Peak (SVC)** | **Region** |
|---|---|---|---|---|---|---|---|---|---|---|---|
| | | x | y | z | Z | p(Z) *unc.* | k | p(k) *unc.* | p(k) *FWE* | p(Z) *FWE* | |
| - | L | -11.5 | 52.5 | 31.5 | 3.171 | 0.001 | 3 | 0.616 | 0.987 | | Frontal Sup 2 L, Superior Frontal Gyrus |
| - | L | -21.5 | 0 | 39 | 3.164 | 0.001 | 3 | 0.616 | 0.987 | | Sub-Gyral |
| - | R | 31 | -17.5 | 39 | 3.14 | 0.001 | 1 | 0.792 | 0.996 | | Sub-Gyral |
| + | R | 16 | 22.5 | 6.5 | 5.134 | 0 | 89 | 0.01 | 0.003 | 0 | Caudate R, Extra-Nuclear |
| + | R | 11 | 5 | 6.5 | 3.217 | 0.001 | | | | 0.178 | Caudate R, Caudate Body, Caudate |
| + | R | 18.5 | 0 | 24 | 4.952 | 0 | 45 | 0.053 | 0.014 | 0 | Caudate R, Extra-Nuclear |
| + | R | 18.5 | -7.5 | 26.5 | 4.445 | 0 | | | | 0.003 | Cingulate Gyrus |
| + | R | 16 | -15 | 24 | 4.139 | 0 | | | | 0.01 | Caudate Body, Caudate |
| + | L | -16.5 | 0 | 24 | 3.12 | 0.001 | 2 | 0.691 | 0.174 | 0.226 | Caudate L, Lateral Ventricle |

**Table S12** | Sample > baseline (Encoding).

| Direction | Hemisphere | MNI Coordinates | | | Peak | | Cluster | | | Peak (SVC) | Region |
|---|---|---|---|---|---|---|---|---|---|---|---|
| | | x | y | z | Z | p(Z) *unc.* | k | p(k) *unc.* | p(k) *FWE* | p(Z) *FWE* | |
| + | R | 3.5 | -42.5 | -11 | Inf | 0 | 5619 | 0 | 0 | | Vermis 3, Cerebellar Lingual |
| + | R | 6 | -25 | -21 | Inf | 0 | | | | | |
| + | R | 13.5 | -65 | 14 | Inf | 0 | | | | | Calcarine R, BA 31, Posterior Cingulate |
| + | R | 31 | -55 | -21 | Inf | 0 | 102 | 0.004 | 0.031 | | Cerebelum 6 R, Culmen |
| + | R | 36 | 27.5 | 49 | 7.48 | 0 | 1785 | 0 | 0 | | Frontal Mid 2 R, Superior Frontal Gyrus |
| + | R | 3.5 | 37.5 | 6.5 | 6.76 | 0 | | | | | Cingulate Ant R, BA 24, Anterior Cingulate |
| + | R | 1 | 15 | 41.5 | 6.693 | 0 | | | | | Frontal Sup Medial L, Cingulate Gyrus |
| + | R | 48.5 | -12.5 | 26.5 | 4.86 | 0 | 61 | 0.019 | 0.144 | | Precentral Gyrus |
| + | R | 23.5 | 35 | 6.5 | 4.758 | 0 | 118 | 0.002 | 0.017 | | Sub-Gyral |
| + | R | 26 | 42.5 | 4 | 4.568 | 0 | | | | | Sub-Gyral |
| + | R | 18.5 | -5 | 29 | 4.221 | 0 | 50 | 0.032 | 0.224 | | Cingulate Gyrus |
| + | R | 21 | -15 | 26.5 | 3.54 | 0 | | | | | Sub-Gyral |

*Continued on next page*

Table S12 – *Continued from previous page*

| Direction | Hemisphere | MNI Coordinates | | | Peak | | Cluster | | | Peak (SVC) | Region |
|---|---|---|---|---|---|---|---|---|---|---|---|
| | | x | y | z | Z | p(Z) *unc.* | k | p(k) *unc.* | p(k) *FWE* | p(Z) *FWE* | |
| + | R | 33.5 | -15 | 26.5 | 3.568 | 0 | 10 | 0.307 | 0.914 | | Sub-Gyral |
| + | R | 43.5 | 40 | 6.5 | 3.549 | 0 | 10 | 0.307 | 0.914 | | Frontal Mid 2 R, BA 46, Inferior Frontal Gyrus |
| + | R | 48.5 | -67.5 | -6 | 3.479 | 0 | 1 | 0.776 | 0.998 | | Temporal Inf R, Middle Occipital Gyrus |
| + | R | 16 | -25 | 19 | 3.401 | 0 | 8 | 0.362 | 0.944 | | Lateral Ventricle |
| + | R | 63.5 | -15 | 24 | 3.285 | 0.001 | 2 | 0.668 | 0.995 | | SupraMarginal R, Post-central Gyrus |
| + | R | 41 | -60 | -3.5 | 3.174 | 0.001 | 1 | 0.776 | 0.998 | | Sub-Gyral |
| - | L | -46.5 | -15 | -1 | Inf | 0 | 25232 | 0 | 0 | | Temporal Sup L, BA 22, Superior Temporal Gyrus |
| - | L | -34 | -17.5 | -1 | Inf | 0 | | | | | Extra-Nuclear |
| - | R | 38.5 | -17.5 | -1 | Inf | 0 | | | | | Insula R, Extra-Nuclear |
| - | R | 8.5 | 12.5 | 1.5 | 4.021 | 0 | 8 | 0.362 | 0.944 | | Caudate R, Caudate Head, Caudate |
| - | L | -31.5 | -72.5 | 4 | 3.72 | 0 | 2 | 0.668 | 0.995 | | Middle Occipital Gyrus |

*Continued on next page*

Table S12 – *Continued from previous page*

| Direction | Hemisphere | MNI Coordinates | | | Peak | | Cluster | | | Peak (SVC) | Region |
|---|---|---|---|---|---|---|---|---|---|---|---|
| | | x | y | z | Z | p(Z) *unc.* | k | p(k) *unc.* | p(k) *FWE* | p(Z) *FWE* | |
| - | L | -6.5 | 12.5 | 1.5 | 3.677 | 0 | 4 | 0.527 | 0.985 | | Caudate L, Caudate Head, Caudate |
| - | L | -9 | -60 | -41 | 3.664 | 0 | 3 | 0.589 | 0.991 | | Cerebelum 8 L, Inferior Semi-Lunar Lobule |
| - | R | 11 | -2.5 | 1.5 | 3.592 | 0 | 7 | 0.395 | 0.957 | | Medial Globus Pallidus, Lentiform Nucleus |
| - | R | 1 | -10 | 21.5 | 3.411 | 0 | 5 | 0.476 | 0.978 | | Corpus Callosum |
| - | R | 43.5 | 42.5 | 24 | 3.297 | 0 | 2 | 0.668 | 0.995 | | Frontal Mid 2 R, Middle Frontal Gyrus |
| - | L | -4 | -15 | 9 | 3.251 | 0.001 | 2 | 0.668 | 0.995 | | Thalamus L, Medial Dorsal Nucleus, Thalamus |
| - | R | 18.5 | -72.5 | -26 | 3.157 | 0.001 | 1 | 0.776 | 0.998 | | Cerebelum 6 R, Uvula |
| + | R | 31 | -12.5 | -3.5 | 5.857 | 0 | 30 | 0.086 | 0.026 | 0 | Pallidum R, Extra-Nuclear |
| + | R | 18.5 | -15 | 24 | 3.457 | 0 | 6 | 0.432 | 0.123 | 0.101 | Caudate R, Caudate Body, Caudate |
| - | L | -31.5 | -15 | 1.5 | Inf | 0 | 208 | 0 | 0 | 0 | Extra-Nuclear |

*Continued on next page*

Table S12 – *Continued from previous page*

| Direction | Hemisphere | MNI Coordinates | | | Peak | | Cluster | | | Peak (SVC) | Region |
|---|---|---|---|---|---|---|---|---|---|---|---|
| | | x | y | z | Z | p(Z) *unc.* | k | p(k) *unc.* | p(k) *FWE* | p(Z) *FWE* | |
| - | L | -26.5 | 10 | -8.5 | 5.607 | 0 | | | | 0 | Putamen L, Extra-Nuclear |
| - | R | 28.5 | -10 | 1.5 | 7.632 | 0 | 245 | 0 | 0 | 0 | Putamen R, Putamen, Lentiform Nucleus |
| - | R | 26 | 10 | -8.5 | 5.543 | 0 | | | | 0 | Putamen R, Extra-Nuclear |
| - | R | 8.5 | 12.5 | 1.5 | 4.021 | 0 | 7 | 0.395 | 0.113 | 0.016 | Caudate R, Caudate Head, Caudate |
| - | R | 11 | 15 | -1 | 3.932 | 0 | | | | 0.022 | Caudate R, Caudate Head, Caudate |
| - | L | -6.5 | 12.5 | 1.5 | 3.677 | 0 | 4 | 0.527 | 0.147 | 0.052 | Caudate L, Caudate Head, Caudate |

**Table S13** | Retro-cue > baseline (Retrieval).

| Direction | Hemisphere | MNI Coordinates | | | Peak | | Cluster | | | Peak (SVC) | Region |
|---|---|---|---|---|---|---|---|---|---|---|---|
| | | x | y | z | Z | p(Z) *unc.* | k | p(k) *unc.* | p(k) *FWE* | p(Z) *FWE* | |
| + | R | 11 | -40 | 44 | Inf | 0 | 2042 | 0 | 0 | | Cingulate Mid R, Precuneus |
| + | R | 8.5 | -47.5 | 36.5 | 7.369 | 0 | | | | | Cingulate Mid R, BA 31, Precuneus |
| + | L | -1.5 | -30 | 36.5 | 6.221 | 0 | | | | | Cingulate Mid L, BA 31, Cingulate Gyrus |
| + | R | 3.5 | 17.5 | 41.5 | 7.654 | 0 | 3704 | 0 | 0 | | Cingulate Mid R, BA 32, Cingulate Gyrus |
| + | R | 8.5 | 47.5 | 1.5 | 7.409 | 0 | | | | | Frontal Sup Medial R, Medial Frontal Gyrus |
| + | L | -6.5 | 42.5 | 26.5 | 7.183 | 0 | | | | | Frontal Sup Medial L, Medial Frontal Gyrus |
| + | L | -4 | -40 | -13.5 | 7.002 | 0 | 1096 | 0 | 0 | | Cerebelum 3 L, Culmen |
| + | L | -16.5 | -27.5 | 9 | 6.686 | 0 | | | | | Thalamus L, Pulvinar, Thalamus |
| + | L | -14 | -15 | -1 | 6.472 | 0 | | | | | Thalamus L |

*Continued on next page*

| **Direction** | **Hemisphere** | **MNI Coordinates** | | | **Peak** | | **Cluster** | | | **Peak (SVC)** | **Region** |
|---|---|---|---|---|---|---|---|---|---|---|---|
| | | x | y | z | Z | p(Z) *unc.* | k | p(k) *unc.* | p(k) *FWE* | p(Z) *FWE* | |
| + | R | 36 | -30 | -16 | 5.591 | 0 | 63 | 0.024 | 0.157 | | ParaHippocampal R, BA 36, Parahippocampal Gyrus |
| + | R | 31 | -52.5 | -23.5 | 5.24 | 0 | 37 | 0.072 | 0.407 | | Cerebelum 6 R, Culmen |
| + | R | 63.5 | -30 | 39 | 5.171 | 0 | 15 | 0.237 | 0.82 | | SupraMarginal R |
| + | L | -19 | 10 | -3.5 | 4.717 | 0 | 6 | 0.457 | 0.964 | | Putamen L, Putamen, Lentiform Nucleus |
| + | R | 38.5 | -15 | 21.5 | 4.671 | 0 | 142 | 0.002 | 0.011 | | Insula R, Extra-Nuclear |
| + | R | 48.5 | -17.5 | 24 | 4.568 | 0 | | | | | Postcentral Gyrus |
| + | R | 31 | -2.5 | -1 | 4.507 | 0 | 23 | 0.147 | 0.656 | | Putamen R, Extra-Nuclear |
| + | R | 51 | 0 | -1 | 4.27 | 0 | 26 | 0.125 | 0.596 | | Insula R, Superior Temporal Gyrus |
| + | R | 63.5 | -25 | 4 | 3.589 | 0 | 3 | 0.611 | 0.988 | | Temporal Sup R, BA 22, Superior Temporal Gyrus |
| + | R | 8.5 | 57.5 | 34 | 3.294 | 0 | 5 | 0.5 | 0.973 | | Frontal Sup Medial R, Superior Frontal Gyrus |

*Continued on next page*

Table S13 – *Continued from previous page*

| Direction | Hemisphere | MNI Coordinates | | | Peak | | Cluster | | | Peak (SVC) | Region |
|---|---|---|---|---|---|---|---|---|---|---|---|
| | | x | y | z | Z | p(Z) *unc.* | k | p(k) *unc.* | p(k) *FWE* | p(Z) *FWE* | |
| + | R | 21 | -20 | 1.5 | 3.24 | 0.001 | 3 | 0.611 | 0.988 | | Thalamus R, Extra-Nuclear |
| - | L | -24 | -62.5 | -16 | Inf | 0 | 9929 | 0 | 0 | | Fusiform L, Declive |
| - | R | 23.5 | -52.5 | -16 | 7.348 | 0 | | | | | Fusiform R, Culmen |
| - | L | -61.5 | -27.5 | 19 | 7.191 | 0 | | | | | SupraMarginal L, Post-central Gyrus |
| - | R | 38.5 | -12.5 | -1 | 7.062 | 0 | 1710 | 0 | 0 | | Insula |
| - | R | 26 | 10 | 11.5 | 7.036 | 0 | | | | | Putamen R, Extra-Nuclear |
| - | R | 31 | 2.5 | 11.5 | 6.918 | 0 | | | | | Putamen R, Claustrum |
| - | R | 61 | -22.5 | 29 | 7.024 | 0 | 657 | 0 | 0 | | SupraMarginal R, BA 40, Inferior Parietal Lobule |
| - | R | 1 | 32.5 | 14 | 6.3 | 0 | 1558 | 0 | 0 | | Cingulate Ant R |
| - | L | -6.5 | 40 | 6.5 | 6.243 | 0 | | | | | Cingulate Ant L, Anterior Cingulate |
| - | L | -9 | 50 | 19 | 5.308 | 0 | | | | | Frontal Sup Medial L, Medial Frontal Gyrus |

*Continued on next page*

Table S13 – *Continued from previous page*

| Direction | Hemisphere | MNI Coordinates | | | Peak | | Cluster | | | Peak (SVC) | Region |
|---|---|---|---|---|---|---|---|---|---|---|---|
| | | x | y | z | Z | p(Z) *unc.* | k | p(k) *unc.* | p(k) *FWE* | p(Z) *FWE* | |
| - | R | 48.5 | 35 | 1.5 | 5.121 | 0 | 74 | 0.015 | 0.105 | | Frontal Inf Tri R, Inferior Frontal Gyrus |
| - | R | 18.5 | -52.5 | 66.5 | 4.604 | 0 | 87 | 0.01 | 0.067 | | Parietal Sup R, Postcentral Gyrus |
| - | R | 11 | 40 | 49 | 4.445 | 0 | 52 | 0.037 | 0.234 | | Frontal Sup Medial R, BA 8, Superior Frontal Gyrus |
| - | L | -39 | -80 | 34 | 4.299 | 0 | 62 | 0.024 | 0.162 | | Occipital Mid L, BA 19, Superior Occipital Gyrus |
| - | L | -51.5 | -65 | 31.5 | 3.584 | 0 | | | | | Angular L, BA 39, Angular Gyrus |
| - | L | -24 | 22.5 | 44 | 3.792 | 0 | 17 | 0.209 | 0.78 | | Frontal Mid 2 L, BA 8, Middle Frontal Gyrus |
| - | L | -6.5 | 40 | 49 | 3.569 | 0 | 32 | 0.092 | 0.486 | | Frontal Sup Medial L, BA 8, Superior Frontal Gyrus |

*Continued on next page*

Table S13 – *Continued from previous page*

| Direction | Hemisphere | MNI Coordinates | | | Peak | | Cluster | | | Peak (SVC) | Region |
|---|---|---|---|---|---|---|---|---|---|---|---|
| | | x | y | z | Z | p(Z) *unc.* | k | p(k) *unc.* | p(k) *FWE* | p(Z) *FWE* | |
| - | L | -11.5 | 32.5 | 54 | 3.544 | 0 | | | | | Frontal Sup 2 L, BA 6, Superior Frontal Gyrus |
| + | L | -24 | -12.5 | 6.5 | 4.959 | 0 | 3 | 0.611 | 0.158 | 0 | Putamen, Lentiform Nucleus |
| + | L | -19 | 10 | -3.5 | 4.717 | 0 | 6 | 0.457 | 0.121 | 0.001 | Putamen L, Putamen, Lentiform Nucleus |
| + | R | 31 | -2.5 | -1 | 4.507 | 0 | 12 | 0.289 | 0.078 | 0.002 | Putamen R, Extra-Nuclear |
| + | L | -26.5 | -15 | 9 | 3.904 | 0 | 1 | 0.789 | 0.2 | 0.023 | Putamen L, Putamen, Lentiform Nucleus |
| - | R | 26 | 12.5 | 6.5 | 7.015 | 0 | 316 | 0 | 0 | 0 | Putamen R, Extra-Nuclear |
| - | R | 28.5 | 2.5 | 11.5 | 6.894 | 0 | | | | 0 | Putamen R, Extra-Nuclear |
| - | R | 26 | 0 | 1.5 | 6.31 | 0 | | | | 0 | Putamen R, Putamen, Lentiform Nucleus |

*Continued on next page*

Table S13 – *Continued from previous page*

| Direction | Hemisphere | MNI Coordinates | | | Peak | | Cluster | | | Peak (SVC) | Region |
|---|---|---|---|---|---|---|---|---|---|---|---|
| | | x | y | z | Z | p(Z) *unc.* | k | p(k) *unc.* | p(k) *FWE* | p(Z) *FWE* | |
| - | R | 26 | 7.5 | -6 | 6.289 | 0 | | | | 0 | Putamen R, Putamen, Lentiform Nucleus |
| - | L | -26.5 | 2.5 | 11.5 | 6.771 | 0 | 287 | 0 | 0 | 0 | Putamen L, Putamen, Lentiform Nucleus |
| - | L | -26.5 | 5 | -8.5 | 6.182 | 0 | | | | 0 | Putamen L, Extra-Nuclear |
| - | L | -29 | -20 | 6.5 | 5.351 | 0 | | | | 0 | Putamen L, Putamen, Lentiform Nucleus |
| - | R | 18.5 | 12.5 | 11.5 | 5.31 | 0 | 53 | 0.035 | 0.01 | 0 | Caudate R, Extra-Nuclear |
| - | L | -14 | -12.5 | 19 | 4.144 | 0 | 20 | 0.175 | 0.048 | 0.01 | Caudate L, Caudate Body, Caudate |
| - | L | -16.5 | 5 | 19 | 3.253 | 0.001 | 4 | 0.551 | 0.144 | 0.165 | Caudate L, Extra-Nuclear |

**Table S14** | Probe > baseline (Response).

| Direction | Hemisphere | MNI Coordinates | | | Peak | | Cluster | | | Peak (SVC) | Region |
|---|---|---|---|---|---|---|---|---|---|---|---|
| | | x | y | z | Z | p(Z) *unc.* | k | p(k) *unc.* | p(k) *FWE* | p(Z) *FWE* | |
| + | R | 13.5 | -40 | -13.5 | Inf | 0 | 9994 | 0 | 0 | | Cerebelum 4 5 R, Culmen |
| + | R | 3.5 | -45 | -16 | Inf | 0 | | | | | Vermis 3, Cerebellar Lingual |
| + | L | -6.5 | -10 | 54 | Inf | 0 | | | | | Supp Motor Area L, Medial Frontal Gyrus |
| + | R | 31 | -55 | -23.5 | Inf | 0 | 125 | 0.002 | 0.015 | | Cerebelum 6 R, Culmen |
| + | R | 43.5 | -40 | -8.5 | 5.605 | 0 | | | | | Sub-Gyral |
| + | R | 51 | -67.5 | -6 | 3.943 | 0 | 3 | 0.595 | 0.99 | | Temporal Inf R, Middle Occipital Gyrus |
| + | R | 41 | 37.5 | 6.5 | 3.537 | 0 | 5 | 0.482 | 0.977 | | Frontal Inf Tri R, Inferior Frontal Gyrus |
| - | L | -4 | -62.5 | 21.5 | Inf | 0 | 26749 | 0 | 0 | | Precuneus L, BA 31, Precuneus |
| - | L | -11.5 | 47.5 | 1.5 | Inf | 0 | | | | | Cingulate Ant L, Medial Frontal Gyrus |
| - | R | 41 | -12.5 | -1 | Inf | 0 | | | | | Insula R, Insula |
| - | R | 16 | -35 | -28.5 | 5.672 | 0 | 60 | 0.022 | 0.156 | | Cerebelum 3 R |

*Continued on next page*

Table S14 – *Continued from previous page*

| Direction | Hemisphere | MNI Coordinates | | | Peak | | Cluster | | | Peak (SVC) | Region |
|---|---|---|---|---|---|---|---|---|---|---|---|
| | | x | y | z | Z | p(Z) *unc.* | k | p(k) *unc.* | p(k) *FWE* | p(Z) *FWE* | |
| - | R | 11 | -47.5 | -41 | 4.904 | 0 | 16 | 0.205 | 0.797 | | Cerebelum 9 R, Cerebellar Tonsil |
| - | R | 18.5 | -67.5 | -28.5 | 4.62 | 0 | 11 | 0.291 | 0.896 | | Cerebelum 6 R, Pyramis |
| - | L | -1.5 | -2.5 | -3.5 | 3.33 | 0 | 1 | 0.779 | 0.998 | | Extra-Nuclear |
| - | L | -11.5 | -60 | -41 | 3.117 | 0.001 | 1 | 0.779 | 0.998 | | Cerebelum 8 L, Inferior Semi-Lunar Lobule |
| + | R | 31 | -5 | -6 | 7.714 | 0 | 241 | 0 | 0 | 0 | Putamen R, Extra-Nuclear |
| + | R | 23.5 | -2.5 | 6.5 | 7.64 | 0 | | | | 0 | Pallidum R, Putamen, Lentiform Nucleus |
| + | R | 28.5 | 2.5 | -1 | 7.247 | 0 | | | | 0 | Putamen R, Putamen, Lentiform Nucleus |
| + | L | -14 | 0 | 19 | 5.247 | 0 | 34 | 0.073 | 0.021 | 0 | Caudate L, Caudate Body, Caudate |
| + | L | -9 | 15 | 11.5 | 4.939 | 0 | 8 | 0.368 | 0.104 | 0 | Caudate L, Caudate Body, Caudate |

*Continued on next page*

Table S14 – *Continued from previous page*

| Direction | Hemisphere | MNI Coordinates | | | Peak | | Cluster | | | Peak (SVC) | Region |
|---|---|---|---|---|---|---|---|---|---|---|---|
| | | x | y | z | Z | p(Z) *unc.* | k | p(k) *unc.* | p(k) *FWE* | p(Z) *FWE* | |
| + | L | -9 | 17.5 | 6.5 | 4.837 | 0 | | | | 0.001 | Caudate L, Caudate Body, Caudate |
| - | R | 31 | -12.5 | 4 | Inf | 0 | 273 | 0 | 0 | 0 | Putamen R, Putamen, Lentiform Nucleus |
| - | R | 26 | 5 | -11 | 6.252 | 0 | | | | 0 | Subcallosal Gyrus |
| - | R | 26 | 0 | -8.5 | 6.072 | 0 | | | | 0 | Extra-Nuclear |
| - | R | 28.5 | 5 | 9 | 5.237 | 0 | | | | 0 | Putamen R, Putamen, Lentiform Nucleus |
| - | L | -29 | -22.5 | 4 | 6.035 | 0 | 51 | 0.032 | 0.01 | 0 | Putamen, Lentiform Nucleus |
| - | L | -26.5 | -10 | 14 | 5.405 | 0 | | | | 0 | Putamen L, Extra-Nuclear |
| - | L | -26.5 | -15 | 11.5 | 5.364 | 0 | | | | 0 | Extra-Nuclear |
| - | R | 18.5 | 15 | 16.5 | 5.721 | 0 | 48 | 0.037 | 0.011 | 0 | Caudate R, Extra-Nuclear |
| - | R | 18.5 | 5 | 21.5 | 5.003 | 0 | | | | 0 | Caudate R, Extra-Nuclear |

*Continued on next page*

Table S14 – *Continued from previous page*

| Direction | Hemisphere | MNI Coordinates | | | Peak | | Cluster | | | Peak (SVC) | Region |
|---|---|---|---|---|---|---|---|---|---|---|---|
| | | x | y | z | Z | p(Z) *unc.* | k | p(k) *unc.* | p(k) *FWE* | p(Z) *FWE* | |
| - | L | -29 | -15 | -11 | 5.028 | 0 | 3 | 0.595 | 0.162 | 0 | Hippocampus L, Hippocampus, Parahippocampal Gyrus |
| - | L | -26.5 | 10 | -8.5 | 4.82 | 0 | 22 | 0.141 | 0.041 | 0.001 | Putamen L, Extra-Nuclear |
| - | L | -16.5 | 10 | 19 | 3.957 | 0 | 9 | 0.339 | 0.096 | 0.02 | Caudate L, Extra-Nuclear |
| - | L | -16.5 | -15 | 24 | 3.555 | 0 | 6 | 0.438 | 0.122 | 0.075 | Caudate L, Caudate Body, Caudate |
| - | R | 18.5 | -22.5 | 21.5 | 3.326 | 0 | 1 | 0.779 | 0.207 | 0.143 | Lateral Ventricle |



**Table S15** | Selective input-gating (selective > global 1 at sample).

| Direction | Hemisphere | MNI Coordinates | | | Peak | | Cluster | | | Peak (SVC) | Region |
|---|---|---|---|---|---|---|---|---|---|---|---|
| | | x | y | z | Z | p(Z) *unc.* | k | p(k) *unc.* | p(k) *FWE* | p(Z) *FWE* | |
| + | L | -31.5 | -60 | -13.5 | Inf | 0 | 10741 | 0 | 0 | | Fusiform L, Declive |
| + | L | -39 | -70 | -13.5 | Inf | 0 | | | | | Fusiform L, Fusiform Gyrus |

*Continued on next page*

| Direction | Hemisphere | MNI Coordinates | | | Peak | | Cluster | | | Peak (SVC) | Region |
|---|---|---|---|---|---|---|---|---|---|---|---|
| | | x | y | z | Z | p(Z) *unc.* | k | p(k) *unc.* | p(k) *FWE* | p(Z) *FWE* | |
| + | R | 38.5 | -60 | -16 | Inf | 0 | | | | | Fusiform R, Declive |
| + | L | -1.5 | 12.5 | 49 | Inf | 0 | 6957 | 0 | 0 | | Supp Motor Area L, BA 6, Superior Frontal Gyrus |
| + | L | -31.5 | -5 | 54 | Inf | 0 | | | | | Precentral L, BA 6, Middle Frontal Gyrus |
| + | L | -39 | 15 | 6.5 | Inf | 0 | | | | | Insula L, Insula |
| + | L | -16.5 | 12.5 | 6.5 | 6.575 | 0 | 124 | 0.002 | 0.018 | | Extra-Nuclear |
| + | R | 1 | -52.5 | -33.5 | 5.941 | 0 | 63 | 0.022 | 0.15 | | Vermis 9, Fourth Ventricle |
| + | R | 16 | 10 | 9 | 4.581 | 0 | 42 | 0.054 | 0.33 | | Caudate R, Extra-Nuclear |
| + | L | -6.5 | -97.5 | 11.5 | 4.353 | 0 | 1 | 0.785 | 0.997 | | Cuneus L, BA 18, Cuneus |
| + | L | -6.5 | -30 | 26.5 | 4.11 | 0 | 24 | 0.134 | 0.631 | | Cingulate Gyrus |
| + | R | 28.5 | 0 | -21 | 4.064 | 0 | 4 | 0.544 | 0.983 | | Amygdala R, Uncus |
| + | R | 6 | -32.5 | 26.5 | 3.564 | 0 | 10 | 0.326 | 0.911 | | Cingulate Gyrus |
| + | R | 21 | -5 | -16 | 3.447 | 0 | 6 | 0.45 | 0.965 | | Amygdala R, Amygdala, Parahippocampal Gyrus |

*Continued on next page*

Table S15 – *Continued from previous page*

| Direction | Hemisphere | MNI Coordinates | | | Peak | | Cluster | | | Peak (SVC) | Region |
|---|---|---|---|---|---|---|---|---|---|---|---|
| | | x | y | z | Z | p(Z) *unc.* | k | p(k) *unc.* | p(k) *FWE* | p(Z) *FWE* | |
| + | R | 8.5 | -52.5 | 51.5 | 3.409 | 0 | 4 | 0.544 | 0.983 | | Precuneus R, Precuneus |
| + | L | -9 | -35 | -26 | 3.249 | 0.001 | 4 | 0.544 | 0.983 | | Cerebelum 3 L |
| + | L | -11.5 | -25 | 14 | 3.152 | 0.001 | 1 | 0.785 | 0.997 | | Thalamus L, Pulvinar, Thalamus |
| + | R | 58.5 | -42.5 | 26.5 | 3.133 | 0.001 | 1 | 0.785 | 0.997 | | SupraMarginal R, BA 40, Inferior Parietal Lobule |
| + | R | 8.5 | -22.5 | 26.5 | 3.115 | 0.001 | 1 | 0.785 | 0.997 | | Cingulate Gyrus |
| - | L | -61.5 | -30 | 14 | Inf | 0 | 13256 | 0 | 0 | | Temporal Sup L, Superior Temporal Gyrus |
| - | R | 63.5 | -25 | 1.5 | Inf | 0 | | | | | Temporal Sup R, BA 22, Superior Temporal Gyrus |
| - | L | -61.5 | -22.5 | 1.5 | Inf | 0 | | | | | Temporal Sup L, Superior Temporal Gyrus |
| - | R | 13.5 | 42.5 | 49 | Inf | 0 | 4619 | 0 | 0 | | Frontal Sup 2 R |
| - | R | 26 | 30 | 49 | Inf | 0 | | | | | Frontal Sup 2 R, Superior Frontal Gyrus |

*Continued on next page*

Table S15 – *Continued from previous page*

| Direction | Hemisphere | MNI Coordinates | | | Peak | | Cluster | | | Peak (SVC) | Region |
|---|---|---|---|---|---|---|---|---|---|---|---|
| | | x | y | z | Z | p(Z) *unc.* | k | p(k) *unc.* | p(k) *FWE* | p(Z) *FWE* | |
| - | R | 21 | 27.5 | 56.5 | Inf | 0 | | | | | Frontal Sup 2 R, BA 6, Superior Frontal Gyrus |
| - | R | 56 | 25 | 14 | 6.464 | 0 | 176 | 0.001 | 0.004 | | Frontal Inf Tri R, Inferior Frontal Gyrus |
| - | R | 43.5 | 50 | -1 | 6.377 | 0 | | | | | Frontal Mid 2 R, Middle Frontal Gyrus |
| - | R | 46 | 40 | -3.5 | 5.446 | 0 | | | | | Frontal Inf Orb 2 R, Middle Frontal Gyrus |
| - | L | -29 | -22.5 | -13.5 | 4.562 | 0 | 42 | 0.054 | 0.33 | | Hippocampus L, Hippocampus, Parahippocampal Gyrus |
| - | L | -11.5 | -30 | 4 | 4.363 | 0 | 13 | 0.263 | 0.859 | | Thalamus L, Pulvinar, Thalamus |
| - | L | -6.5 | 17.5 | 1.5 | 4.156 | 0 | 5 | 0.493 | 0.975 | | Caudate L, Caudate Head, Caudate |
| - | L | -9 | -52.5 | -41 | 4.017 | 0 | 8 | 0.38 | 0.941 | | Cerebelum 9 L, Cerebellar Tonsil |

*Continued on next page*

Table S15 – *Continued from previous page*

| **Direction** | **Hemisphere** | **MNI Coordinates** | | | **Peak** | | **Cluster** | | | **Peak (SVC)** | **Region** |
|---|---|---|---|---|---|---|---|---|---|---|---|
| | | x | y | z | Z | p(Z) *unc.* | k | p(k) *unc.* | p(k) *FWE* | p(Z) *FWE* | |
| - | R | 16 | -40 | -28.5 | 3.947 | 0 | 10 | 0.326 | 0.911 | | |
| - | R | 16 | -20 | -21 | 3.719 | 0 | 7 | 0.413 | 0.954 | | |
| - | R | 11 | 22.5 | -1 | 3.225 | 0.001 | 2 | 0.681 | 0.994 | | Caudate R, Lateral Ventricle |
| - | R | 46 | 42.5 | 19 | 3.213 | 0.001 | 1 | 0.785 | 0.997 | | Frontal Mid 2 R, BA 46, Middle Frontal Gyrus |
| - | R | 16 | -45 | 14 | 3.194 | 0.001 | 1 | 0.785 | 0.997 | | Corpus Callosum, Extra-Nuclear |
| - | R | 6 | 12.5 | 1.5 | 3.103 | 0.001 | 1 | 0.785 | 0.997 | | Caudate R, Caudate Head, Caudate |
| + | L | -16.5 | 12.5 | 9 | 6.313 | 0 | 70 | 0.016 | 0.005 | 0 | Caudate L, Extra-Nuclear |
| + | R | 16 | 12.5 | 9 | 4.43 | 0 | 14 | 0.246 | 0.068 | 0.003 | Caudate R, Extra-Nuclear |
| - | R | 31 | -17.5 | 4 | 6.767 | 0 | 148 | 0.001 | 0 | 0 | Putamen R, Putamen, Lentiform Nucleus |

*Continued on next page*

Table S15 – *Continued from previous page*

| **Direction** | **Hemisphere** | **MNI Coordinates** | | | **Peak** | | **Cluster** | | | **Peak (SVC)** | **Region** |
|---|---|---|---|---|---|---|---|---|---|---|---|
| | | x | y | z | Z | p(Z) *unc.* | k | p(k) *unc.* | p(k) *FWE* | p(Z) *FWE* | |
| - | R | 31 | 2.5 | -6 | 4.944 | 0 | | | | 0 | Putamen R, Extra-Nuclear |
| - | L | -31.5 | -15 | 4 | 6.182 | 0 | 105 | 0.005 | 0.001 | 0 | Extra-Nuclear |
| - | L | -26.5 | -15 | 11.5 | 4.038 | 0 | | | | 0.013 | Extra-Nuclear |
| - | L | -26.5 | 0 | -8.5 | 3.777 | 0 | | | | 0.033 | Putamen L, Extra-Nuclear |
| - | L | -29 | -2.5 | -6 | 3.666 | 0 | | | | 0.048 | Putamen L, Extra-Nuclear |
| - | L | -6.5 | 15 | 1.5 | 3.7 | 0 | 2 | 0.681 | 0.178 | 0.043 | Caudate L, Caudate Head, Caudate |
| - | R | 11 | 22.5 | -1 | 3.225 | 0.001 | 2 | 0.681 | 0.178 | 0.179 | Caudate R, Lateral Ventricle |
| - | R | 6 | 12.5 | 1.5 | 3.103 | 0.001 | 1 | 0.785 | 0.202 | 0.243 | Caudate R, Caudate Head, Caudate |

**Table S16** | Selective output-gating (selective > global 1 at retro-cue).

| Direction | Hemisphere | MNI Coordinates | | | Peak | | Cluster | | | Peak (SVC) | Region |
|---|---|---|---|---|---|---|---|---|---|---|---|
| | | x | y | z | Z | p(Z) *unc.* | k | p(k) *unc.* | p(k) *FWE* | p(Z) *FWE* | |
| + | L | -4 | 12.5 | 51.5 | Inf | 0 | 8949 | 0 | 0 | | Supp Motor Area L, Superior Frontal Gyrus |
| + | L | -31.5 | 25 | 1.5 | Inf | 0 | | | | | Insula L, BA 47, Inferior Frontal Gyrus |
| + | L | -36.5 | -2.5 | 59 | Inf | 0 | | | | | Precentral L, Middle Frontal Gyrus |
| + | L | -29 | -60 | 49 | Inf | 0 | 4193 | 0 | 0 | | Parietal Inf L, BA 7, Superior Parietal Lobule |
| + | L | -44 | -65 | -13.5 | Inf | 0 | | | | | Fusiform L, Fusiform Gyrus |
| + | L | -29 | -70 | 36.5 | Inf | 0 | | | | | Occipital Mid L, Precuneus |
| + | R | 8.5 | -75 | -21 | Inf | 0 | 155 | 0.001 | 0.006 | | Cerebelum 6 R, Declive |
| + | L | -9 | -72.5 | -21 | 6.66 | 0 | | | | | Cerebelum 6 L, Declive |
| + | R | 6 | -72.5 | -33.5 | 3.478 | 0 | | | | | Cerebelum Crus2 R, Uvula |
| + | R | 28.5 | -60 | -28.5 | Inf | 0 | 32 | 0.083 | 0.472 | | Cerebelum 6 R, Pyramis |

*Continued on next page*

Table S16 – *Continued from previous page*

| Direction | Hemisphere | MNI Coordinates | | | Peak | | Cluster | | | Peak (SVC) | Region |
|---|---|---|---|---|---|---|---|---|---|---|---|
| | | x | y | z | Z | p(Z) *unc.* | k | p(k) *unc.* | p(k) *FWE* | p(Z) *FWE* | |
| + | R | 61 | -45 | 24 | 6.894 | 0 | 638 | 0 | 0 | | SupraMarginal R, BA 40, Supramarginal Gyrus |
| + | R | 48.5 | -27.5 | -6 | 6.074 | 0 | | | | | Middle Temporal Gyrus |
| + | R | 51 | -37.5 | 6.5 | 5.679 | 0 | | | | | Temporal Mid R, Superior Temporal Gyrus |
| + | R | 38.5 | -45 | -18.5 | 5.967 | 0 | 69 | 0.016 | 0.114 | | Fusiform R, Fusiform Gyrus |
| + | R | 33.5 | -57.5 | 9 | 5.376 | 0 | 515 | 0 | 0 | | Sub-Gyral |
| + | R | 3.5 | -77.5 | 14 | 4.537 | 0 | | | | | Calcarine L, BA 18, Cuneus |
| + | L | -14 | -80 | 14 | 4.346 | 0 | | | | | Cuneus L, Cuneus |
| + | L | -4 | -52.5 | -11 | 4.833 | 0 | 81 | 0.01 | 0.073 | | Cerebelum 4 5 L, Culmen |
| + | L | -9 | -30 | 61.5 | 4.674 | 0 | 349 | 0 | 0 | | Paracentral Lobule L, Sub-Gyral |
| + | R | 8.5 | -30 | 61.5 | 4.606 | 0 | | | | | Paracentral Lobule R, Paracentral Lobule |
| + | R | 8.5 | -35 | 69 | 4.587 | 0 | | | | | Paracentral Lobule R |

*Continued on next page*

Table S16 – *Continued from previous page*

| Direction | Hemisphere | MNI Coordinates | | | Peak | | Cluster | | | Peak (SVC) | Region |
|---|---|---|---|---|---|---|---|---|---|---|---|
| | | x | y | z | Z | p(Z) *unc.* | k | p(k) *unc.* | p(k) *FWE* | p(Z) *FWE* | |
| + | R | 33.5 | -70 | 31.5 | 4.446 | 0 | 37 | 0.065 | 0.391 | | Occipital Mid R, Sub-Gyral |
| + | R | 43.5 | -20 | 49 | 4.116 | 0 | 57 | 0.026 | 0.18 | | Postcentral R, Postcentral Gyrus |
| + | R | 8.5 | -70 | -38.5 | 3.905 | 0 | 2 | 0.676 | 0.994 | | Cerebelum 8 R, Inferior Semi-Lunar Lobule |
| + | R | 28.5 | -25 | 69 | 3.869 | 0 | 41 | 0.053 | 0.335 | | Precentral R |
| + | R | 36 | -27.5 | 64 | 3.613 | 0 | | | | | Precentral R, Postcentral Gyrus |
| + | L | -9 | -70 | -38.5 | 3.751 | 0 | 5 | 0.487 | 0.976 | | Cerebelum 8 L, Inferior Semi-Lunar Lobule |
| + | L | -11.5 | -12.5 | 1.5 | 3.657 | 0 | 18 | 0.184 | 0.755 | | Thalamus L, Thalamus |
| + | L | -19 | -47.5 | 19 | 3.603 | 0 | 13 | 0.256 | 0.859 | | Extra-Nuclear |
| + | L | -4 | -32.5 | 24 | 3.576 | 0 | 16 | 0.209 | 0.798 | | BA 23, Posterior Cingulate |
| + | L | -9 | -20 | -16 | 3.26 | 0.001 | 5 | 0.487 | 0.976 | | |
| + | R | 26 | -27.5 | 51.5 | 3.21 | 0.001 | 2 | 0.676 | 0.994 | | BA 4, Precentral Gyrus |

*Continued on next page*

Table S16 – *Continued from previous page*

| Direction | Hemisphere | MNI Coordinates | | | Peak | | Cluster | | | Peak (SVC) | Region |
|---|---|---|---|---|---|---|---|---|---|---|---|
| | | x | y | z | Z | p(Z) *unc.* | k | p(k) *unc.* | p(k) *FWE* | p(Z) *FWE* | |
| + | L | -31.5 | -27.5 | 54 | 3.204 | 0.001 | 3 | 0.599 | 0.99 | | Precentral L, BA 4, Precentral Gyrus |
| + | L | -26.5 | -60 | 6.5 | 3.132 | 0.001 | 2 | 0.676 | 0.994 | | Calcarine L, Extra-Nuclear |
| - | R | 21 | 10 | 6.5 | Inf | 0 | 2449 | 0 | 0 | | Putamen R, Putamen, Lentiform Nucleus |
| - | R | 26 | 5 | -3.5 | Inf | 0 | | | | | Putamen R, Putamen, Lentiform Nucleus |
| - | R | 61 | -25 | 26.5 | Inf | 0 | | | | | SupraMarginal R, Inferior Parietal Lobule |
| - | L | -6.5 | -65 | 21.5 | Inf | 0 | 5878 | 0 | 0 | | Cuneus L, Precuneus |
| - | R | 8.5 | -47.5 | 34 | Inf | 0 | | | | | Cingulate Mid R, BA 31, Precuneus |
| - | L | -11.5 | -52.5 | 29 | Inf | 0 | | | | | Precuneus L, Cingulate Gyrus |
| - | R | 28.5 | -72.5 | -13.5 | Inf | 0 | 3108 | 0 | 0 | | Fusiform R, Declive |

*Continued on next page*

Table S16 – *Continued from previous page*

| Direction | Hemisphere | MNI Coordinates | | | Peak | | Cluster | | | Peak (SVC) | Region |
|---|---|---|---|---|---|---|---|---|---|---|---|
| | | x | y | z | Z | p(Z) *unc.* | k | p(k) *unc.* | p(k) *FWE* | p(Z) *FWE* | |
| - | L | -24 | -72.5 | -8.5 | Inf | 0 | | | | | Fusiform L, Lingual Gyrus |
| - | R | 8.5 | -82.5 | -8.5 | Inf | 0 | | | | | Lingual R, BA 18, Lingual Gyrus |
| - | L | -1.5 | 32.5 | -1 | Inf | 0 | 2501 | 0 | 0 | | Cingulate Ant L, BA 24, Anterior Cingulate |
| - | R | 26 | 27.5 | 51.5 | Inf | 0 | | | | | Frontal Sup 2 R, BA 8, Superior Frontal Gyrus |
| - | L | -34 | 20 | 49 | 7.179 | 0 | | | | | Frontal Mid 2 L, Superior Frontal Gyrus |
| - | L | -44 | -75 | 39 | Inf | 0 | 293 | 0 | 0 | | |
| - | R | 43.5 | -72.5 | 36.5 | 7.723 | 0 | 397 | 0 | 0 | | Angular R, BA 39, Precuneus |
| - | R | 46 | -67.5 | 44 | 7.661 | 0 | | | | | Angular R |
| - | L | -24 | -50 | 69 | 7.495 | 0 | 307 | 0 | 0 | | Parietal Sup L |
| - | L | -29 | -40 | 61.5 | 5.495 | 0 | | | | | Postcentral L, BA 5, Postcentral Gyrus |

*Continued on next page*

Table S16 – *Continued from previous page*

| Direction | Hemisphere | MNI Coordinates | | | Peak | | Cluster | | | Peak (SVC) | Region |
|---|---|---|---|---|---|---|---|---|---|---|---|
| | | x | y | z | Z | p(Z) *unc.* | k | p(k) *unc.* | p(k) *FWE* | p(Z) *FWE* | |
| - | R | 56 | -57.5 | -3.5 | 5.749 | 0 | 40 | 0.056 | 0.348 | | Temporal Inf R, BA 37, Inferior Temporal Gyrus |
| - | R | 21 | -50 | 69 | 5.749 | 0 | 144 | 0.001 | 0.009 | | Parietal Sup R |
| - | L | -24 | -25 | -16 | 4.916 | 0 | 65 | 0.019 | 0.133 | | BA 35, Parahippocampal Gyrus |
| - | R | 48.5 | 45 | -1 | 4.801 | 0 | 35 | 0.072 | 0.422 | | Frontal Inf Tri R, Inferior Frontal Gyrus |
| - | L | -16.5 | -12.5 | 64 | 4.222 | 0 | 34 | 0.075 | 0.438 | | Supp Motor Area L, Superior Frontal Gyrus |
| - | R | 11 | -12.5 | 24 | 3.546 | 0 | 5 | 0.487 | 0.976 | | Lateral Ventricle |
| - | R | 8.5 | -10 | 64 | 3.361 | 0 | 3 | 0.599 | 0.99 | | Supp Motor Area R, BA 6, Medial Frontal Gyrus |
| - | R | 56 | 27.5 | 14 | 3.319 | 0 | 1 | 0.782 | 0.997 | | Frontal Inf Tri R, BA 45, Inferior Frontal Gyrus |
| - | R | 28.5 | -20 | -16 | 3.302 | 0 | 6 | 0.443 | 0.966 | | Hippocampus R, Parahippocampal Gyrus |

*Continued on next page*

Table S16 – *Continued from previous page*

| Direction | Hemisphere | MNI Coordinates | | | Peak | | Cluster | | | Peak (SVC) | Region |
|---|---|---|---|---|---|---|---|---|---|---|---|
| | | x | y | z | Z | p(Z) *unc.* | k | p(k) *unc.* | p(k) *FWE* | p(Z) *FWE* | |
| - | R | 61 | 10 | 24 | 3.228 | 0.001 | 1 | 0.782 | 0.997 | | Precentral R, BA 9, Inferior Frontal Gyrus |
| - | R | 21 | 7.5 | 6.5 | Inf | 0 | 320 | 0 | 0 | 0 | Putamen R, Putamen, Lentiform Nucleus |
| - | R | 26 | 5 | -3.5 | Inf | 0 | | | | 0 | Putamen R, Putamen, Lentiform Nucleus |
| - | R | 31 | -17.5 | 6.5 | 6.534 | 0 | | | | 0 | Putamen R, Putamen, Lentiform Nucleus |
| - | L | -26.5 | 2.5 | -8.5 | Inf | 0 | 301 | 0 | 0 | 0 | Putamen L, Extra-Nuclear |
| - | L | -24 | 7.5 | 4 | Inf | 0 | | | | 0 | Putamen L, Putamen, Lentiform Nucleus |
| - | L | -29 | -5 | 1.5 | 7.709 | 0 | | | | 0 | Putamen L, Putamen, Lentiform Nucleus |
| - | L | -26.5 | -10 | 14 | 7.453 | 0 | | | | 0 | Putamen L, Extra-Nuclear |
| - | L | -31.5 | -15 | 4 | 6.883 | 0 | | | | 0 | Extra-Nuclear |

*Continued on next page*

Table S16 – *Continued from previous page*

| Direction | Hemisphere | MNI Coordinates | | | Peak | | Cluster | | | Peak (SVC) | Region |
|---|---|---|---|---|---|---|---|---|---|---|---|
| | | x | y | z | Z | p(Z) *unc.* | k | p(k) *unc.* | p(k) *FWE* | p(Z) *FWE* | |
| - | R | 18.5 | 15 | 9 | Inf | 0 | 105 | 0.004 | 0.001 | 0 | Caudate R, Extra-Nuclear |
| - | R | 18.5 | 2.5 | 19 | 5.004 | 0 | | | | 0 | Caudate R, Extra-Nuclear |
| - | L | -16.5 | -17.5 | 21.5 | 3.966 | 0 | 4 | 0.538 | 0.146 | 0.017 | Caudate L, Caudate Body, Caudate |
| - | L | -14 | -15 | 19 | 3.125 | 0.001 | | | | 0.235 | Extra-Nuclear |

**Table S17** | Sulpiride > placebo during input-gating (sulpiride > placebo at sample).

| Direction | Hemisphere | MNI Coordinates | | | Peak | | Cluster | | | Peak (SVC) | Region |
|---|---|---|---|---|---|---|---|---|---|---|---|
| | | x | y | z | Z | p(Z) *unc.* | k | p(k) *unc.* | p(k) *FWE* | p(Z) *FWE* | |
| + | R | 11 | -2.5 | 19 | 5.257 | 0 | 193 | 0 | 0.002 | | Lateral Ventricle |
| + | R | 8.5 | 10 | 11.5 | 4.38 | 0 | | | | | Caudate Body, Caudate |
| + | R | 11 | 15 | -1 | 3.351 | 0 | | | | | Caudate R, Caudate Head, Caudate |
| + | L | -11.5 | 15 | 9 | 4.012 | 0 | 25 | 0.127 | 0.61 | | Caudate L, Caudate Body, Caudate |

*Continued on next page*

Table S17 – *Continued from previous page*

| Direction | Hemisphere | MNI Coordinates | | | Peak | | Cluster | | | Peak (SVC) | Region |
|---|---|---|---|---|---|---|---|---|---|---|---|
| | | x | y | z | Z | p(Z) *unc.* | k | p(k) *unc.* | p(k) *FWE* | p(Z) *FWE* | |
| + | R | 23.5 | 10 | -3.5 | 3.888 | 0 | 23 | 0.142 | 0.651 | | Putamen R, Putamen, Lentiform Nucleus |
| + | L | -9 | -32.5 | 36.5 | 3.822 | 0 | 79 | 0.012 | 0.083 | | Cingulate Mid L, Cingulate Gyrus |
| + | R | 8.5 | -32.5 | 36.5 | 3.601 | 0 | | | | | Cingulate Mid R, Cingulate Gyrus |
| + | R | 1 | -37.5 | 41.5 | 3.418 | 0 | | | | | Cingulate Mid R, Cingulate Gyrus |
| + | L | -39 | -5 | 26.5 | 3.812 | 0 | 12 | 0.282 | 0.877 | | Precentral Gyrus |
| + | R | 8.5 | 0 | 1.5 | 3.802 | 0 | 7 | 0.413 | 0.954 | | Extra-Nuclear |
| + | R | 3.5 | -57.5 | -18.5 | 3.775 | 0 | 14 | 0.246 | 0.839 | | Vermis 4 5, Culmen |
| + | L | -46.5 | -70 | 41.5 | 3.689 | 0 | 39 | 0.062 | 0.37 | | Angular L |
| + | L | -51.5 | -62.5 | 41.5 | 3.55 | 0 | | | | | Parietal Inf L, BA 39, Inferior Parietal Lobule |
| + | L | -36.5 | -67.5 | 34 | 3.252 | 0.001 | | | | | Occipital Mid L, Precuneus |
| + | R | 21 | -52.5 | -28.5 | 3.605 | 0 | 9 | 0.351 | 0.927 | | Cerebelum 6 R |

*Continued on next page*

Table S17 – *Continued from previous page*

| Direction | Hemisphere | MNI Coordinates | | | Peak | | Cluster | | | Peak (SVC) | Region |
|---|---|---|---|---|---|---|---|---|---|---|---|
| | | x | y | z | Z | p(Z) *unc.* | k | p(k) *unc.* | p(k) *FWE* | p(Z) *FWE* | |
| + | L | -41.5 | 45 | 14 | 3.449 | 0 | 11 | 0.302 | 0.895 | | Frontal Inf Tri L, Middle Frontal Gyrus |
| + | L | -19 | 20 | 44 | 3.355 | 0 | 3 | 0.605 | 0.989 | | Frontal Sup 2 L, BA 8, Superior Frontal Gyrus |
| + | L | -14 | -57.5 | 29 | 3.339 | 0 | 2 | 0.681 | 0.994 | | Cuneus L, Precuneus |
| + | L | -36.5 | -20 | 29 | 3.332 | 0 | 3 | 0.605 | 0.989 | | Postcentral Gyrus |
| + | L | -11.5 | 2.5 | 14 | 3.329 | 0 | 6 | 0.45 | 0.965 | | Caudate L, Caudate Body, Caudate |
| + | R | 38.5 | 52.5 | 16.5 | 3.257 | 0.001 | 3 | 0.605 | 0.989 | | Frontal Mid 2 R, Superior Frontal Gyrus |
| + | L | -34 | -7.5 | 36.5 | 3.248 | 0.001 | 4 | 0.544 | 0.983 | | Sub-Gyral |
| + | R | 53.5 | -57.5 | 44 | 3.174 | 0.001 | 6 | 0.45 | 0.965 | | Parietal Inf R |
| + | R | 48.5 | -67.5 | 41.5 | 3.115 | 0.001 | 1 | 0.785 | 0.997 | | Angular R |
| - | L | -34 | 30 | 6.5 | 4.481 | 0 | 130 | 0.002 | 0.015 | | Frontal Inf Tri L, Sub-Gyral |
| - | L | -56.5 | 12.5 | 1.5 | 3.865 | 0 | | | | | Frontal Inf Oper L |

*Continued on next page*

Table S17 – *Continued from previous page*

| Direction | Hemisphere | MNI Coordinates | | | Peak | | Cluster | | | Peak (SVC) | Region |
|---|---|---|---|---|---|---|---|---|---|---|---|
| | | x | y | z | Z | p(Z) *unc.* | k | p(k) *unc.* | p(k) *FWE* | p(Z) *FWE* | |
| - | L | -34 | 20 | -3.5 | 3.108 | 0.001 | | | | | Insula L, Inferior Frontal Gyrus |
| - | R | 8.5 | -20 | 24 | 3.959 | 0 | 30 | 0.097 | 0.514 | | Corpus Callosum, Extra-Nuclear |
| - | L | -1.5 | -45 | -1 | 3.657 | 0 | 9 | 0.351 | 0.927 | | Vermis 4 5, Culmen |
| - | R | 23.5 | -17.5 | 29 | 3.449 | 0 | 7 | 0.413 | 0.954 | | Sub-Gyral |
| - | L | -1.5 | -15 | -3.5 | 3.246 | 0.001 | 2 | 0.681 | 0.994 | | |
| - | R | 26 | -70 | 14 | 3.172 | 0.001 | 3 | 0.605 | 0.989 | | Cuneus |
| - | R | 13.5 | -30 | -1 | 3.146 | 0.001 | 2 | 0.681 | 0.994 | | Thalamus R, Thalamus |
| + | R | 11 | -2.5 | 16.5 | 5.109 | 0 | 111 | 0.004 | 0.001 | 0 | Caudate Body, Caudate |
| + | R | 8.5 | 10 | 11.5 | 4.38 | 0 | | | | 0.003 | Caudate Body, Caudate |
| + | R | 11 | 15 | -1 | 3.351 | 0 | | | | 0.127 | Caudate R, Caudate Head, Caudate |
| + | L | -11.5 | 15 | 9 | 4.012 | 0 | 24 | 0.134 | 0.038 | 0.014 | Caudate L, Caudate Body, Caudate |
| + | R | 23.5 | 10 | -3.5 | 3.888 | 0 | 23 | 0.142 | 0.04 | 0.023 | Putamen R, Putamen, Lentiform Nucleus |

*Continued on next page*

Table S17 – *Continued from previous page*

| Direction | Hemisphere | MNI Coordinates | | | Peak | | Cluster | | | Peak (SVC) | Region |
|---|---|---|---|---|---|---|---|---|---|---|---|
| | | x | y | z | Z | p(Z) *unc.* | k | p(k) *unc.* | p(k) *FWE* | p(Z) *FWE* | |
| + | L | -11.5 | 2.5 | 14 | 3.329 | 0 | 6 | 0.45 | 0.122 | 0.135 | Caudate L, Caudate Body, Caudate |
| + | R | 8.5 | 5 | 1.5 | 3.146 | 0.001 | 1 | 0.785 | 0.202 | 0.218 | Caudate Head, Caudate |

**Table S18** | Sulpiride > placebo during selective input-gating ([sulpiride > placebo] x [selective > global 1] at sample).

| Direction | Hemisphere | MNI Coordinates | | | Peak | | Cluster | | | Peak (SVC) | Region |
|---|---|---|---|---|---|---|---|---|---|---|---|
| | | x | y | z | Z | p(Z) *unc.* | k | p(k) *unc.* | p(k) *FWE* | p(Z) *FWE* | |
| + | L | -39 | -10 | 36.5 | 3.561 | 0 | 8 | 0.38 | 0.941 | | Precentral Gyrus |
| + | L | -26.5 | 5 | 16.5 | 3.41 | 0 | 4 | 0.544 | 0.983 | | Extra-Nuclear |
| + | L | -21.5 | -55 | 31.5 | 3.336 | 0 | 2 | 0.681 | 0.994 | | Precuneus |
| + | L | -14 | -5 | 31.5 | 3.257 | 0.001 | 4 | 0.544 | 0.983 | | Cingulate Gyrus |
| + | L | -19 | -42.5 | 39 | 3.204 | 0.001 | 1 | 0.785 | 0.997 | | Sub-Gyral |
| + | L | -19 | -87.5 | 1.5 | 3.202 | 0.001 | 2 | 0.681 | 0.994 | | Lingual Gyrus |
| + | L | -16.5 | -47.5 | 41.5 | 3.174 | 0.001 | 1 | 0.785 | 0.997 | | Precuneus |
| + | R | 21 | -87.5 | 6.5 | 3.14 | 0.001 | 1 | 0.785 | 0.997 | | Occipital Sup R, Cuneus |
| - | L | -14 | -55 | 4 | 4.079 | 0 | 82 | 0.01 | 0.075 | | Calcarine L, Lingual Gyrus |

*Continued on next page*

Table S18 – *Continued from previous page*

| **Direction** | **Hemisphere** | **MNI Coordinates** | | | **Peak** | | **Cluster** | | | **Peak (SVC)** | **Region** |
|---|---|---|---|---|---|---|---|---|---|---|---|
| | | x | y | z | Z | p(Z) *unc.* | k | p(k) *unc.* | p(k) *FWE* | p(Z) *FWE* | |
| - | L | -21.5 | -47.5 | 1.5 | 4.015 | 0 | | | | | Precuneus L, Sub-Gyral |
| - | L | -6.5 | -45 | 1.5 | 3.296 | 0 | | | | | Culmen |
| - | R | 21 | -15 | -11 | 4.019 | 0 | 33 | 0.083 | 0.462 | | Parahippocampal Gyrus |
| - | R | 26 | -25 | -8.5 | 3.27 | 0.001 | | | | | Hippocampus R, Parahippocampal Gyrus |
| - | L | -26.5 | -15 | -13.5 | 3.779 | 0 | 17 | 0.202 | 0.778 | | Hippocampus L, Parahippocampal Gyrus |
| - | R | 46 | -50 | 21.5 | 3.674 | 0 | 16 | 0.215 | 0.799 | | Angular R, Supra-marginal Gyrus |
| - | R | 11 | -35 | -3.5 | 3.606 | 0 | 14 | 0.246 | 0.839 | | Lingual R |
| - | R | 8.5 | -30 | -21 | 3.38 | 0 | 8 | 0.38 | 0.941 | | |
| - | R | 8.5 | -52.5 | -8.5 | 3.311 | 0 | 3 | 0.605 | 0.989 | | Cerebelum 4 5 R, Culmen |

**Table S19** | Sulpiride > placebo during selective output-gating of scenes ([sulpiride > placebo] x [selective > global 1] x [faces > scenes] at retro-cue).

| **Direction** | **Hemisphere** | **MNI Coordinates** | | | **Peak** | | **Cluster** | | | **Peak (SVC)** | **Region** |
|---|---|---|---|---|---|---|---|---|---|---|---|
| | | x | y | z | Z | p(Z) *unc.* | k | p(k) *unc.* | p(k) *FWE* | p(Z) *FWE* | |
| + | R | 6 | -45 | -8.5 | 3.785 | 0 | 20 | 0.163 | 0.712 | | Vermis 3, Cerebellar Lingual |
| + | L | -1.5 | -70 | -26 | 3.253 | 0.001 | 3 | 0.599 | 0.99 | | Vermis 7, Tuber of Vermis |
| - | R | 13.5 | 12.5 | 1.5 | 3.697 | 0 | 21 | 0.153 | 0.69 | | Caudate R, Extra-Nuclear |
| - | L | -31.5 | -42.5 | 9 | 3.336 | 0 | 5 | 0.487 | 0.976 | | Sub-Gyral |
| - | R | 8.5 | 17.5 | 16.5 | 3.265 | 0.001 | 1 | 0.782 | 0.997 | | Corpus Callosum, Extra-Nuclear |
| - | R | 16 | -52.5 | 56.5 | 3.179 | 0.001 | 2 | 0.676 | 0.994 | | Parietal Sup R, Precuneus |
| - | R | 26 | -72.5 | 16.5 | 3.133 | 0.001 | 2 | 0.676 | 0.994 | | Cuneus |
| - | R | 11 | 12.5 | 1.5 | 3.686 | 0 | 13 | 0.256 | 0.072 | 0.046 | Caudate R, Caudate Head, Caudate |
| - | R | 13.5 | 15 | 4 | 3.409 | 0 | | | | 0.109 | Caudate R, Caudate Head, Caudate |

**Table S20** | Input-gating load (global 2 > global 1 at sample).

| Direction | Hemisphere | MNI Coordinates | | | Peak | | Cluster | | | Peak (SVC) | Region |
|---|---|---|---|---|---|---|---|---|---|---|---|
| | | x | y | z | Z | p(Z) *unc.* | k | p(k) *unc.* | p(k) *FWE* | p(Z) *FWE* | |
| + | R | 1 | 12.5 | 49 | 7.174 | 0 | 830 | 0 | 0 | | Supp Motor Area L, Superior Frontal Gyrus |
| + | R | 36 | 17.5 | 6.5 | 5.719 | 0 | 244 | 0 | 0.001 | | Insula R, BA 13, Insula |
| + | L | -1.5 | -77.5 | -3.5 | 5.644 | 0 | 184 | 0 | 0.003 | | Lingual L, Lingual Gyrus |
| + | R | 6 | -72.5 | -21 | 3.978 | 0 | | | | | Vermis 6, Declive |
| + | R | 3.5 | -72.5 | -8.5 | 3.791 | 0 | | | | | Vermis 6 |
| + | L | -41.5 | 15 | 4 | 5.053 | 0 | 183 | 0 | 0.003 | | Insula L, BA 13, Insula |
| + | L | -31.5 | 27.5 | 6.5 | 4.306 | 0 | | | | | Insula L, Sub-Gyral |
| + | R | 31 | -5 | 54 | 4.674 | 0 | 167 | 0.001 | 0.005 | | Frontal Mid 2 R, BA 6, Middle Frontal Gyrus |
| + | L | -31.5 | -5 | 49 | 4.523 | 0 | 205 | 0 | 0.002 | | Precentral L, Precentral Gyrus |
| + | L | -41.5 | -5 | 44 | 3.936 | 0 | | | | | Precentral L, BA 6, Precentral Gyrus |
| + | R | 36 | 37.5 | 29 | 4.375 | 0 | 110 | 0.004 | 0.029 | | Frontal Mid 2 R, Middle Frontal Gyrus |
| + | R | 36 | -57.5 | -21 | 4.017 | 0 | 57 | 0.028 | 0.187 | | Fusiform R, Declive |

*Continued on next page*

Table S20 – *Continued from previous page*

| Direction | Hemisphere | MNI Coordinates | | | Peak | | Cluster | | | Peak (SVC) | Region |
|---|---|---|---|---|---|---|---|---|---|---|---|
| | | x | y | z | Z | p(Z) *unc.* | k | p(k) *unc.* | p(k) *FWE* | p(Z) *FWE* | |
| + | L | -29 | -60 | 51.5 | 4.013 | 0 | 93 | 0.007 | 0.051 | | Parietal Sup L, BA 7, Superior Parietal Lobule |
| + | R | 31 | -52.5 | -6 | 3.847 | 0 | 36 | 0.072 | 0.414 | | Fusiform R, Parahippocampal Gyrus |
| + | L | -29 | -55 | -11 | 3.757 | 0 | 152 | 0.001 | 0.008 | | Fusiform L, Declive |
| + | L | -39 | -65 | -16 | 3.755 | 0 | | | | | Fusiform L, Declive |
| + | L | -36.5 | -57.5 | -23.5 | 3.414 | 0 | | | | | Cerebelum 6 L, Culmen |
| + | L | -29 | -77.5 | 21.5 | 3.566 | 0 | 32 | 0.087 | 0.479 | | Occipital Mid L, Sub-Gyral |
| + | R | 26 | -52.5 | 54 | 3.479 | 0 | 17 | 0.202 | 0.778 | | Parietal Inf R, Sub-Gyral |
| + | L | -4 | -27.5 | -6 | 3.401 | 0 | 7 | 0.413 | 0.954 | | |
| + | R | 1 | -17.5 | -11 | 3.291 | 0 | 6 | 0.45 | 0.965 | | |
| + | R | 21 | -55 | 11.5 | 3.29 | 0.001 | 11 | 0.302 | 0.895 | | Calcarine R, Posterior Cingulate |
| + | R | 33.5 | -72.5 | 24 | 3.23 | 0.001 | 4 | 0.544 | 0.983 | | Occipital Mid R, Sub-Gyral |

*Continued on next page*

Table S20 – *Continued from previous page*

| Direction | Hemisphere | MNI Coordinates | | | Peak | | Cluster | | | Peak (SVC) | Region |
|---|---|---|---|---|---|---|---|---|---|---|---|
| | | x | y | z | Z | p(Z) *unc.* | k | p(k) *unc.* | p(k) *FWE* | p(Z) *FWE* | |
| + | L | -36.5 | 45 | 19 | 3.117 | 0.001 | 1 | 0.785 | 0.997 | | Frontal Mid 2 L, Middle Frontal Gyrus |
| - | R | 43.5 | -70 | 41.5 | 4.218 | 0 | 73 | 0.015 | 0.103 | | Angular R, BA 19, Precuneus |
| - | R | 58.5 | -52.5 | -1 | 3.887 | 0 | 43 | 0.051 | 0.318 | | Temporal Mid R, Middle Temporal Gyrus |
| - | R | 66 | -35 | 4 | 3.511 | 0 | 7 | 0.413 | 0.954 | | Temporal Mid R, BA 22, Middle Temporal Gyrus |

**Table S21** | Face bias during encoding (global 1 face > global 1 scene at sample).

| Direction | Hemisphere | MNI Coordinates | | | Peak | | Cluster | | | Peak (SVC) | Region |
|---|---|---|---|---|---|---|---|---|---|---|---|
| | | x | y | z | Z | p(Z) *unc.* | k | p(k) *unc.* | p(k) *FWE* | p(Z) *FWE* | |
| + | R | 41 | -57.5 | -18.5 | Inf | 0 | 156 | 0.001 | 0.007 | | Fusiform R, Declive |
| + | R | 41 | -80 | -8.5 | 6.334 | 0 | | | | | Occipital Inf R, BA 18, Middle Occipital Gyrus |
| + | L | -41.5 | -85 | -11 | 5.997 | 0 | 82 | 0.01 | 0.075 | | Occipital Inf L, BA 18, Inferior Occipital Gyrus |

*Continued on next page*

Table S21 – *Continued from previous page*

| Direction | Hemisphere | MNI Coordinates | | | Peak | | Cluster | | | Peak (SVC) | Region |
|---|---|---|---|---|---|---|---|---|---|---|---|
| | | x | y | z | Z | p(Z) *unc.* | k | p(k) *unc.* | p(k) *FWE* | p(Z) *FWE* | |
| + | L | -41.5 | -72.5 | -16 | 4.797 | 0 | | | | | Fusiform L, Declive |
| + | L | -41.5 | -60 | -16 | 4.356 | 0 | | | | | Fusiform L |
| + | R | 21 | -5 | -16 | 5.133 | 0 | 34 | 0.079 | 0.445 | | Amygdala R, Amygdala, Parahippocampal Gyrus |
| + | R | 51 | -60 | 14 | 4.295 | 0 | 151 | 0.001 | 0.008 | | Temporal Mid R, Middle Temporal Gyrus |
| + | R | 48.5 | -52.5 | 21.5 | 3.582 | 0 | | | | | Temporal Sup R, Supramarginal Gyrus |
| + | L | -36.5 | 2.5 | -16 | 3.63 | 0 | 22 | 0.15 | 0.672 | | Superior Temporal Gyrus |
| + | L | -34 | 10 | -18.5 | 3.489 | 0 | | | | | Inferior Frontal Gyrus |
| + | L | -41.5 | 27.5 | -3.5 | 3.179 | 0.001 | 4 | 0.544 | 0.983 | | Frontal Inf Orb 2 L, Inferior Frontal Gyrus |
| + | R | 13.5 | -15 | 21.5 | 3.174 | 0.001 | 1 | 0.785 | 0.997 | | Lateral Ventricle |
| + | L | -24 | -5 | -16 | 3.151 | 0.001 | 1 | 0.785 | 0.997 | | Amygdala L, Amygdala, Parahippocampal Gyrus |
| + | L | -6.5 | -67.5 | -3.5 | 3.135 | 0.001 | 2 | 0.681 | 0.994 | | Lingual L |

*Continued on next page*

Table S21 – *Continued from previous page*

| Direction | Hemisphere | MNI Coordinates | | | Peak | | Cluster | | | Peak (SVC) | Region |
|---|---|---|---|---|---|---|---|---|---|---|---|
| | | x | y | z | Z | p(Z) *unc.* | k | p(k) *unc.* | p(k) *FWE* | p(Z) *FWE* | |
| + | R | 48.5 | 20 | 24 | 3.122 | 0.001 | 1 | 0.785 | 0.997 | | Frontal Inf Tri R, BA 46, Middle Frontal Gyrus |
| - | L | -24 | -45 | -11 | Inf | 0 | 2252 | 0 | 0 | | Fusiform L, Culmen |
| - | L | -29 | -52.5 | -8.5 | Inf | 0 | | | | | Fusiform L, Fusiform Gyrus |
| - | L | -34 | -85 | 21.5 | Inf | 0 | | | | | Occipital Mid L, Superior Occipital Gyrus |
| - | R | 28.5 | -42.5 | -11 | Inf | 0 | 840 | 0 | 0 | | Fusiform R, BA 37, Fusiform Gyrus |
| - | R | 38.5 | -82.5 | 26.5 | 7.358 | 0 | 1091 | 0 | 0 | | Occipital Mid R,Superior Occipital Gyrus |
| - | R | 18.5 | -55 | 14 | 7.348 | 0 | | | | | Precuneus R, Posterior Cingulate |
| - | R | 41 | -82.5 | 16.5 | 7.154 | 0 | | | | | Occipital Mid R, Middle Occipital Gyrus |
| - | L | -1.5 | -52.5 | -38.5 | 3.123 | 0.001 | 1 | 0.785 | 0.997 | | Cerebelum 9 L, Cerebellar Tonsil |

**Table S22** | Face bias during retrieval (global 1 face > global 1 scene at retro-cue).

| Direction | Hemisphere | MNI Coordinates | | | Peak | | Cluster | | | Peak (SVC) | Region |
|---|---|---|---|---|---|---|---|---|---|---|---|
| | | x | y | z | Z | p(Z) *unc.* | k | p(k) *unc.* | p(k) *FWE* | p(Z) *FWE* | |
| + | R | 23.5 | -80 | -11 | 3.995 | 0 | 123 | 0.002 | 0.017 | | Fusiform R, Fusiform Gyrus |
| + | L | -29 | -75 | -11 | 3.683 | 0 | 35 | 0.072 | 0.422 | | Fusiform L, Fusiform Gyrus |
| + | R | 46 | 22.5 | 21.5 | 3.475 | 0 | 31 | 0.088 | 0.489 | | Frontal Inf Tri R, Middle Frontal Gyrus |
| + | L | -24 | -90 | -11 | 3.444 | 0 | 11 | 0.296 | 0.896 | | Occipital Inf L, Fusiform Gyrus |
| + | L | -26.5 | 32.5 | 4 | 3.12 | 0.001 | 1 | 0.782 | 0.997 | | Insula L, Sub-Gyral |
| - | R | 6 | -35 | -8.5 | 3.3 | 0 | 1 | 0.782 | 0.997 | | Vermis 3 |
| - | R | 41 | -35 | 11.5 | 3.138 | 0.001 | 1 | 0.782 | 0.997 | | Temporal Sup R, BA 41, Superior Temporal Gyrus |

#### Questionnaires

**I. Phone screening inclusion criteria.** The yes/no questions have been administered during the phone screening.

1. Have you suffered from a **chronic illness** in the last 12 months and are under medical surveillance or treatment?

2. Have you ever been diagnosed with a **psychiatric disorder*** or **neurological disorder*** and have you been under medical surveillance or treatment for this?
   (*e.g., depression, anxiety disorder, schizophrenia, anorexia, epilepsy, or Parkinson's disease)

3. Have you ever been diagnosed with **glaucoma or increased pressure in your eye** and have you been under medical surveillance or treatment for this?

4. Have you ever been diagnosed with an **endocrine, hormonal disease or metabolic disease*** and are you under medical surveillance or treatment for this?
   (*e.g. Cushing's syndrome, Addison's disease, or thyroid disease; or Diabetes Mellitus)

5. Have you ever been diagnosed with an **obstructive pulmonary disease*** and have you been under medical surveillance or treatment for this?
   (*e.g., asthma or chronic bronchitis)

6. Have you ever been diagnosed with a **heart disease, such as an irregular heartbeat, or with anaemia, hyperthyroidism** (increased activity of the thyroid), **kidney problems** or **high blood pressure**, and have you been under medical surveillance or treatment for this?

7. Do you currently suffer from an acute infection (fever can be a symptom)?

8. Do you regularly suffer from vertigo?

9. Have you an allergy, such as **lactose intolerance or any allergies** (such as eczema, or hay fever)?

10. Do you have **poor vision** and can this not be compensated by glasses or contact lenses?

11. Do you use **medication***?
    (*other than birth control pill, homeopathy, herbal extracts or supplements, such as vitamins)

12. Do you have problems with **not smoking, not drinking alcohol and not using drugs** for 24 hours before each test day?

13. Do you have family members who suffer from **heart problems**, mainly an irregular heartbeat, for which they are treated?

    Father/mother, brothers/sisters

    More than one grandfather, grandmother, uncle or aunt (not by marriage)

14. Do you have family members who suffer from schizophrenia or manic depression?

    Father/mother, brothers/sisters

15. Do you practice top-level sport?

16. Do you suffer from claustrophobia?

17. Are you familiar with **fainting*** in certain situations?
    (*e.g., venapunction or standing for a long time)?

18. Do you have any **metal objects*** in your body?
    (*exception: dental fillings or crowns)

19. Do you have problems with lying still for ~1.5 hours?

20. Do you have problems with swallowing large pills?

**II. Medical screening questionnaire.** This questionnaire was administered during the intake session.

Subject number: .................

Following are some questions about your education, occupation, health and medical background. Most of the questions can be answered yes or no (circle what applies). Some questions you cannot answer yes or no alone. In such cases, extra space has been left blank.

#### II.a. Demographic variables

1. What is your highest level of education? .........................

2. What is your current profession? student / working / other

   If other, please specify .........................

3. What is your nationality? . . . . . . . . . . . . . . . . . . . . . . . .

4. What is your gender? male / female / other

5. Are you right or left handed? right / left / both

   If both, please specify . . . . . . . . . . . . . . . . . . . . . . . .

6. What is your current living situation (e.g., in a student house, with family etc.) alone / student house or with friends / with family / with partner / other

7. What is your age at the start of this experiment (today)? . . . . . . . . . . . . . . . . . . . . . . . .

8. On the study day, you will receive a lunch provided by us. Do you have any dietary restrictions we need to adhere to? . . . . . . . . . . . . . . . . . . . . . . . .

##### II.b. General questions

1. What is your weight (kg)? . . . . . . . . . . . . . . . . . . . . . . . .

   What is your height (cm)? . . . . . . . . . . . . . . . . . . . . . . . .

   **BMI***: **(*to be filled in by the researcher)** . . . . . . . . . . . . . . . . . . . . . . . .

2. 2. Do you do daily intensive physical training, such as topsport? yes / no

   If other, please specify . . . . . . . . . . . . . . . . . . . . . . . .

##### II.c. Medical questions

1. Do you have hearing problems? yes / no

   If yes, please specify . . . . . . . . . . . . . . . . . . . . . . . .

2. Do you have vision problems? yes / no

   If yes, please specify . . . . . . . . . . . . . . . . . . . . . . . .

   If you have corrected vision, please specify:

| | | |
|---|---|---|
| | - What is the prescription for your left eye: | + / - ................ |
| | - What is the prescription for your right eye: | + / - ................ |
| | - Do you wear glasses? | yes / no |
| | - Do you wear contact lenses? | yes / no |
| | - Is your vision good with lenses/glasses? | yes / no |
| 3. | Are you color blind? | yes / no |
| 4. | Do you have reading problems? | yes / no |
| | If yes, please specify | ........................ |
| 5. | Do you stutter? | yes / no |
| | If yes, to which extent? | always / regularly / occasionally |
| 6. | Are there any objects in/around your body, except for dental fillings/crowns, such as: | |
| | - Metal plates or screws? | yes / no |
| | - Metal fragments or splinters? | yes / no |
| | - Active implants (e.g., pacemaker, cochlear implant, insulin pump, or neurostimulator)? | yes / no |
| | - Other electronic devices? | yes / no |
| | - Unremovable piercings or jewellery? | yes / no |
| | - Prosthetics? | yes / no |
| | - Hearing aids? | yes / no |
| | - Braces? | yes / no |
| | - Mechanical contraception (e.g., IUD)? | yes / no |
| | - Medical plaster (e.g., nicotine)? | yes / no |
| | - Dental wire behind your teeth? | yes / no |
| | - Tattoos? | yes / no |
| 7. | Do you suffer from claustrophobia? | yes / no |
| 8. | Do you regularly drink alcoholic beverages? | yes / no |

| | | |
|---|---|---|
| | If yes, do you drink >20 glasses per week? | yes / no |
| | If yes, do you drink >3 glasses per day? | yes / no |
| | Do you experience mental or physical discomfort if you do not drink alcohol for 24 hours? | yes / no |
| 9. | Do you use recreational drugs weekly? | yes / no |
| | Do you experience mental or physical discomfort if you do not use drugs for three days? | yes / no |
| 10. | Do you use cannabis weekly? | yes / no |
| 11. | Do you smoke or have you smoked in the past? | yes / no |
| | If yes, do you currently smoke more than one package of cigarettes per week? | yes / no |
| | Do you experience mental or physical discomfort if you do not smoke for three days? | yes / no |
| 12. | Do you suffer or have you suffered from health problems or an illness (e.g., Cushing's syndrome, elevated thyroid production, diabetes mellitus, or a neurological disorder)? | yes / no |
| | If yes, please specify | ........................ |
| | If yes, have you been treated for this? | yes / no |
| 13. | Have you ever had surgery? | yes / no |
| | If yes, please specify | ........................ |
| 14. | Have you ever had brain surgery or head trauma? | yes / no |
| 15. | Do you suffer from skin allergies or a severe skin disease? | yes / no |
| 16. | Have you recently suffered from or have you been treated recently for an infection of the tissues that surround and support the teeth (periodontitis)? | yes / no |
| 17. | Do you suffer from skin allergies or a severe skin disease? | yes / no |

18. Concerning the lungs:

- Have you suffered from chronic bronchitis? yes / no
- Have you suffered from wheezing in the last 12 months? yes / no
- If yes, were you also short of breath? yes / no
- Do you sometimes cough up mucus when you do not have a cold? yes / no
- Have you most of the time had to cough immediately after getting up in the morning for the last 12 months? yes / no

19. Do you suffer from allergies (e.g., hay fever)? yes / no

    If yes, are you currently experiencing symptoms from this allergy? yes / no

    If yes, do you regularly take medication for this allergy? yes / no

20. Do you currently suffer from an acute infection (with fever)? yes / no

21. Do you suffer or have you suffered from a cardiovascular disease, high blood pressure, or embolism? yes / no

22. Do you suffer from vascular cramps and/or discoloration of the fingers and/or toes (e.g., Raynaud's)? yes / no

    - Do you suffer from numb fingers or toes in response to cold weather or emotional stress? yes / no

23. Are you familiar with fainting in certain situations (such as venapunction or standing for a long time)? yes / no

24. Do you often suffer from vertigo (dizziness)? yes / no

25. Do you have a current diagnosis or are being treated for:

- Major depressive disorder yes / no
- Dysthymia yes / no
- Suicidality yes / no
- (Hypo)mania yes / no
- Panic disorder yes / no
- Agoraphobia yes / no
- Social phobia yes / no

- Obsessive-compulsive disorder (OCD) yes / no
- Post-traumatic stress disorder (PTSD) yes / no
- Alcohol abuse or dependence yes / no
- Psychoactive substance abuse or dependence yes / no
- Psychotic disorders yes / no
- Eating disorders yes / no
- General anxiety disorder yes / no
- ADHD yes / no

If any are answered yes, please elaborate . . . . . . . . . . . . . . . . . . . . . . . .

26. Do you suffer from a neurological disease (e.g., Parkinson's, epilepsy)? yes / no

    If yes, please specify . . . . . . . . . . . . . . . . . . . . . . . .

27. Are you familiar with or have you recently been treated for anemia? yes / no
28. Do you suffer or have you suffered from a blood disease (e.g., porphyria)? yes / no
29. Do you have or have you had problems with urinating or emptying your bladder? yes / no
30. Do you suffer or have you suffered from kidney- or adrenal gland problems? yes / no
31. (If gender is not female:) Do you have prostate hyperplasia or hypertrophia? yes / no
32. Do you have problems with your bowel movements? yes / no
33. Do you have or have you had problems with your intestines (e.g., ileus, stenosis)? yes / no
34. Do you currently suffer from gastric and duodenal ulcers? yes / no
35. Do you suffer from increased eyeball pressure (glaucoma)? yes / no
36. Do you use medication (other than homeopathy, herbal extracts, or supplements such as vitamins)? yes / no

Are you hypersensitive to certain medications, including beta blockers? yes / no

If yes, please specify .........................

37. Have you recently used the following medication? yes / no

    - Sedative drugs
    - Corticosteroids (tablets, creams, puffs, nasal sprays)
    - Antihistamines
    - Medication for skin conditions (such as eczema)
    - Medication for asthma
    - Medication for hay fever

    If yes, please specify .........................

38. (If gender is not male:) Do you use birth control (e.g., condoms, the pill)? yes / no

    If yes, please specify .........................

39. Please write down the name and address of your general practitioner .........................

**III. Hand dominance questionnaire.** This questionnaire was administered during the intake session.

For the activities below, please indicate whether you prefer to perform them with a certain hand (left or right)? If you do not have a preference, please fill in both boxes. If you are not familiar with a particular activity, leave both boxes blank.

| | **Left** | **Right** |
|---|---|---|
| 29. Writing | ☐ | ☐ |
| 30. Drawing and painting | ☐ | ☐ |
| 31. Throwing a ball | ☐ | ☐ |
| 32. Cutting with scissors | ☐ | ☐ |
| 33. Eating with a spoon | ☐ | ☐ |

| | | |
|---|---|---|
| 34. Combing your hair | ☐ | ☐ |
| 35. Brushing your teeth | ☐ | ☐ |
| 36. Using a hammer | ☐ | ☐ |
| 37. Holding a tennis racket | ☐ | ☐ |
| 38. Turning a page | ☐ | ☐ |
| 39. Striking a match | ☐ | ☐ |
| 40. Opening a box (lid) | ☐ | ☐ |
| 41. Dealing cards | ☐ | ☐ |

**IV. Inclusion/exclusion checklist.** After the screening session and prior to inclusion, the inclusion/exclusion criteria were verified using the checklist below.

| **IV.a. Inclusion criteria:** | **Yes** | **No** |
|---|---|---|
| 1. Has the participant given written informed consent? | ☐ | ☐ |
| 2. Is the participant at least 18 years, but not older than 45 years? | ☐ | ☐ |
| 3. Is the participant right-handed? | ☐ | ☐ |
| 4. Does the participant have metal objects in or around the body? | ☐ | ☐ |
| Is the participant claustrophobic? | ☐ | ☐ |
| 5. Does the participant have an abnormal hearing or uncorrected vision? | ☐ | ☐ |
| 6. Does the participant use:<br>- More than 3 alcoholic beverages daily?<br>- Psychotropic medication or recreational drugs weekly?<br>- Cannabis weekly or more?<br>- More than a package of cigarettes weekly? | ☐ | ☐ |
| 7. Does the participant report not being able to cease use of: | ☐ | ☐ |

- Psychotropic medication or recreational drugs? (over a period of 72 hours prior to testing)
- Alcohol? (over a period of 24 hours prior to testing)
- Smoking? (over a period of 24 hours prior to testing)

8. Vital signs:

| Vital Sign | Measured value | Normal value | Markedly abnormal value |
|---|---|---|---|
| BMI | | 18.5–30 | |
| Syst. BP | | 95–180 mm Hg | |
| Diast. BP | | 50–95 mm Hg | |
| Heart rate | | 45–120 bpm | |
| ECG | | Normal ECG | |

Does the participant meet all the **vital signs** criteria? ☐ ☐

1129

| **IV.b. Exclusion criteria:** | **Yes** | **No** |
|---|---|---|
| 9. Does the participant have a **history** of a clinically relevant: | ☐ | ☐ |

- Psychiatric disease
- Neurological disease
- Endocrine / metabolic disease
- Obstructive respiratory disease
- Hepatic, cardiac, or renal disease
- Heart-related disease
- Cerebrovascular, metabolic, or pulmonary disease
- Epilepsy
- Drug dependence (e.g., opiates, LSD, (meth)amphetamine, cocaine, solvents, barbiturates)
- Alcohol dependence
- Raynaud's syndrome
- Glaucoma

- Diabetes

Or is **currently** being treated for a clinically relevant: ☐ ☐

- Chronic disease
- Hypo- or hypertension
- Renal failure
- Hyperthyroidism
- Acute inflammatory disease
- Peptic or duodenal ulcers
- Glaucoma

10. In the week prior to the start of the study did the participant use: ☐ ☐

- Corticosteroids
- MAO inhibitors
- Antidepressants
- Antipsychotics
- Anaesthetics

11. Is the participant oversensitive to methylphenidate, sulpiride, carbidopa or entacapone? ☐ ☐

12. Does the participant have a history of: ☐ ☐

- prescribed medications within the last month (with exception of regular use of contraceptive mediation)
- 'over the counter' medication within the last 2 months? (with exception of occasional use of paracetamol, acetylsalicylic acid, and ibuprofen)
- regular use of corticosteroids?

13. Does the participant have (a history of) frequent autonomic failure? ☐ ☐

14. Does the participant have abnormal hearing or abnormal corrected vision? ☐ ☐

15. Does the participant have epilepsy? ☐ ☐

16. Does the participant have a family history of sudden death or ventricular arrhythmia? ☐ ☐

17. Does the participant have a first-order family history of schizophrenia, bipolar disorder, or major depressive disorder? ☐ ☐

18. Does the participant have an irregular sleep/wake rhythm? ☐ ☐

19. Does the participant carry out daily intense physical exercise? ☐ ☐

20. Does the participant currently have periodontitis? ☐ ☐

21. Is the participant pregnant or breastfeeding? ☐ ☐

22. Is the participant using appropriate contraception? ☐ ☐

**Final check:** Does the participant meet **all** the above listed inclusion criteria and none of the above listed exclusion criteria?

Please accept the participant by entering a specific participant number:

. . . . . . . . . . . . . . . . . . . . . .

#### V. Medical screening before drug intake.

1. How are you feeling? How is your general health? . . . . . . . . . . . . . . . . . . . . . . .

2. Have you used cannabis during the last two weeks? yes / no

3. Have you:

   - used psychotropic medication, or recreational drugs in the last 72 hours? yes / no
   - used alcohol within the last 24 hours? yes / no
   - smoked during the last 24 hours? yes / no

4. Have you been ill during the last weeks, or started a treatment with a doctor or specialist? yes / no

5. Have you started using any medication since the screening session? (e.g., for hay fever, sedative medication, corticosteroids (with the exception of occasional use of paracetamol, aspirin, ibuprofen and regular use of contraception) yes / no

6. In the last weeks, did you use corticosteroids, MAO inhibitors, antidepressants, antipsychotics or anesthetics? yes / no

7. Do you currently have an acute infection (with fever)? yes / no

8. Do you currently suffer from problems with your stomach or intestines (e.g., a gastric / duodenal ulcer)? yes / no

9. (If not male:) Are you pregnant or do you expect to become pregnant in the coming few months? yes / no

10. (If not male:) Are you using contraception? yes / no

    If yes, please specify ........................
